# *In situ* estimation of genetic variation of functional and ecological traits in *Quercus petraea* and *Q.robur*

**DOI:** 10.1101/501387

**Authors:** Hermine Alexandre, Laura Truffaut, Alexis Ducousso, Jean-Marc Louvet, Gérard Nepveu, José M. Torres-Ruiz, Frédéric Lagane, Cyril Firmat, Brigitte Musch, Sylvain Delzon, Antoine Kremer

## Abstract

**Background:** Predicting the evolutionary potential of natural tree populations requires the estimation of heritability and genetic correlations among traits on which selection acts, as differences in evolutionary success between species may rely on differences for these genetic parameters. *In situ* estimates are expected to be more accurate than measures done under controlled conditions which do not reflect the natural environmental variance.

**Aims:** The aim of the current study was to estimate three genetic parameters (i.e. heritability, evolvability and genetic correlations) in a natural mixed oak stand composed of *Quercus petraea* and *Quercus robur* about 100 years old, for 58 traits of ecological and functional relevance (growth, reproduction, phenology, physiology, resilience, structure, morphology and defence).

**Methods:** First we estimated genetic parameters directly *in situ* using realized genomic relatedness of adult trees and parentage relationships over two generations to estimate the traits additive variance. Secondly, we benefited from existing *ex situ* experiments (progeny tests and conservation collection) installed with the same populations, thus allowing comparisons of *in situ* heritability estimates with more traditional methods.

**Results:** Heritability and evolvability estimates obtained with different methods varied substantially and showed large confidence intervals, however we found that *in situ* were less precise than *ex situ* estimates, and assessments over two generations (with deeper relatedness) improved estimates of heritability while large sampling sizes are needed for accurate estimations. At the biological level, heritability values varied moderately across different ecological and functional categories of traits, and genetic correlations among traits were conserved over the two species.

**Conclusion:** We identified limits for using realized genomic relatedness in natural stands to estimate the genetic variance, given the overall low variance of genetic relatedness and the rather low sampling sizes of currently used long term genetic plots in forestry. These limits can be overcome if larger sample sizes are considered, or if the approach is extended over the next generation.

## 1 Introduction

Natural populations live in constantly changing environments that trigger natural selection and contribute to genetic changes of phenotypic traits. Evolutionary changes of traits are constrained by the level of genetic variation and by genetic correlations with other traits. Heritability is a standardized measure of genetic variation, and is a pivotal genetic parameter used in theoretical and practical oriented genetic investigations in domesticated and wild organisms (Visscher, 2008; Lynch and Walsh, 1998). The estimation of heritability builds on the comparison of phenotypic resemblance of relatives with their genetic relatedness (Lynch and Walsh, 1998). In most domesticated species submitted to artificial selection, heritability was assessed in families with known pedigree generated by controlled crosses. In trees, because the long lifetime of individuals prevents from making controlled crosses, a traditional approach is to collect progenies from open pollination on mother trees and estimate genetic variance in a common garden with a half-sib family design. This procedure has been proven to bias additive variance estimates due to a bad estimation of relatedness among individuals (Gauzere et al., 2013). In populations undergoing natural selection, where pedigree relationships are unknown, alternative approaches to estimate relatedness among individuals were implemented, making use of genetic markers (Mousseau et al., 1998; Ritland, 2000). However these attempts mainly based on a few dozen of microsatellites were disappointing, as variation of relatedness *in situ* was limited and the sampling variance of relatedness between individuals was still too high to provide reliable estimates of heritability (Coltman, 2005). More recently next generation sequencing allowed to call thousands of markers thus improving the estimation of genomic relatedness (Robinson et al., 2013; Vinkhuyzen et al., 2013). As shown in recent reports stemming from wild or domesticated organisms, a few thousand markers are sufficient to access the realized relatedness among individuals (Bérénos et al., 2014; Stanton-geddes et al., 2013). Thus, at least theoretically, the estimation of the additive variance of traits in natural population of trees at unique time points became possible. We aimed at implementing this approach in long-lived forest trees as oaks.

However, the tentative use of heritability for predicting evolutionary shifts in wild populations undergoing natural or human mediated selection pressures is constrained by other theoretical and biological limitations (Charmantier et al., 2014; Kruuk, 2004). Spatial genetic structure may build in naturally regenerated forest stands and add up to the ecological structure (Bontemps et al., 2016; Kruuk and Hadfield, 2007). Ecological and spatial genetic structures may lead to potential common environment effects, and non independence of genetic and environmental effects may thus increase biases of the genetic variance. This issue can be tackled by the ecological description of the study plot with a recording of the spatial distribution of the trees. Hence, ecological and spatial source of variation can be explicitly introduced in the statistical model aiming at estimating the genetic variance. Finally, the contribution of environmental variance to heritability, which is likely to be inflated under natural settings, has challenged its use in evolutionary studies and lead to the use of evolvability, an alternative parameter to account for the evolutionary potential of populations (Hansen et al., 2011). On the other hand, besides limitations there are also assets for assessing heritability *in situ* in forest trees. As trees are long-lived species, two or even more generations of trees may actually coexist in the same forest stand over very short time scales, thus allowing to access phenotypic values over multiple generations while concomitantly increasing the variance of relatedness. Perennial plants such as trees also allow to assess longitudinal traits over multiple years (as growth or bud phenology) which enables to take account of temporal environmental variance.

Building on this background knowledge and experience, we attempted to assess heritability in a 100 year old oak stand in a France North West forest comprising two common European white oak species (*Quercus petraea*, Sessile oak and *Q. robur*, Pedunculate oak) growing under natural selection pressures and human driven management. Previous studies conducted on this mixed stand showed that *Q. petraea* and *Q. robur* occur in different geographic and ecological positions over the stand and that over two generations *Q. petraea* gained ground over *Q. robur*. Thus, different genetic and/or plastic responses are expected for these two species based on their ecology and distribution. Accessing their genetic variation would allow to anticipate their respective ability to respond to ongoing selection pressures. In an earlier companion paper, we implemented a genomic capture approach to call thousands of SNPs and evaluate their use for recovering realized relatedness (Lesur et al., 2018). Estimates of realized relatedness were validated by comparing their values with known pedigree relationships. Another previous study enabled to retrieve geographic position of each tree over the stand as well as the characterization of five environmental variables (Truffaut et al., 2017). Finally progeny tests and clonal collections stemming from the same stand were earlier established thus allowing to compare heritability estimates between *in situ* and *ex situ* settings. The study forest stand therefore offered an ideal design for addressing methodological and biological issues regarding genetic variation in natural populations of trees. At the methodological level, we focused our investigations on methods for assessing parameters of genetic variation *in situ*, and comparing *in situ* with *ex situ* estimates. Indeed previous studies in trees have shown that heritability estimates can vary widely depending on experimental conditions (i.e. *in situ* or in common garden Castellanos et al. (2015)). At the biological level, we explored a large spectrum of traits involved in various functions putatively contributing to the adaptive value of oak trees: growth, phenology, water metabolism, morphology, secondary metabolites composition and wood structure. Thanks to decades of research in this forest stand (Truffaut et al., 2017), phenotypes of these trees were dissected in multiple traits in order to grasp the distribution of genetic variation at a broad scale.

To sum up, the goals of the current study were thus threefold: (1) Implement and compare methods for assessing the additive genetic variance (heritability and evolvability) in oak forests. (2) Describe the distribution of heritability and evolvability across traits having different functional and adaptive values and analyze genetic correlations among traits. (3) Examine whether the distribution of heritability and evolvability exhibits notable species differences in line with their expected response to ongoing environmental changes and their current demography trajectories.

## 2 Material and Methods

### 2.1 Study population: first and second generations *in situ* (Figure 1)

The studied forest stand is a mixed oak population composed of *Q. petraea* and *Q. robur* in the Petite Charnie forest in North West of France and covers 5.19 ha (square of 230 × 226 m). Intensive investigations have been conducted during the past three decades in this even aged stand addressing spatial genetic structure, mating system, gene flow, hybridization and species dynamics over two successive generations (Bacilieri et al., 1994, 1996; Streiff et al., 1998; Truffaut et al., 2017). The present study benefited from the previous results and extends over two successive generations of the stand. Detailed background information on the study stand is provided in Truffaut et al. (2017) and Appendix 1. Generation one (G1) consists of 422 hundred-year-old trees (196 *Q. petraea* and 226 *Q. robur*) that mated naturally to produce generation 2 (G2) between 1989 and 2001. As the saplings of G2 got established, a clear cut of the parental canopy was done between 1998 and 2001. The saplings were extremely dense and a systematic sampling of 2510 saplings (one every 3 to 6 m.) was implemented to reconstruct half or full sib families for the estimation of the additive genetic variance, while admixed individuals were discarded from the analysis. A parentage analysis was therefore conducted which resulted in the selection of a subset of 370 saplings of *Q. petraea* and 390 of *Q. robur*, corresponding to 169 full sib (FS) and half sib (HS) families in *Q. petraea* and 233 FS and HS families in *Q. robur* (Appendix 1) used for the estimation of heritability and evolvability. All trees of G1 and G2 were mapped by recording their GPS coordinates, using post-processed differential corrections. As described in Truffaut et al. (2017), a floristic survey (119 plant species) conducted in 1992 in 34 plots distributed systematically according to a grid system allowed to derive indicator values of key ecological characteristics: pH, soil moisture, ratio of carbon to nitrogen (C/N) and organic matter content for each sampling plot. These variables were further downscaled to the level of each tree of G1 and G2 after mapping the distribution of the indicator values by kriging (Supplemental material Methods S2 in Truffaut et al. (2017)). Altitude was also recorded for each tree.

### 2.2 Study population: first and second generations *ex situ* (Figure 1)

Before the final cut of the G1 trees, between 1995 and 2001, scions were collected on the 298 remaining trees and grafted in an *ex situ* conservation collection located in a State Nursery of Guéméné Penfao (latitude: 47.631° N; longitude: 1.892°W). Multiple clonal copies were done for each genotype, but grafting was not always successful. Thus only 147 G1 *Q. robur* genotypes and 116 G1 *Q. petraea* genotypes with a mean number of 2.93 copies per genotype were successfully propagated by grafting. The grafts were planted in a fully randomized design. Besides the conservation collection of the G1 trees, a common garden experiment comprising open pollinated progenies of G1 trees was also set up. In the fall 1995, 28 open pollinated (OP) progenies of *Q. robur* and 23 OP progenies of *Q. petraea* were collected and raised in the State Nursery of Guéméné Penfao and transplanted in march 1998 back in the Petite Charnie State Forest near the study stand comprising the G1 trees. Two common garden experiments corresponding to the two species were planted next to each other according to an incomplete block design (Appendix 2). A parentage analysis was performed to assign parents to the offspring using genotypic arrays obtained with 12 microsatellite loci using CERVUS v.3.0.7 (Marshall et al., 1998) and published in an earlier paper (Lagache et al., 2013). The parentage analysis was conducted assuming no errors in genotyping (a strict exclusion analysis: 0.0 error rate) and a high confidence level (95%). All female parents were confirmed by the parentage analysis, and male parents were identified for 641 and 982 offspring of *Q. petraea* and *Q. robur* (Appendix 2), which constitute the sample material for estimating heritability in the *ex situ* progeny test.

**Figure 1:**
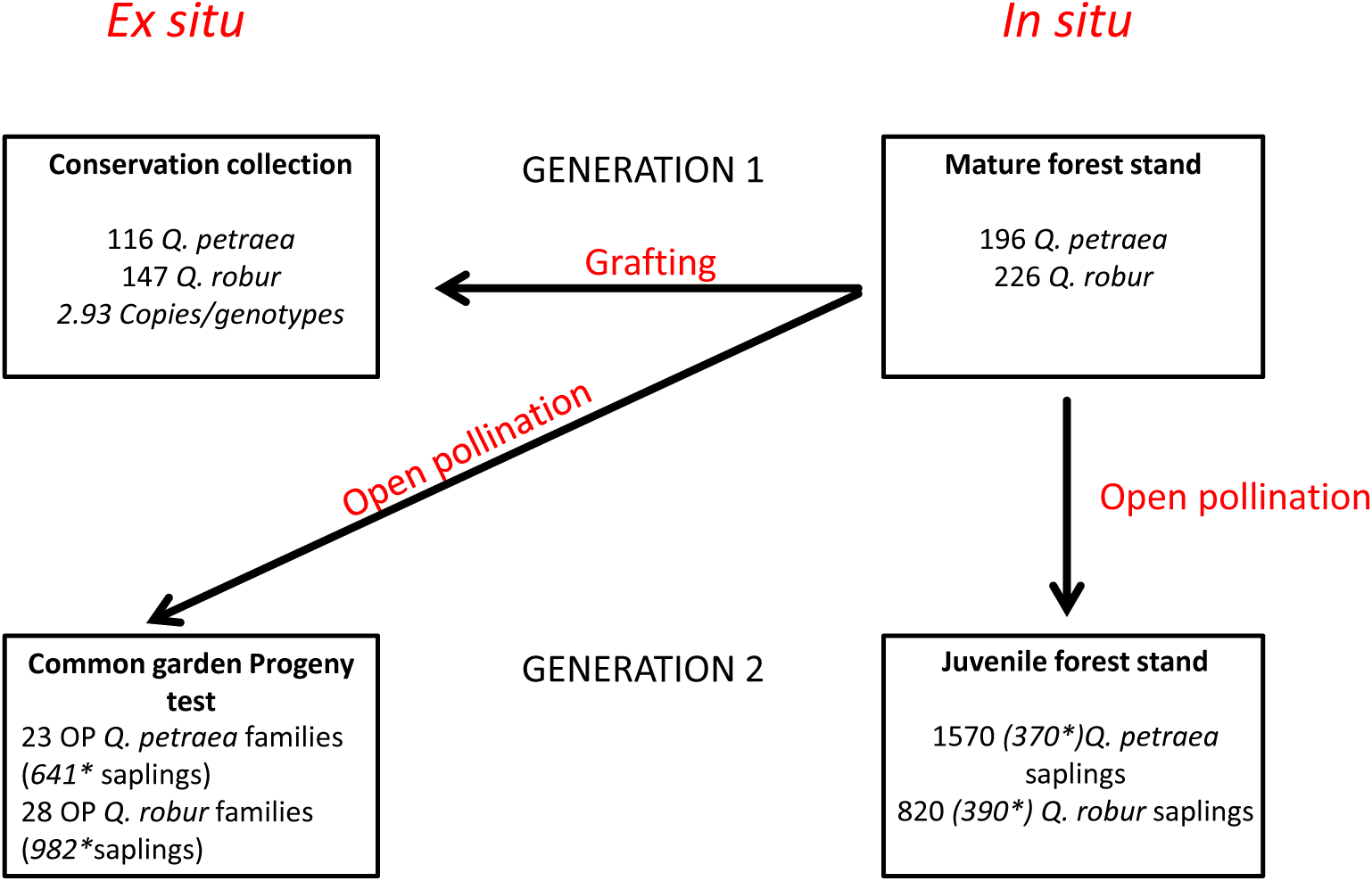
Schematic representation of the study experimental design. *: Numbers in italics correspond to number of saplings with reconstructed pedigree by parentage analysis.

**Table 1:**
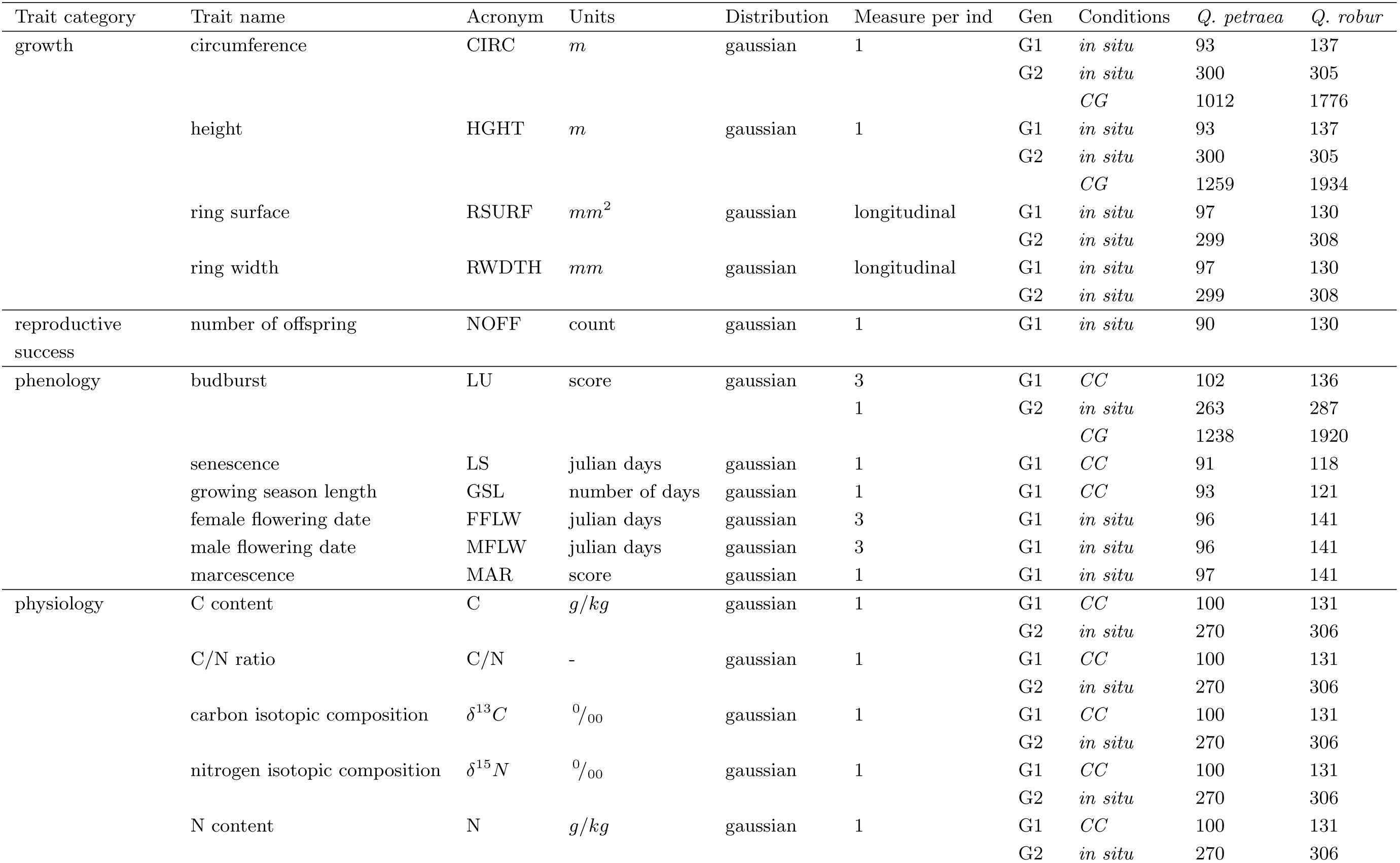
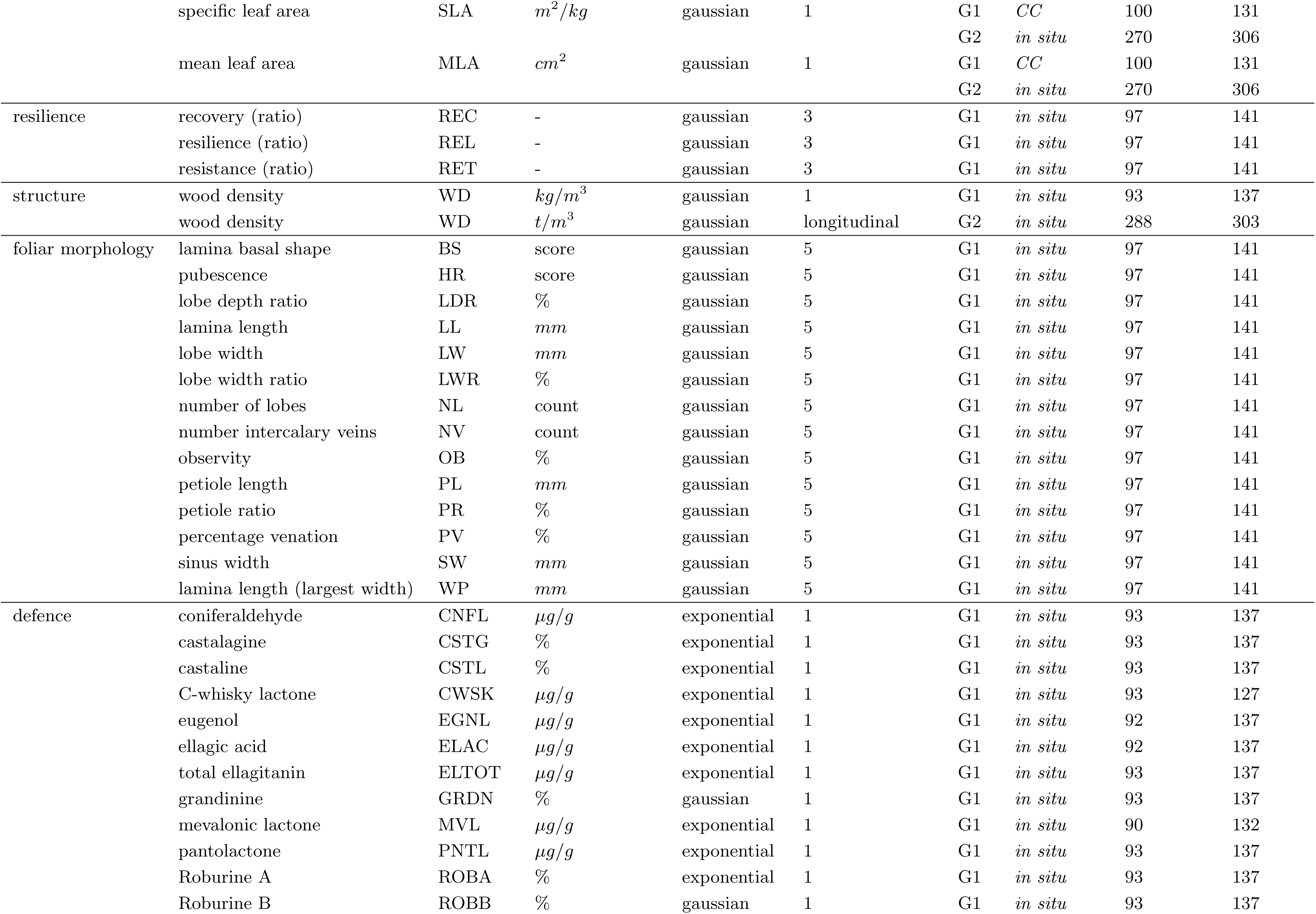
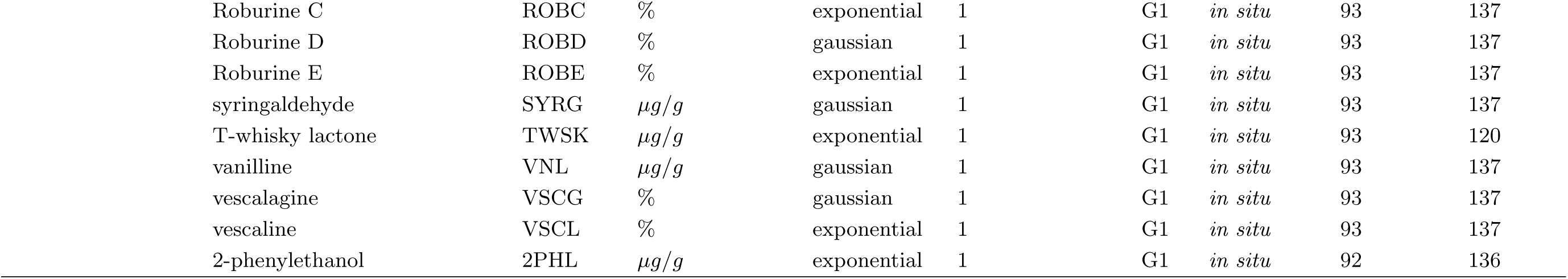
Description of each trait analysed in the study. ”Distribution” represents whether the trait followed a gaussian or exponential distribution (see analysis details in material and methods). ”Measure per ind” is the number of measure per genotype.”Gen” stands for Generation 1 or 2. ”Conditions” indicates whether the trait was measured *in situ* or *ex situ* (CC: conservation collection, CG: common garden). *Q. petraea* and *Q. robur*: the number of phenotyped individuals.

### 2.3 Phenotypic assessments in generation G1 (Table 1)

#### 2.3.1 Growth

We took advantage of the final removal cut of G1 trees to assess numerous dimensional phenotypic traits of these 100 year old trees. Cutting operations were subdivided over three years (December 1998 to March 2001) which facilitated the recording of traits. Circumference (CIRC) at breast height was recorded before the trees were felled, and the total height (HGHT) of the trees was assessed once they were felled. Two 10 cm wide sections (section 1 and 2) were collected on the main stem between 1.30 m to 1.50 m from the ground level for later assessments in the lab. Section 1 was used for measuring all yearly tree ring widths on four radius along the four cardinal directions, and for assessing wood density (see paragraph Structure - Wood density). Ring width (RWDTH) and ring surface (RSURF) was recorded for each ring in each individual. Section 2 was subsequently used for extracting and analysing secondary metabolites composition of the wood (see paragraph Defence: secondary metabolites).

#### 2.3.2 Reproductive success

Reproductive success (NOFF) of each tree of G1 was estimated by performing a parentage analysis based on 80 SNP loci, and using stringent parameters, assuming no errors in genotyping (a strict exclusion analysis: 0.0 error rate) and a high confidence level (95%) (see Truffaut et al. (2017) for further details). Reproductive success of a parent tree corresponded to the number of living offspring it produced as male or as female parent (Appendix 1).

#### 2.3.3 Phenology

Male and female flowering (MFLW and FFLW) of each single tree was monitored every three days in spring 1990, every 14 days in spring 1991 and every 7 days in spring 1992 (Bacilieri et al., 1994). Floral development was recorded in the upper part of the crown with a telescope with 25x or 40x magnification using a grading system with five classes from early (1) to late (5) flowering stage. From these observations, a flowering date for male and female flowers was computed, separately for 1990, 1991 and 1992 (date in Julian days at which flowers attain the stage 3, *i.e*. catkins releasing pollen for males and receptive flowers when the pistil exhibits a bright red colour for females). Leaf unfolding (LU) of apical buds was monitored in April 2016 in the *ex situ* grafted conservation collection by scoring the stage of development (5 classes) using the protocol of Vitasse et al. (2009). LU was used as an assessment of the phenological development of the vegetative apical bud. Two measures were used: *LU_s_* is the score at each observation date, and *LU_d_* is the date of unfolding in julian days derived from *LU_s_*. Leaf senescence (LS) was also assessed in the late season of 2016 in the *ex situ* collection. Starting on October 26th and ending on November 21st, the percentage of leaves turning yellow or falling was visually scored on each tree, and leaf senescence was considered to be completed when more than 50% of the crown turned yellow. LS is the date in julian days at which senescence was complete. We computed the growing season length (GSL) as the number of days from leaves unfolding to senescense *(GSL* = *LS* − *LU_d_*). In January 1998 and February 2000, leaf retention (MAR for marcescence) within the canopy of the trees was assessed *in situ* using a scoring method from 0 to 5.

#### 2.3.4 Physiology

Branches (about 50 cm in length) were cut in the upper crown of the grafts located in the *ex situ* conservation plantation using a pole pruner in summer 2016. Branches were wrapped after collection in sealed plastic bags to avoid desiccation and transported to the lab. Leaf area was measured on 6 to 8 leaves collected on the branches using a desktop scanner (Expression 10000 XL, Epson, Japan) and WinFolia software (Regent Instruments Inc., Quebec, Canada) and averaged to obtain mean leaf area (MLA). Leaves were then dried in an oven at 65° C up to reach constant mass. Specific Leaf Area (SLA) was then measured as the ratio between leaf area and dry mass. The same leaves were used for determining the carbon and nitrogen content (C and N g/kg, respectively) and assessing stable isotopic composition (*δ*^13^*C* and *δ*^15^*N* respectively) following the procedure described in Torres-Ruiz et al. (in prep), according to the formula:

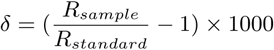

where *δ* stands for *δ*^13^*C* or *δ*^15^*N* and *R* is the ratio ^13^*C*/^12^*C* or ^15^*N*/^14^*N*.

#### 2.3.5 Resilience

Resilience components of the trees were assessed by scanning the response of tree cambial growth after a severe disturbance. The method consisted in comparing ring width before, during and after so called negative pointer years when the disturbance occurred (Lloret et al., 2011). We used the whole data set of tree rings available over 100 years to identify 7 and 10 negative pointer years in *Q. petraea* and *Q. robur*, respectively (Appendix 3). The following three resilience components were calculated. Resistance (RET): inverse of ring width reduction during the disturbance. Recovery (REC): increased ring width after the disturbance relative to the minimum ring width during the disturbance, which expresses the ability of tree growth to recover after the disturbance. Resilience (REL): ring width after recovery relative to ring width before the disturbance, which reflects the ability of trees to reach pre-disturbance growth levels (Folke et al., 2004). Resilience components were derived from tree ring analysis for three periods of the lifetime of a tree: the juvenile period before age 30, the intermediate period when trees were between 30 and 60 years old and the mature stage when trees were older than 60 years (Appendix 3).

#### 2.3.6 Structure - Wood density

Two radial wood bars (6cm thickness) extending from the cambium to the core of the tree were randomly delineated and extracted from section 1 collected on the stem (see paragraph Growth) and were saturated in water during 48 hours and later dried at 100°C during 28 hours. Wood density (WD) of G1 trees was assessed as infradensity by taking the ratio of the dry weight to the water saturated volume of the bars (Guilley, 2000).

#### 2.3.7 Foliar morphology

Leaf morphology data were extracted from a previous study (Kremer et al., 2002). The data comprise nine raw traits (LL: lamina length, PL: petiole length, LW: lobe width, SW: sinus width, WP: length of lamina at largest width, NL: number of lobes, NV: number of intercalary veins, BS: basal shape of the lamina, HR: pubescence) and five synthetic traits computed from the previous nine (OB: lamina shape obversity, PR: petiole ratio, LDR: lobe depth ratio, PV: percentage venation, LWR: lobe width ratio). Data were available for five leaves collected in the upper part of the crown within each tree sampled *in situ*.

#### 2.3.8 Defence: secondary metabolites

Wood metabolite compounds were analysed from section 2 collected on the stem of all adult trees ; to do so a 10 cm wide diametral strip was extracted from the section, and wood shaving was carried out at its two extremities (excluding sapwood). Wood shavings from about 35 to 40 rings were used for subsequent extractions and analysis of the compounds either by HPLC (for ellagitannins, Prida et al. (2006)) or GC/MS (for other volatile compounds Prida et al. (2007)). In total 21 compounds were identified. While their ecological roles have not yet been studied in oaks, tannins and volatile compounds are suspected to be involved in tree resistance to insect and pathogens, and chemical signaling (Tumlinson, 2014).

### 2.4 Phenotypic assessments in generation G2 (Table 1)

A subset of traits was also assessed on the 370 and 390 trees of *Q. petraea* and *Q. robur* sampled *in situ* within G2 or in the common garden for estimating heritability. Besides the age related differences between G1 (about 100 years old) and G2 (14 to 26 years old), there were also some differences in the procedures and protocols used for assessments in G2 in comparison to G1. Total height (HGHT) was assessed *in situ* on the standing saplings using a vertex, and in the common garden using a pole. Leaf unfolding (*LU_s_*) was monitored *in situ* on April 11–12 2017 and in common garden on 16 April 2002 and on 14 and 21 April 2011, by scoring the stage of development of the apical bud (from 0 to 5). There was not enough observation time points to transform *LU_s_* into *LU_d_*. We thus only analyzed *LU_s_* in the second generation. Wood density (WD) was measured *in situ* on the increment cores by using an x-ray image calibration procedure. Increment cores were exposed to X rays and were then scanned with a microdensitometer following the procedure by Polge and Nicholls (1972). To get one measure per individual for better comparison between generations, we used the mean of all the cores for each individuals, multiplied by 1000. We used the same cores to estimate ring width (RWDTH) and surface (RSURF). Physiology related traits were measured *in situ* following the same protocol than for G1.

A summary table of all traits assessed in the two generations and in the different experiments *(in situ*, conservation collection, common garden experiment) is provided in Table 1.

### 2.5 Estimation of the additive genetic variance using the animal model

We used the animal model to estimate the additive variance in the different experimental settings (see Figure 2a for a summary of experimental settings). There are two different models that were applied to the data, depending on the experimental design (i.e. *in situ* or *ex situ*):

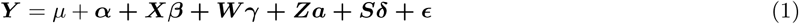

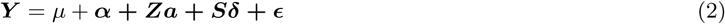

Model 1 corresponds to *in situ* estimations while model 2 was used for estimations in *ex situ* settings. ***Y*** is the vector of phenotypic traits, *μ* is the population mean, ***X***, ***W***, ***Z*** and ***S*** are incidence matrices related to each effect, *β* is the fixed competition effect, *γ* is the fixed environmental effect, *δ* is the random spatial effect either associated to a covariance structure among individuals for model 1 or independent effects in model 2, *α* is the random additive genetic effect associated with a covariance structure among individuals computed differently if relatedness information is available among all individuals (i.e. in natural population and in the *ex situ* conservation collection of G1) or if the only relatedness information available is the mother-offspring pedigree relationship (i.e. for offspring in the *ex situ* common garden), and *ε* is the residual effect. The main parameter we want to estimate with these models is the variance of the additive genetic effect *V_a_*,

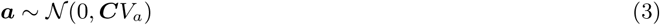

where *C* is the additive genetic relationship matrix, corresponding to the non genetic independence of individuals. From the animal model, we can estimate heritability as *V_a_/V_p_* where *V_p_* is the phenotypic variance. Usually *V_p_* is estimated as the sum of variance of all the random effects (and heritability computed this way is called 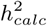). However, heritability estimates are highly dependent on the fixed effects included in the model (Wilson, 2008), and all designs do not include the same fixed effects. For heritability estimates to be comparable between traits and experimental designs, we thus decided to calculate also the observed phenotypic variance value, computed directly from phenotypes (heritability called 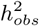). We also computed evolvability (*I_a_*) as *V_a_/x*^2^ where *x*^2^ is the squared population phenotypic mean (Hansen et al., 2011), which is an estimate less dependent on environmental variance than heritability. However, this way of computing evolvability is meaningless for traits measured on an interval scale (Hansen et al., 2011) (e.g. for julian days). Also, for isotopic compositions, mean scaled additive variances are dependent on the standard used for the determination of isotopic composition (see section 2.3.4) (Brendel, 2014). Thus evolvability was not reported for some phenology and physiology traits.

The estimation of additive genetic variance was done with a linear mixed model (LMM) enabling for a covariance structure between random effects (usually denoted as the animal model), implemented in the R package breedR under the function remlf90 (Muñoz and Sanchez, 2018). This is an estimation based on restricted maximum likelihood. We took account of spatial autocorrelation among phenotypes including a spatial random effect and environmental variables were included as fixed effects. We report here the estimated values of heritability and evolvability as well as the 95% confidence intervals (CI) obtained by bootstraping 1000 times with data simulation using a modified version of the function breedR.sample.phenotype (see breedR tutorial for details) taking quantiles 2.5% and 97.5% of the bootstrap distribution. Bootstrap estimations of confidence intervals are indeed recommended when using LMM models with restricted maximum likelihood (Schweiger et al., 2016). The fit of animal model was compared to the fit of the same model without the additive genetic effect according to their AIC values with to the formula

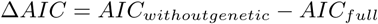

A positive Δ*AIC* reflects that including genetic additive effect in the model increases model fitting, while a negative value shows that the data are better explained without the additive effect. The same was done estimate the importance of the spatial random effect.

For traits with longitudinal or repeated measures, the genotype (or individual) effect was simply included via introducing an individual effect. Model 1 and model 2 become therefore

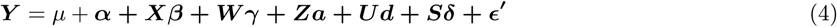

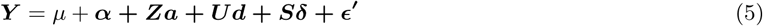

where *d* is the individual non genetic effect, *U* the corresponding incidence matrix and *ε*′ is now the within individual residual effect (due to the replicated assessments over years or over samples within a tree). For such traits the phenotypic variance to compute heritability does not include *V_ε_*′.

#### 2.5.1 Genotype as random effect

The genetic relationships structure between individuals (***C*** matrix in eq. (3)) can be either introduced as a theoretical relationship coefficient among individuals derived from pedigree information or as the effective genomic relatedness estimated with multiple genetic markers (Kruuk, 2004). One generation pedigree relationships were inferred from parentage analysis between G2 saplings and G1 trees and were reported in earlier companion publications (Lagache et al. (2013) for *ex situ* G2 saplings, and Truffaut et al. (2017) for *in situ* G2 saplings). The relatedness information extracted from pedigree, denoted as ***A*** matrix was thus useful to estimate heritability from G2 offspring. We estimated pedigree relationship for all G2 individuals with at least one parent retrieved from parentage analysis, while results for individuals with both parents known are presented in appendix. Genomic relatedness was estimated among G1 individuals in a previous study (Lesur et al., 2018) following the method by Van Raden (2008) and using multiple SNP markers derived from target sequence capture. Relatedness was computed separately for each species and using 6 different sets of SNPs selected according to different thresholds of Minimum Allele Frequencies (MAF: 0.01, 0.05, 0.1, 0.15, 0.3 and 0.4) representing 33,000 to 1,500 markers (Appendix 4). In what follows, the genomic relatedness matrix is called ***G*** matrix and was used to estimate the additive genetic variance in the G1 population, by replacing ***C*** by ***G*** in equation (3). When phenotypic information was available for two generations but genomic information is lacking for one generation (e.g. pedigree was missing for the first generation or genomic data were missing for the second), one can combine the genetic information from the first generation with the pedigree information from the second. We thus combined genomic relatedness of G1 with theoretical relatedness of G2 in a global ***H*** matrix according to the formula:

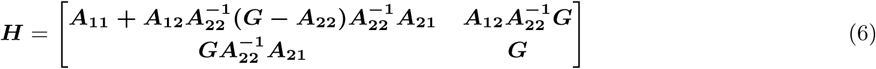

where ***A*_11_** is the part of pedigree matrix containing non genotyped individuals, ***A*_22_** is the part of the pedigree matrix containing the genotyped individuals and ***A*_12_** and ***A*_21_** are the pedigree relationships between genotyped and non genotyped individuals (Ratcliffe et al., 2017).

#### 2.5.2 Spatial random effect

In the *in situ* natural population, each individual was previously geolocalised (Truffaut et al., 2017). From decimal coordinates of each individual, we built a spatial model in breedR according to the splines model which introduces a covariance structure between neighbourhood plots. In the natural population individuals were not regularly distributed; we thus artificially separated the study plot into a 10*10 grid where each individual was assigned to a square of the grid according to its coordinates. Thus individuals occurring in the same square shared the same spatial effect. For the *ex situ* conservation collections, spatial information was available as blocks and lines, with 2 blocks each one containing 1 to 11 lines. For physiological traits measured *ex situ*, spatial information was missing so we did not include a spatial effect in the model. In the *ex situ* common garden experiment, spatial information was also available as blocks with 90 blocks. These nested effects were introduced in the model as classical random effect without a covariance structure.

#### 2.5.3 Competition as fixed effect

The uneven spatial distribution of standing trees *in situ* may have generated varying competitive interactions between trees, which in turn may have contributed to the variation of the phenotypic traits we have assessed. We accounted for the competitive interactions by computing for each tree the Hegyi competitive index (Hegyi, 1974; Contreras et al., 2011) as:

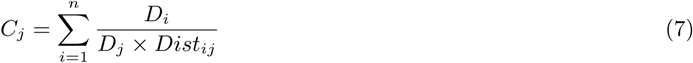

with *n* the number of trees within the neighborhood of the subject tree *j* (neighborhood of the subject tree is a circle of radius 10 meters), *D_j_* the diameter at breast height of the subject tree, *D_i_* the diameter at breast height of tree *i* standing in the neighborhood of the subject tree and *Dist_ij_* the distance between subject tree and tree *i*. ***C*** was included as a fixed effect variable.

#### 2.5.4 Environmental variables as fixed effect

In the *in situ* natural population, five environmental variables (i.e. altitude, pH, soil moisture, organic matter soil content and carbon/ nitrogen ratio) were recorded by kriging at the level of each individual (Truffaut et al., 2017). These environmental variables are highly correlated, which may cause problems in linear regressions for parameter estimation. To remove colinearity among these variables, we performed a principal component analysis (PCA) with all phenotyped individuals and used the first principal component as a fixed effect variable instead of the raw environmental variables. When no environmental variables were measured (*ex situ* populations), these variables were not included in the model.

#### 2.5.5 Taking account of non-gaussian trait distribution

The model implemented in breedR stands for traits with a normal distribution only. However several traits showed an exponential distribution (see Table 1). In this case we log-transformed the phenotypic data and performed the animal model on the transformed data. Evolvability of the trait on the original scale is equivalent to the additive genetic variance of the transformed trait (Hansen et al., 2011), and heritability of the original trait was computed with the function *QGparams* of the QGglmm R package (Villemereuil, 2018). This function is designed to estimate quantitative genetic parameters on the data scale when fitting a generalized linear mixed model. As our case is not a GLMM but a LMM fitted on log-transformed data, we used an exponential as the derivative of the inverse link function no distribution variance (see Villemereuil (2018) for details).

### 2.6 Summary of the models for estimating heritability and evolvability

Figure 2 summarizes the different models used to estimate heritability and evolvability in the different experimental settings. Models differ according to the phenotypic assessments made in the *in situ* and *ex situ* settings, but also according to the approaches used to assess genetic relatedness (***C*** matrix) among trees within and between the two generations. There are major differences among the different models. On the one hand there are models based on an unstructured population where the overall relatedness is low (M1 and M2), and others making use of family structured population based on pedigree reconstruction (M3, M4, M6, M7) where relatedness is on average higher and more variable, and finally models combining both (M5). While our main focus is to compare experimental designs to estimate additive variances *in situ* in natural populations (comparison of M1, M3, M4 and M5), we took also advantage of the existing settings to compare estimates obtained between *in situ* and *ex situ* (comparison of M3 and M7), and over two generations (comparison of M1 and M3 and comparison of M2 and M7). Finally we compared heritability and evolvability among traits and species for the first generation (M1 and M2).

**Figure 2:**
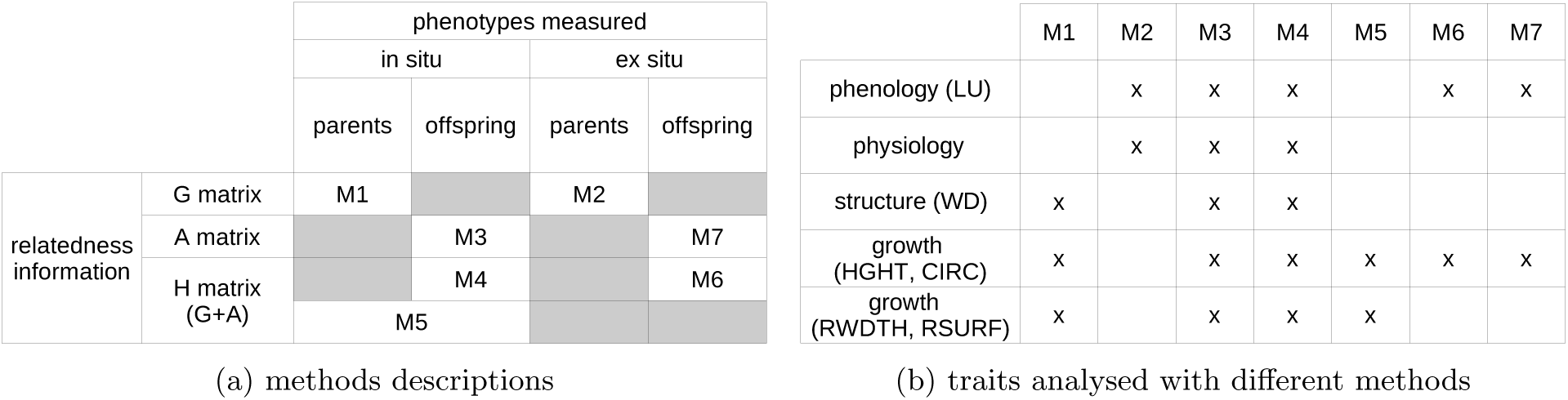
Description of the 7 methods used to estimate genetic parameters with the animal model, depending on the genetic relatedness information available and on the phenotypic measure conditions (a). Summary of the traits allowing comparisons across different methods (b).

### 2.7 Estimation of genetic correlations among traits

Finally, we estimated genetic correlations among traits. As different traits were assessed most of the time in different settings (Table 1), bivariate mixed model could hardly be implemented, due to different fixed effects used in different models. Moreover breedR algorithm showed convergence problem for bivariate models and bayesian analyses with MCMCglmm was strongly dependent on the priors distribution suggesting that most of the information in bivariate models comes from the priors. To overcome these difficulties, we extracted the BLUP (best linear unbiased predictor) of the breeding values from the univariate models random genetic effect. We then calculated Pearson correlation coefficients among breeding values of the different traits, although we acknowledge some drawbacks of using this method (Hadfield et al., 2010).

The R scripts used for these analyses and details on phenotypic measures are available as supplementary material.

## 3 Results

### 3.1 Comparison of performance of different experimental designs to measure heritability

#### 3.1.1 *In situ vs. ex situ* comparisons

A few traits were assessed at the same age *in situ* and *ex situ* (in the common garden progeny test). The comparison of methods M3 and M7 shows that for leaf unfolding, heritability is lower when estimated in common garden than *in situ* (Figure 3a), while a similar pattern can be observed in *Q.petraea* for height but not in *Q. robur*. However for all three traits (HGHT, CIRC and LUs), confidence intervals overlap over the two methods. Finally, confidence intervals are larger *in situ* than *ex situ*. There are clear unbalances in the sampling of parents between the two settings which may underpin the discrepancies of confidence intervals. Progeny tests comprise a limited number of families, but a large number of sibs within family, while *in situ* family samples show the opposite trend (Appendix 1 and 2). The comparison of *in situ* and *ex situ* estimation was extended to the case when genetic relatedness among sibs in the G2 included also relatedness of the parents, thus providing deeper relatedness information over two generations (Figure 3b). We found no noticeable difference between the two methods, neither for the heritability nor for its confidence interval.

**Figure 3:**
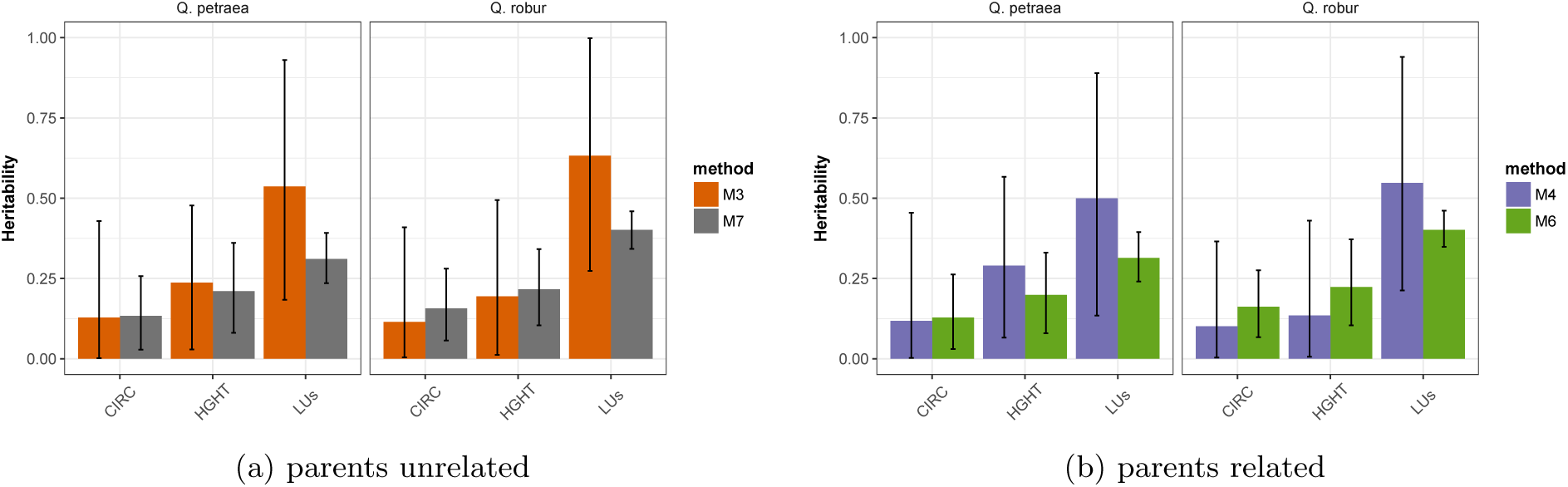
Comparison of heritability estimates 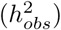 computed in G2 *in situ* (M3, M4) or in common garden (M6, M7) for *Q. petraea* and *Q. robur*, considering parents are non related (a) or taking account of genomic relatedness among parents (b). CIRC: circumference, HGHT height, LUs: leaf unfolding. Bars represent the estimates from REML and error bars represent the 95% CI obtained from 1000 bootstrap simulations.

#### 3.1.2 *In situ* heritability estimations in an unstructured vs. family structured population

We compared estimates obtained within the G1 generation (M1) with those obtained in the G2 generation (M3, M4) and with those obtained by combining both generations (M5). Heritability estimates based on unstructured natural populations (M1) exhibit on average very large confidence intervals and highly variable estimates for the same traits among species (Figure 4a) and show on average higher values than the other methods. Heterogeneity of estimates across methods may be generated by the poor precision of estimations. Estimates based on family structured sets in M3 and M4 provide similar estimates across species and reduced confidence intervals in comparison to M1. There was no evidence for improved reduction of confidence intervals with deeper genetic relatedness and combined phenotypic assessments over the two generations (M5 versus M3,M4 and M1).

We tested whether estimates are comparable between both generations by comparing estimates from M1 vs. M3 and M4 (Figure 4a) and M2 vs. M6 and M7 (Figure 4b). For growth and wood density, heritability is higher when estimated on G1 (Figure 4a). For leaf unfolding (measured in the conservation collection), heritability is also higher in generation G1 (Figure 4b), but in both cases associated confidence intervals are much larger in G1. Finally it is worthwhile noting that more precise pedigree information in the second generation (methods M3, M4, M5 M6 and M7) tends to increase heritability values (Appendix 6). As indicated in the Method section, we used all second generation individuals with at least one parent known for heritability estimation, but when we restricted the analysis to individuals with both parents known, heritability values tended to increase (Appendix 6).

#### 3.1.3 *In situ* heritability estimation using different G matrices

We compared heritability values based on the genomic relatedness matrix calculated with variable number of markers depending on the MAF threshold for SNP calling (from 1,500 SNPs with *MAF >* 40% to 32,500 with *MAF >* 1%, Figure 5). We selected a panel of several traits across the different categories assessed *in situ* in the first generation to illustrate changes in heritability estimates. Whatever the trait and the species, the estimates are very stable when markers were selected for MAF varying from 1% to 15% (number of markers varied between 33,000 to 6,700). Discrepancies among estimates increased when higher threshold values for MAF were used, thus corresponding to a lower number of markers (from 1,400 to 3,000). Hence, in what follows we considered only results obtained with the ***G*** matrix computed for SNPs selected at a *MAF* = 5%.

**Figure 4:**
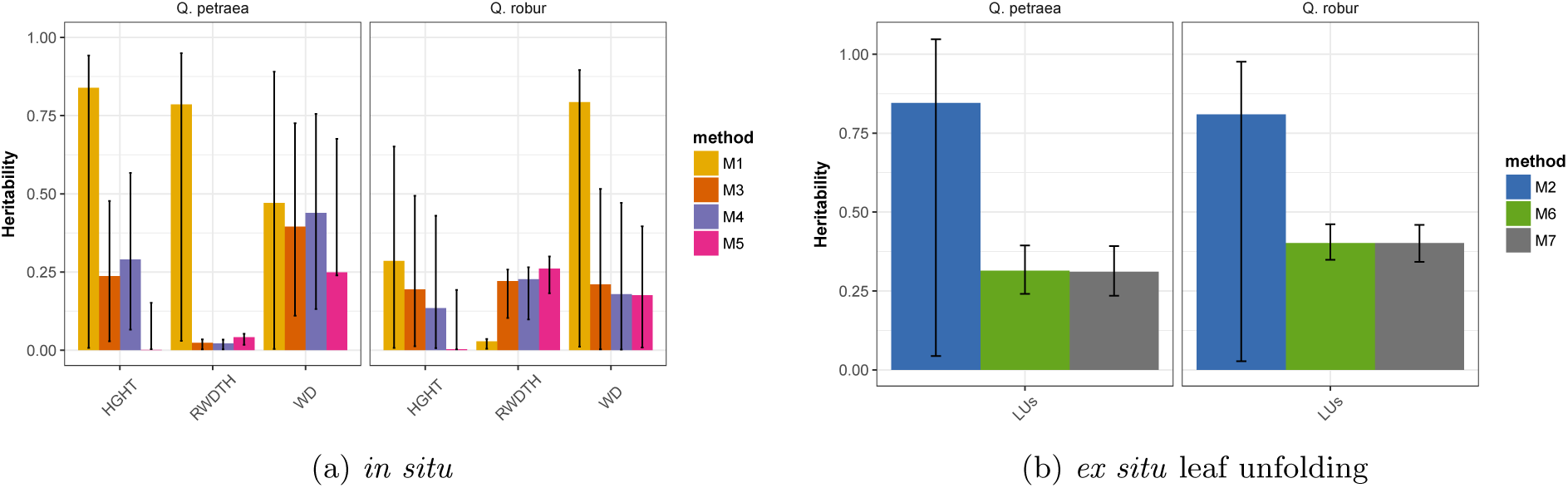
Comparison of heritability estimates 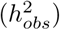 computed (a) *in situ* in G1 (M1), G2 (M3 and M4) or G1 and G2 (M5) for *Q. petraea* and *Q. robur*. HGHT: height, RWDTH: ring width, WD: wood density. and (b) *ex situ* for leaf unfolding (LUs) in G1 (M2) and G2 (M6, M7). Bars represent the estimates from REML and error bars represent the 95% CI obtained from 1000 bootstrap simulations.

**Figure 5:**
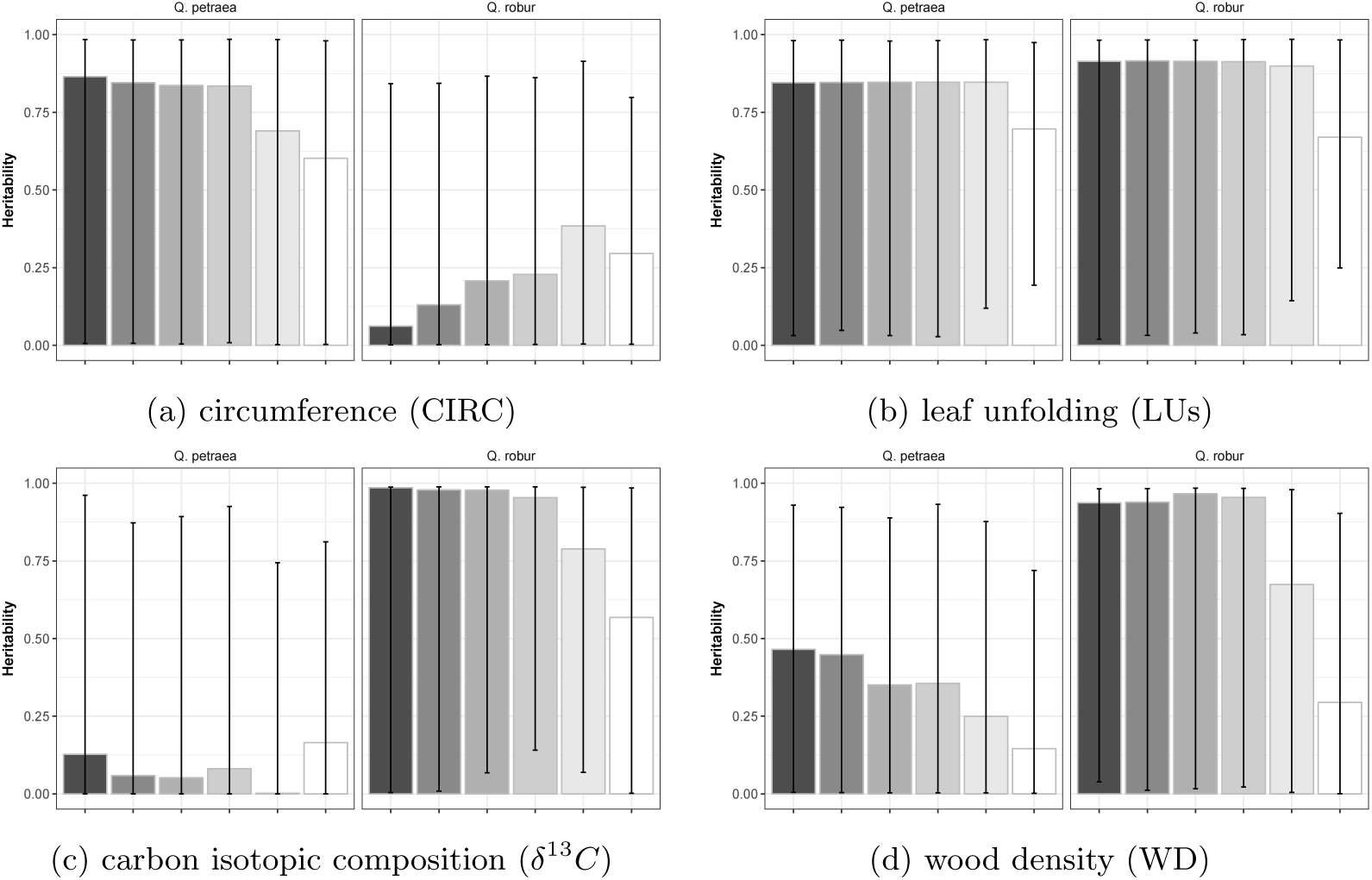
Comparison of heritability estimates 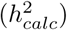 computed *in situ* in Gl (Ml or M2) with marker sets selected from different MAF thresholds, for circumference (a), leaf unfolding (b), carbon isotopic composition (c) and wood density (d) in *Q. petraea* and *Q. robur*. Bars represent the estimates of heritability and error bars represent the 95% CI obtained from 1GGG bootstrap simulations. MAF thresholds of 1%, 5%, 1G%, 15%, 3G% and 4G% correspond to 32G4T, 1B2T4, 9BG2, 6TB3, 2849 and 1454 markers in *Q. petraea* and 33131, 164G8, 1G225, T143, 3G58 and 1561 markers in *Q. robur*. The grey scale corresponds to MAF thresholds, from darker (MAF=1%) to lighter grey (MAF=4G%).

### 3.2 Genetic variation of traits measured *in situ*

#### 3.2.1 Heritability vs. evolvability

We performed a batch analysis to estimate heritability and evolvability of all traits measured *in situ*, or *ex situ* in the conservation collection (which is a replicated copy of the G1 generation) (methods M1 and M2 Table 1). Parameter estimation with methods M3 to M7 are presented in Appendix 5. There was at least one trait per species and category - except for resilience - that exhibited an important amount of genetic additive variance, encompassing a large range of variation of heritability (from 0.1219 to 0.8851 in *Q. petraea* and from 0.3871 to 0.8938 in *Q. robur*) with very large confidence intervals. For most traits, heritability computed with the raw phenotypic variance 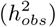 or the phenotypic variance computed as the sum of random effects variances 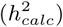 provided similar estimates, suggesting that the fixed effects included in the models did not significantly change the phenotypic variance, except for flowering dates for which 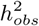 is far lower than 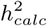.

Evolvability values showed also different trends of variation across traits in comparison to heritability and *I_a_* and *h*^2^ are not correlated (Figure 6). some traits harbour high heritability and low evolvability values (e.g. HGHT in *Q. petraea*) while other show the oposite pattern (e.g. MAR in *Q. petraea*).

#### 3.2.2 Differences between species and categories of traits

Regardless of the parameter considered (either heritability or evolvabilty) there is as much variation of genetic variation within a trait category than between the different categories. There is a slight trend towards lower genetic variation in secondary metabolite compounds and leaf morphological traits and higher genetic variation in phenology and growth traits, while physiological traits show intermediate values of heritability (Table 2 and 3). For resilience traits, phenotypic variance was negligibly caused by genetic variance (low values of heritability and evolvability).

When comparing genetic variation between the two species, no major trend emerges, except the very large discrepancy for growth traits and physiological traits. All growth traits showed striking larger heritabilities in *Q. petraea* than in *Q. robur*, and vice versa for physiological traits.

**Table 2:** Evolutionary parameters estimates for each trait, for *Q. petraea. V_p_*: observed inter-individual phenotypic variance; 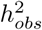: Heritability computed with the observed phenotypic variance; 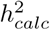: Heritability computed with the phenotypic variance estimated from the model (sum of variances of random effects); *I_a_:* Evolvability; method: the method as described in Figure 2a with which the trait was analysed; Δ*AIC*: Δ*AIC* between full model and model without additive effect (a positive value represent a better fit of the model with an additive effect). (exp) stands for traits with an exponential distribution. Numbers in brackets represent the 95% CI obtained from 1000 bootstrap simulations.

| class | trait | $V_p$ | $h_{obs}^2$ | $h_{calc}^2$ | $I_a$ | method | $\Delta AIC$ |
| --- | --- | --- | --- | --- | --- | --- | --- |
| Growth | **CIRC** | **948.387** | **0.7745 [0.006 - 0.9371]** | **0.8455 [0.0059 - 0.9831]** | **0.0202 [1e-04 - 0.0279]** | **M1** | **2.3397** |
|  | **HGHT** | **24960.9412** | **0.8393 [0.0075 - 0.9426]** | **0.9089 [0.0075 - 0.9845]** | **0.003 [0 - 0.0039]** | **M1** | **0.7856** |
|  | **RSURF** | **748389.0774** | **0.6382 [0.0171 - 0.7709]** | **0.3804 [0.0117 - 0.5217]** | **0.069 [0.0019 - 0.0879]** | **M1** | **1.7635** |
|  | **RWDTH** | **0.1973** | **0.786 [0.0295 - 0.9497]** | **0.1914 [0.008 - 0.2462]** | **0.0187 [7e-04 - 0.0234]** | **M1** | **2.6609** |
| R Success | **NOFF** | **81.1361** | **0.8237 [0.011 - 0.9238]** | **0.7829 [0.0107 - 0.977]** | **0.8791 [0.0151 - 2.2972]** | **M1** | **0.6427** |
| Phenology | **LU**s | **1.3245** | **0.8456 [0.0442 - 1.0474]** | **0.8465 [0.0481 - 0.9825]** | **0.1909 [0.0093 - 0.2966]** | **M2** | **16.0096** |
|  | **LU**d | **22.7381** | **0.8598 [0.026 - 0.9508]** | **0.9138 [0.0241 - 0.9881]** | - | **M2** | **20.5593** |
|  | **LS** | **15.3089** | **0.827 [0.019 - 0.8977]** | **0.8714 [0.0197 - 0.9793]** | - | **M2** | **3.4352** |
|  | **GSL** | **34.7263** | **0.8172 [0.0139 - 0.9388]** | **0.8537 [0.0145 - 0.9856]** | **0.0007 [1e-05 - 0.0009]** | **M2** | **10.9387** |
|  | FFLW | 47.5872 | 0.2948 [0.0025 - 0.5686] | 0.9547 [0.0189 - 0.9786] | - | M1 | -0.7387 |
|  | **MFLW** | **35.043** | **0.4637 [0.007 - 0.6614]** | **0.9826 [0.0173 - 0.9841]** | - | **M1** | **5.2571** |
|  | MAR | 2.293 | 0.0893 [1e-04 - 0.9216] | 0.2191 [1e-04 - 0.9634] | 0.3268 [2e-04 - 1.6089] | M1 | -1.8562 |
| Physiology | C | 785.3835 | 0.234 [1e-04 - 0.9187] | 0.2357 [1e-04 - 0.9582] | 9e-04 [0 - 0.0036] | M2 | -1.9429 |
|  | C/N | 12.5891 | 0.1041 [1e-04 - 0.9062] | 0.1046 [1e-04 - 0.9447] | 0.0022 [0 - 0.0192] | M2 | -1.9591 |
| | $\delta^{13}C$ | 1.304 | 0.0585 [1e-04 - 0.8374] | 0.0587 [1e-04 - 0.8725] | - | M2 | -1.9814 |
| | $\delta^{15}N$ | **1.8222** | **0.8638 [0.0035 - 0.9487]** | **0.8808 [0.0035 - 0.9886]** | - | **M2** | **0.8272** |
|  | N | 6.3556 | 0.1348 [1e-04 - 0.9018] | 0.1355 [1e-04 - 0.9449] | 0.0024 [0 - 0.0163] | M2 | -1.9491 |
|  | SLA | 5.2619 | 2e-04 [1e-04 - 0.8532] | 2e-04 [1e-04 - 0.8677] | 0 [0 - 0.0309] | M2 | -2.001 |
|  | MLA | 111.6555 | 0.4782 [2e-04 - 0.9333] | 0.4869 [2e-04 - 0.982] | 0.0266 [0 - 0.0599] | M2 | -0.778 |
| Resilience | REC | 0.0223 | 0.0015 [1e-04 - 0.2922] | 0.0788 [0.0167 - 0.9561] | 0 [0 - 0.0037] | M1 | -2.0056 |
|  | REL | 0.0103 | 0.0464 [2e-04 - 0.2983] | 0.2547 [0.0107 - 0.9488] | 7e-04 [0 - 0.0042] | M1 | -1.8761 |
|  | RET | 0.0057 | 0.0854 [1e-04 - 0.354] | 0.9533 [0.0187 - 0.968] | 0.0011 [0 - 0.0047] | M1 | -1.6311 |
| Structure | WD | 886.685 | 0.4707 [0.0038 - 0.8906] | 0.448 [0.0038 - 0.9222] | 0.0013 [0 - 0.0027] | M1 | -0.6863 |
| Leaf | **BS** | **0.6307** | **0.6844 [0.0114 - 0.9068]** | **0.9031 [0.0161 - 0.9853]** | **0.0275 [4e-04 - 0.0369]** | **M1** | **6.0746** |
| Morphology | HR | 0.7707 | 0.0889 [0.0011 - 0.1464] | 0.117 [0.0181 - 0.9782] | 0.006 [1e-04 - 0.01] | M1 | -1.8682 |
|  | LDR | 31.5929 | 0.0757 [0.0017 - 0.1667] | 0.095 [0.0083 - 0.7282] | 0.0035 [1e-04 - 0.0076] | M1 | -1.978 |
|  | LL | 310.0745 | 0.1613 [0.0028 - 0.2354] | 0.1995 [0.0128 - 0.9459] | 0.0058 [1e-04 - 0.0087] | M1 | -1.7996 |
|  | LW | 40.2459 | 0.0743 [0.0015 - 0.1263] | 0.0841 [0.0107 - 0.8963] | 0.0037 [1e-04 - 0.0063] | M1 | -1.9517 |
|  | LWR | 11.3129 | 0.4454 [0.0074 - 0.5917] | 0.5172 [0.0165 - 0.9874] | 0.0054 [1e-04 - 0.0071] | M1 | -0.7868 |
|  | NL | **2.3663** | **0.7801 [0.015 - 0.989]** | **0.978 [0.0209 - 0.9879]** | **0.0147 [3e-04 - 0.019]** | **M1** | **2.7978** |
|  | NV | **0.2549** | **0.7607 [0.0113 - 1.0068]** | **0.9647 [0.0166 - 0.987]** | **1.8582 [0.0284 - 3.2702]** | **M1** | **0.2139** |
|  | OB | 21.6011 | 0.1021 [0.0025 - 0.3051] | 0.225 [0.0074 - 0.697] | 8e-04 [0 - 0.0023] | M1 | -1.7428 |
|  | PL | 25.8648 | 0.0086 [4e-04 - 0.0367] | 0.0098 [0.0046 - 0.3983] | 8e-04 [0 - 0.0033] | M1 | -2.0532 |
|  | PR | 9.1948 | 0.0138 [0 - 0.0443] | 0.0149 [0.0138 - 0.9627] | 5e-04 [0 - 0.0017] | M1 | -2.0375 |
|  | **PV** | **31.5445** | **0.8851 [0.0172 - 1.1388]** | **0.9762 [0.0198 - 0.9877]** | **2.95 [0.053 - 5.4005]** | **M1** | **1.1456** |
|  | SW | 20.674 | 0.019 [4e-04 - 0.0541] | 0.0209 [0.0068 - 0.7697] | 9e-04 [0 - 0.0026] | M1 | -2.0273 |
|  | WP | 120.6494 | 0.4605 [0.0085 - 0.6309] | 0.6686 [0.0157 - 0.9824] | 0.0219 [4e-04 - 0.0302] | M1 | -0.7932 |
| Defence | CNFL (exp) | 3.8295 | 0.0121 [7e-04 - 0.6977] | 0.0157 [7e-04 - 0.7051] | 0.0048 [2e-04 - 0.2036] | M1 | -2.01 |
|  | CSTG (exp) | 23.8251 | 0.0034 [2e-04 - 0.8427] | 0.0036 [2e-04 - 0.8572] | 1e-04 [0 - 0.0147] | M1 | -2.0219 |
|  | CSTL (exp) | 1.1465 | 3e-04 [1e-04 - 0.7069] | 3e-04 [1e-04 - 0.7276] | 1e-04 [0 - 0.2349] | M1 | -2.0009 |
|  | **CWSK (exp)** | **54.4825** | **0.1219 [1e-04 - 0.2756]** | **0.1493 [1e-04 - 0.3011]** | **1.529 [0.001 - 2.7502]** | **M1** | **0.8847** |
|  | EGNL (exp) | 0.4896 | 0.0084 [1e-04 - 0.4794] | 0.0146 [1e-04 - 0.4979] | 0.0305 [1e-04 - 1.0222] | M1 | -2.0111 |
|  | ELAC (exp) | 1.4765 | 0.4049 [2e-04 - 0.8256] | 0.4476 [2e-04 - 0.8673] | 0.1354 [1e-04 - 0.2986] | M1 | -0.0089 |
|  | ELTOT (exp) | 261.8303 | 3e-04 [1e-04 - 0.7706] | 3e-04 [1e-04 - 0.7799] | 1e-04 [0 - 0.1629] | M1 | -2.0009 |
|  | GRDN | 9.1947 | 0.2225 [2e-04 - 0.9266] | 0.2222 [2e-04 - 0.9655] | 0.0313 [0 - 0.1418] | M1 | -1.7102 |
|  | MVL (exp) | 0.5417 | 1e-04 [1e-04 - 0.7283] | 2e-04 [1e-04 - 0.7364] | 1e-04 [0 - 0.3001] | M1 | -2.0009 |
|  | PNTL (exp) | 0.018 | 0.0404 [1e-04 - 0.7852] | 0.056 [1e-04 - 0.804] | 0.0165 [0 - 0.238] | M1 | -1.9896 |
|  | ROBA (exp) | 2.3083 | 0.0941 [0.0044 - 0.6617] | 0.0914 [0.0043 - 0.6517] | 0.0102 [4e-04 - 0.0678] | M1 | -1.92 |
|  | ROBB | 5.7242 | 0.0087 [4e-04 - 0.7703] | 0.0082 [4e-04 - 0.765] | 0.001 [0 - 0.0951] | M1 | -2.0164 |
|  | ROBC (exp) | 7.7695 | 0.032 [2e-04 - 0.7902] | 0.0369 [2e-04 - 0.8021] | 0.006 [0 - 0.1215] | M1 | -1.9922 |
|  | ROBD | 11.6933 | 0.1231 [1e-04 - 0.9139] | 0.1177 [1e-04 - 0.9456] | 0.0191 [0 - 0.1537] | M1 | -1.9537 |
|  | ROBE (exp) | 5.447 | 0.0105 [0.0018 - 0.7153] | 0.0109 [0.0018 - 0.7139] | 9e-04 [1e-04 - 0.0527] | M1 | -2.0186 |
|  | SYRG | 6.4986 | 0.0709 [1e-04 - 0.9207] | 0.1107 [1e-04 - 0.9626] | 0.0163 [0 - 0.1421] | M1 | -1.9701 |
|  | TWSK (exp) | 38.728 | 0.1016 [6e-04 - 0.2121] | 0.0902 [6e-04 - 0.2317] | 1.405 [0.0085 - 2.853] | M1 | -1.1858 |
|  | VNL | 2.5027 | 0.1733 [3e-04 - 0.8947] | 0.2257 [3e-04 - 0.9307] | 0.0468 [1e-04 - 0.2088] | M1 | -1.9173 |
|  | VSCG | 23.872 | 0.0081 [1e-04 - 0.8931] | 0.0082 [1e-04 - 0.9064] | 6e-04 [0 - 0.0638] | M1 | -2.0011 |
|  | VSCL (exp) | 4.2427 | 0.0016 [1e-04 - 0.7322] | 0.0025 [1e-04 - 0.7693] | 0.001 [0 - 0.2957] | M1 | -2.0073 |
|  | 2PHL (exp) | 0.0072 | 0.0029 [1e-04 - 0.7588] | 0.0039 [1e-04 - 0.7791] | 0.0015 [0 - 0.2844] | M1 | -2.0087 |

**Table 3:** Evolutionary parameters estimates for each trait, for *Q. robur. V_p_*: observed inter-individual phenotypic variance; 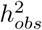: Heritability computed with the observed phenotypic variance; 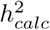: Heritability computed with the phenotypic variance estimated from the model (sum of variances of random effects); *I_a_:* Evolvability; method: the method as described in Figure 2a with which the trait was analysed; Δ*AIC*: Δ*AIC* between full model and model without additive effect (a positive value represent a better fit of the model with an additive effect). (exp) stands for traits with an exponential distribution. Numbers in brackets represent the 95% CI obtained from 1000 bootstrap simulations.

| class | trait | $V_p$ | $h_{obs}^2$ | $h_{calc}^2$ | $I_a$ | method | $\Delta AIC$ |
| --- | --- | --- | --- | --- | --- | --- | --- |
| Growth | CIRC | 554.8436 | 0.1241 [0.0019 - 0.7982] | 0.1308 [0.0019 - 0.8438] | 0.0025 [0 - 0.0155] | M1 | -1.9547 |
|  | HGHT | 26338.5265 | 0.2857 [0.0076 - 0.6516] | 0.2081 [0.0071 - 0.6294] | 0.0012 [0 - 0.0031] | M1 | -1.2132 |
|  | RSURF | 351707.3123 | 0.0171 [0.0022 - 0.0223] | 0.0087 [0.0018 - 0.0214] | 0.0015 [2e-04 - 0.0021] | M1 | -2.0624 |
|  | RWDTH | 0.1383 | 0.0283 [0.0044 - 0.0359] | 0.0059 [0.0011 - 0.0095] | 6e-04 [1e-04 - 8e-04] | M1 | -2.1188 |
| R Success | NOFF | 26.9444 | 0.1094 [0.0019 - 0.8008] | 0.1061 [0.0019 - 0.8353] | 0.0794 [0.001 - 0.7408] | M1 | -1.8921 |
| Phenology | **LUs** | **1.5921** | **0.809 [0.0273 - 0.9762]** | **0.9154 [0.0319 - 0.9826]** | **0.114 [0.0039 - 0.1399]** | **M2** | **5.9259** |
|  | **LUd** | **39.1092** | **0.8198 [0.0294 - 0.9099]** | **0.9005 [0.0315 - 0.9853]** | - | **M2** | **2.1757** |
|  | LS | 14.4259 | 0.0729 [0.0069 - 0.2854] | 0.0663 [0.0088 - 0.3961] | - | M2 | -1.9854 |
|  | **GSL** | **43.1537** | **0.7598 [0.0179 - 0.8593]** | **0.7823 [0.0193 - 0.9767]** | **8e-04 [0 - 0.001]** | **M2** | **0.6816** |
|  | **FFLW** | **27.054** | **0.4983 [0.0109 - 0.655]** | **0.9713 [0.0223 - 0.9823]** | - | **M1** | **2.5067** |
|  | **MFLW** | **32.2575** | **0.5236 [0.0113 - 0.6889]** | **0.9717 [0.0199 - 0.9827]** | - | **M1** | **4.3772** |
|  | MAR | 2.9028 | 0.2226 [0.0014 - 0.8969] | 0.7126 [0.0014 - 0.9811] | 0.6097 [0.0012 - 1.0633] | M1 | -0.4242 |
| Physiology | C | 66.8823 | 0.7415 [9e-04 - 0.9049] | 0.7757 [9e-04 - 0.9865] | 2e-04 [0 - 3e-04] | M2 | -0.4399 |
|  | C/N | 12.7377 | 5e-04 [1e-04 - 0.9293] | 5e-04 [1e-04 - 0.9782] | 0 [0 - 0.0205] | M2 | -2.0013 |
| | $\delta^{13}C$ | **1.1119** | **0.8733 [0.0088 - 0.9029]** | **0.9784 [0.0088 - 0.9882]** | - | **M2** | **8.3192** |
| | $\delta^{15}N$ | **1.5707** | **0.8938 [0.0024 - 0.9052]** | **0.9743 [0.0024 - 0.9877]** | - | **M2** | **4.7475** |
|  | N | 6.0234 | 0.0057 [1e-04 - 0.8576] | 0.0057 [1e-04 - 0.902] | 1e-04 [0 - 0.0126] | M2 | -2.001 |
|  | **SLA** | **9.381** | **0.8768 [0.0293 - 0.9048]** | **0.9415 [0.0293 - 0.9875]** | **0.0435 [0.0013 - 0.0546]** | **M2** | **2.5715** |
|  | **MLA** | **104.5311** | **0.656 [4e-04 - 0.9018]** | **0.6905 [4e-04 - 0.9836]** | **0.0536 [0 - 0.0877]** | **M2** | **0.385** |
| Resilience | REC | 0.1241 | 0.1315 [2e-04 - 0.3353] | 0.7537 [0.0191 - 0.9609] | 0.0062 [0 - 0.0161] | M1 | -1.308 |
|  | REL | 0.0515 | 0.0035 [0.0016 - 0.208] | 0.0128 [0.008 - 0.607] | 2e-04 [1e-04 - 0.0157] | M1 | -2.0389 |
|  | RET | 0.0141 | 0.0029 [4e-04 - 0.1906] | 0.0194 [0.0107 - 0.9043] | 1e-04 [0 - 0.0082] | M1 | -2.0336 |
| Structure | **WD** | **756.3841** | **0.7932 [0.0114 - 0.8962]** | **0.9393 [0.0113 - 0.9828]** | **0.002 [0 - 0.0025]** | **M1** | **3.3434** |
| Leaf | **BS** | **1.5555** | **0.4737 [0.0126 - 0.6188]** | **0.47 [0.0133 - 0.756]** | **0.017 [5e-04 - 0.0233]** | **M1** | **2.8342** |
| Morphology | HR | 0.4631 | 0.0695 [0.0015 - 0.0942] | 0.0817 [0.023 - 0.981] | 0.0182 [4e-04 - 0.0251] | M1 | -1.8387 |
|  | **LDR** | **79.3401** | **0.6417 [0.0137 - 0.7857]** | **0.9607 [0.0213 - 0.9855]** | **0.0276 [6e-04 - 0.0339]** | **M1** | **2.0453** |
|  | LL | 199.7889 | 0.3828 [0.0085 - 0.4777] | 0.4482 [0.0232 - 0.9859] | 0.0124 [3e-04 - 0.0155] | M1 | -0.0828 |
|  | **LW** | **26.5269** | **0.5018 [0.0139 - 0.6281]** | **0.6213 [0.0289 - 0.9859]** | **0.0217 [6e-04 - 0.027]** | **M1** | **1.3548** |
|  | **LWR** | **8.436** | **0.6239 [0.015 - 0.7846]** | **0.9285 [0.0234 - 0.9832]** | **0.0053 [1e-04 - 0.0067]** | **M1** | **0.4713** |
|  | NL | 1.7074 | 0.2549 [0.0057 - 0.3667] | 0.3231 [0.0192 - 0.9655] | 0.0057 [1e-04 - 0.0083] | M1 | -1.5014 |
|  | NV | 2.1461 | 0.4809 [0.0119 - 0.6151] | 0.659 [0.0211 - 0.9623] | 0.1225 [0.003 - 0.1835] | M1 | -0.3589 |
|  | **OB** | **26.9943** | **0.3871 [0.0079 - 0.5724]** | **0.9737 [0.0231 - 0.9776]** | **0.0032 [1e-04 - 0.0047]** | **M1** | **0.9377** |
|  | PL | 4.5531 | 0.0978 [0.0021 - 0.1346] | 0.1075 [0.022 - 0.9793] | 0.0207 [4e-04 - 0.0288] | M1 | -1.9069 |
|  | PR | 5.0951 | 0.4037 [0.0118 - 0.5036] | 0.481 [0.0268 - 0.9852] | 0.0721 [0.002 - 0.0902] | M1 | -1.2302 |
|  | PV | 274.5848 | 0.5277 [0.0133 - 0.6833] | 0.6929 [0.017 - 0.9604] | 0.1265 [0.0033 - 0.1969] | M1 | -0.4726 |
|  | SW | 8.478 | 0.2524 [0.0052 - 0.3353] | 0.3142 [0.0204 - 0.9833] | 0.0113 [2e-04 - 0.0151] | M1 | -1.1659 |
|  | **WP** | **81.5738** | **0.662 [0.0154 - 0.8466]** | **0.9353 [0.0242 - 0.9855]** | **0.0271 [6e-04 - 0.0351]** | **M1** | **3.2263** |
| Defence | CNFL (exp) | 6.9258 | 0.0016 [1e-04 - 0.6908] | 0.0029 [1e-04 - 0.7173] | 0.0011 [1e-04 - 0.2613] | M1 | -2.0061 |
|  | CSTG (exp) | 19.9558 | 0.012 [2e-04 - 0.8275] | 0.0145 [2e-04 - 0.8701] | 2e-04 [0 - 0.0122] | M1 | -2.0012 |
|  | **CSTL (exp)** | **1.308** | **0.6977 [0.0135 - 0.7892]** | **0.8293 [0.0134 - 0.8738]** | **0.2337 [0.0032 - 0.2793]** | **M1** | **5.5538** |
|  | CWSK (exp) | 0.0807 | 0.0283 [1e-04 - 0.4838] | 0.03 [1e-04 - 0.5346] | 0.0577 [2e-04 - 0.936] | M1 | -1.9961 |
|  | EGNL (exp) | 0.1494 | 0.0326 [2e-04 - 0.5273] | 0.0787 [2e-04 - 0.5714] | 0.102 [3e-04 - 0.7043] | M1 | -1.9502 |
|  | ELAC (exp) | 0.7528 | 0.0956 [4e-04 - 0.7563] | 0.1048 [4e-04 - 0.7964] | 0.0209 [1e-04 - 0.1533] | M1 | -1.9036 |
|  | ELTOT (exp) | 393.525 | 0.005 [2e-04 - 0.7187] | 0.0058 [2e-04 - 0.7285] | 0.001 [0 - 0.1225] | M1 | -2.0132 |
|  | GRDN | 8.3491 | 0.5413 [5e-04 - 0.894] | 0.5555 [5e-04 - 0.9798] | 0.1051 [1e-04 - 0.208] | M1 | -0.8921 |
|  | MVL (exp) | 0.2823 | 0.0985 [0.0021 - 0.6133] | 0.0937 [0.0021 - 0.6516] | 0.0624 [0.0011 - 0.401] | M1 | -1.9168 |
|  | PNTL (exp) | 0.0262 | 0.0361 [6e-04 - 0.6607] | 0.0518 [6e-04 - 0.6917] | 0.0233 [3e-04 - 0.2924] | M1 | -1.9722 |
|  | ROBA (exp) | 6.348 | 0.0027 [2e-04 - 0.7385] | 0.0034 [2e-04 - 0.7804] | 4e-04 [0 - 0.0851] | M1 | -2.0272 |
|  | ROBB | 4.3557 | 0.2865 [3e-04 - 0.881] | 0.292 [3e-04 - 0.9548] | 0.0314 [0 - 0.1084] | M1 | -1.6943 |
|  | ROBC (exp) | 6.2893 | 0.1827 [2e-04 - 0.7875] | 0.1978 [2e-04 - 0.8566] | 0.0444 [0 - 0.1954] | M1 | -1.7505 |
|  | ROBD | 12.5036 | 0.004 [2e-04 - 0.7879] | 0.0042 [2e-04 - 0.8345] | 7e-04 [0 - 0.1294] | M1 | -2.0014 |
|  | ROBE (exp) | 3.3626 | 0.5705 [0.0024 - 0.8595] | 0.7185 [0.0024 - 0.945] | 0.0603 [2e-04 - 0.0897] | M1 | -0.5345 |
|  | SYRG | 12.5141 | 0.0051 [2e-04 - 0.8473] | 0.0071 [2e-04 - 0.8994] | 0.0014 [0 - 0.169] | M1 | -2.0067 |
|  | **TWSK (exp)** | **0.3077** | **0.4144 [0.0014 - 0.5226]** | **0.4587 [0.0014 - 0.6016]** | **0.9215 [0.0027 - 1.2623]** | **M1** | **0.9468** |
|  | VNL | 3.8353 | 2e-04 [2e-04 - 0.8456] | 3e-04 [2e-04 - 0.8922] | 1e-04 [0 - 0.1648] | M1 | -2.0013 |
|  | VSCG | 25.0507 | 0.5253 [0.001 - 0.89] | 0.5782 [0.001 - 0.977] | 0.0249 [0 - 0.0455] | M1 | -0.4071 |
|  | **VSCL (exp)** | **3.3081** | **0.6084 [0.0138 - 0.7703]** | **0.7989 [0.0131 - 0.8591]** | **0.2731 [0.0038 - 0.3283]** | **M1** | **2.3264** |
|  | 2PHL (exp) | 0.0179 | 1e-04 [1e-04 - 0.6675] | 2e-04 [1e-04 - 0.7121] | 1e-04 [1e-04 - 0.3599] | M1 | -2.0013 |

**Figure 6:**
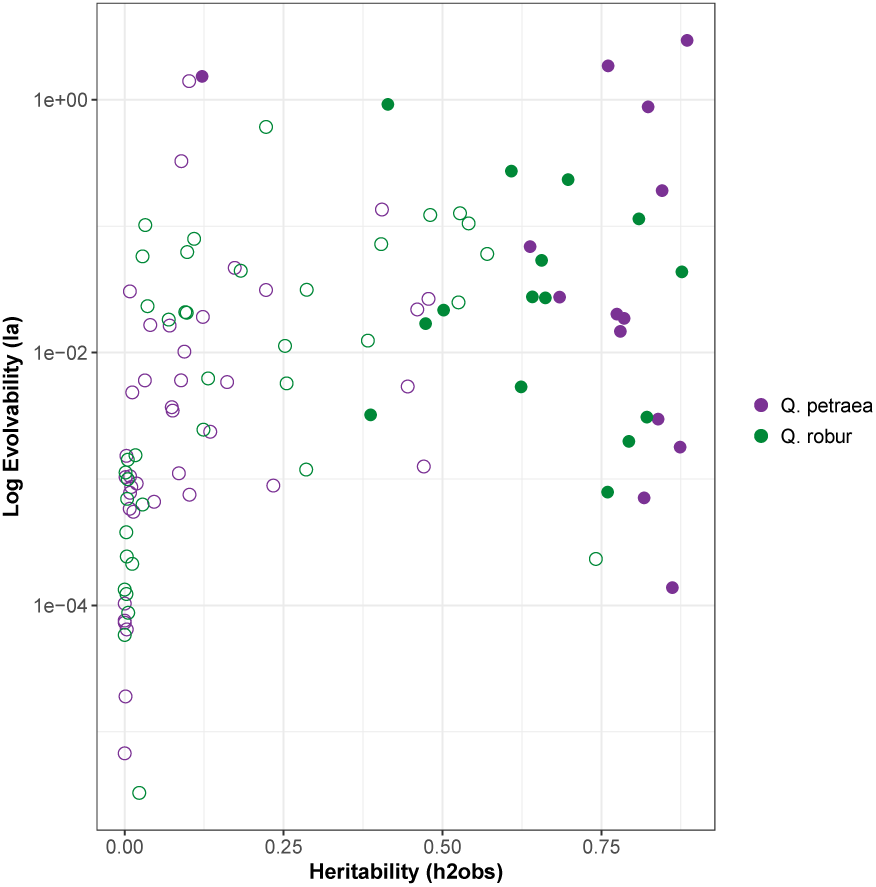
Plot of Evolvability against Heritability for each trait presented in Tables 2 and 3. Open circles represent traits for which the genetic additive effect does not increase model fitting (Δ*AIC <* 0) and filled circles represent traits for which the genetic additive effect increases model fitting (Δ*AIC* > 0). Purple: *Q. petraea*, green: *Q*. *robur*. Evolvability is presented on a log-scale to enhance data visualization.

### 3.3 Genetic correlations among traits

We computed genetic correlations based on BLUPs estimates for the subset of traits that were assessed in the two successive generations (G1 and G2), thus allowing comparisons across different settings with deeper relatedness information (Figure 7). For estimating genetic correlations, we privileged traits for which multiple observations were available in different experimental settings, given the overall low precision of heritability estimates suggesting that BLUPs may as well be estimated with low precision. We found consistent significant genetic correlations between all growth traits in both species and in both experimental settings (either M1and M2 on the one hand or M4 on the other hand), which are expected given the allometric relationship between diameter and height. There was no correlation between growth traits and wood density (WD). LUs is negatively correlated to growth in the first generation, while the correlation becomes positive in the second generation for *Q. robur*. Finally there were also consistent negative correlations among some physiological related traits, as SLA (Specific Leaf Area) with *δ*^13^*C* (carbon isotopic composition) and with C/N (ratio Carbon/Nitrogen content). These correlations were found in M1 or M2 (adult phenotypic traits) and also in M4 (juvenile phenotypic traits) and there were only slight differences between *Q. petraea* and *Q. robur*. Genetic correlations between all traits within each category are attached in Appendix 7, as estimated with M1 or M2. As expected, traits are highly correlated within a given category, for example growth traits and leaf morphology traits, particularly those that are related to leaf shape (LL,WP,SW). Flowering phenological traits (FFLW, MFLW) are also correlated to vegetative bud phenology (LUs, GSL). The length of the growing season is correlated both to the leaf unfolding and leaf senescence. Leaf retention (MAR) is correlated to senescence in both species and to flowering dates in *Q. petraea* only. A few secondary metabolites exhibit strong genetic correlations which cluster in two groups (VNL, CNFL, SYRG, PNTL, 2PHL) ont the one hand and (CSTG, VSCL, VSCG) on the other hand. Interestingly these two groups exhibit also negative correlations. However, except for growth traits, the coefficients of genetic correlations are relatively low.

**Figure 7:**
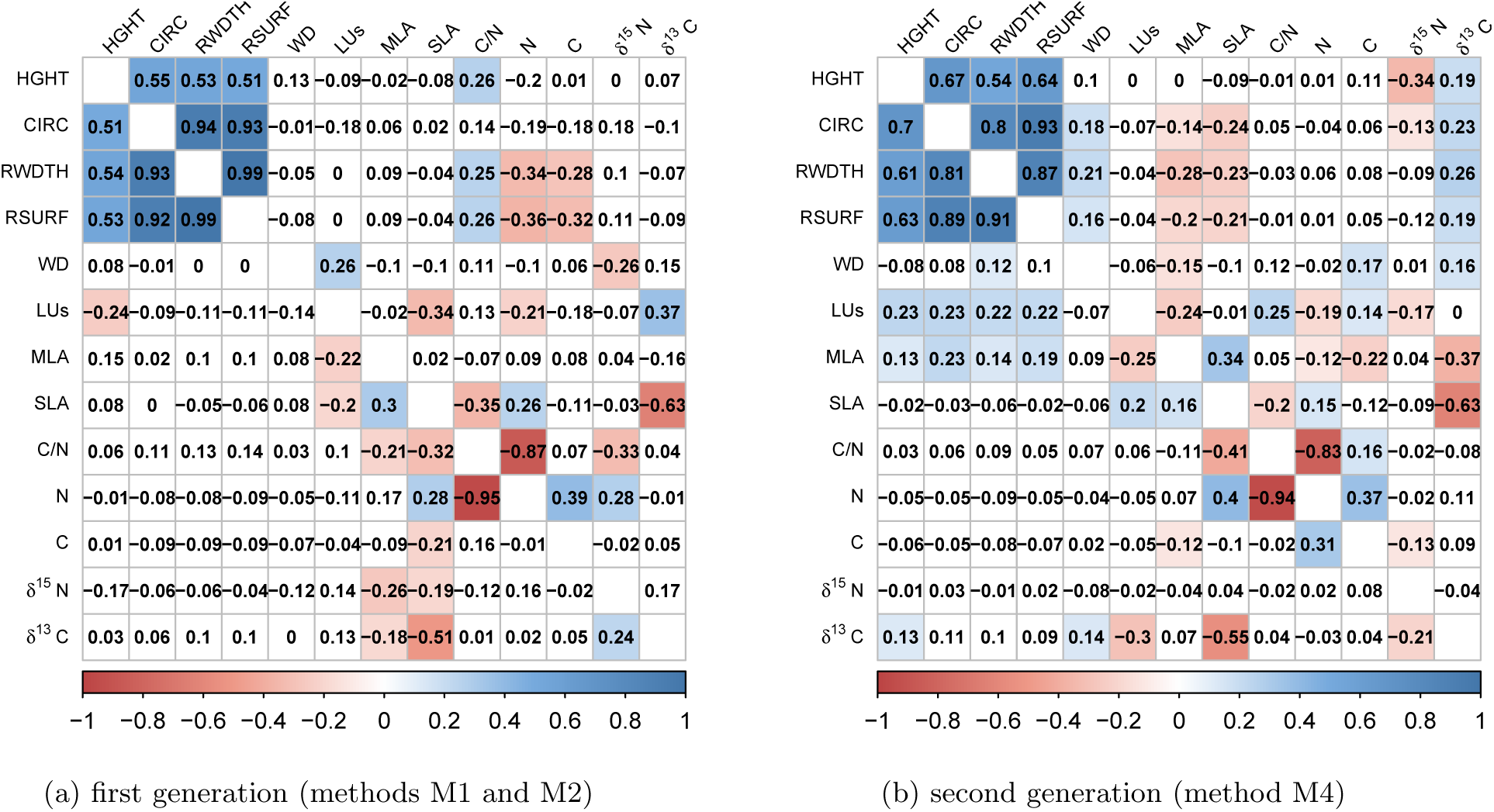
Genetic correlations for traits measured in both generations. (a) correlations for the first generation, (b) correlations for the second generation. Correlation coefficients for *Q. petraea* are above the matrix diagonals and *Q. robur* are below the diagonal. Colors correspond to the correlation sign (blue for positive and red for negative correlations). Only correlations significant at a 5% threshold are colored.

## 4 Discussion

In the present study we attempted to assess the evolutionary potential of phenotypic traits by estimating heritability, evolvability and genetic correlations in two different species of a mixed oak stand from North Western France. Our main objective was to explore ways of implementing different estimation methods directly in natural populations. We therefore compared results under different experimental and ecological settings to which forest geneticists have access. We addressed in our comparisons mean estimates and their confidence intervals of the relevant genetic parameters. Despite substantial variations of heritability and evolvability estimates obtained with different methods and their large confidence intervals, there are important findings which can be summarized in five major outcomes, and which may help to improve future investigations: (i) *in situ* show wider confidence intervals than *ex situ* estimates, (ii) assessments over two generations improve estimates of heritability, (iii) current sampling strategies are exposed to very large confidence intervals, (iv) levels of genetic variation varies moderately across different functional groups of traits, (v) genetic correlations are conserved in the two species.

### 4.1 *In situ* versus *ex situ* estimates

We included in our comparative analysis the more traditional procedure that consisted in raising progeny trials with known parentage and under experimental designs with replicated blocks. Thus our comparisons considered estimates obtained with the same base population, and their offspring stemming from open pollinated progenies that grew in the same forest, either under the canopy of the parents after natural dispersion (*in situ*) or nearby in a plantation installed according an experimental design (*ex situ*). We found wider confidence intervals for heritabilities in the *in situ* setting than in the *ex situ* setting (Figure 3), although they overlap. Everything else being equal, the differences between the two settings include differences of sampling sizes (number of progenies, and number of offspring within progenies), differences in the spatial distribution of offspring, and differences in microenvironmental conditions. The *ex situ* experiment comprised a low number of progenies (less than 25 female parents), but a large number of offspring per female parent in contrast to the *in situ* case that comprised a very large number of families and a reduced number of offspring (less than 3 per family). The larger sample size in the *ex situ* setting can explain the narrower confidence interval. Finally differences in confidence intervals may also be due to the higher environmental variances in non controlled conditions, eg *in situ*. The difference in heritability estimates between *in situ* and *ex situ* is quite low for growth traits, but more important for LUs, while CI overlap. The experimental differences between the two settings should have no effect on the heritability estimates (bias), unless the parents on which acorns were collected for the *ex situ* experiment were somehow selected. This cannot be entirely excluded as large trees may bear more seeds and may have been unintentionally preferred during the acorn harvests operations. Selection of parents may indeed lead to underestimation of the additive variance (Visscher et al., 2008; Ponzoni and James, 1978). Spatial distributions of the offspring differ substantially between the *ex situ* and the *in situ* settings. In the former, offspring are grouped in plots which are replicated in blocks, resulting in an almost completely randomized design. In the latter, offspring are preferentially grouped as seed are usually disseminated at short distances, however occasionally distant offspring resulting from pollen dispersion of the parental gametes were also sampled when possible. Preferential sib grouping may result in confounding of genetic and environmental effects thus inflating the genetic variance. We attempted to limit the bias due to common environmental effects in three different ways: by explicitly introducing a random spatial component in the mixed model (Kruuk and Hadfield, 2007), by accounting for microenvironmental covariates, and by sampling as much as possible distant offspring (this is similar to cross-fostering used in birds (Kruuk and Hadfield, 2007). Finally, despite that the two experiments were located in the same forest (Appendix 2), they occupy different parcels with potential differences in microenvironmental variation (that is accounted for by the spatial effect in model (M1) and (M2)). We suspect that either one of the three causes or their combination may underlie the differences observed in heritability values between the *ex situ* and *in situ* setting, but observe however that the relative ranking of heritability values among traits is conserved in the two experiments.

### 4.2 Assessments over successive generations

The traditional approach in estimating the genetic variance of traits in a tree population with no background genetic information is to raise open pollinated progenies, assuming that parents were unrelated, and using phenotypic information of the offspring only. Here we upgraded this approach in two different ways: by reconstructing genetic relatedness over the two generations (genomic relatedness between parents plus pedigree relationships between parent-offspring and among offspring, ***H*** matrix in (7), and by benefiting of phenotypic information of parents and offspring. Our results show that reconstructing genetic relatedness gave quasi-identical values of heritability whether comparisons were made *in situ* (M3 and M4) or *ex situ* (M6 and M7), see Figures 3 and 4. Theory and simulations predict that heritability is underestimated in the presence of pedigree errors when relatedness is poorly estimated (Morrissey et al., 2007). Experimental results obtained in natural populations show no significant change (Berénos et al., 2014; Akesson et al., 2008), or a slight increase of heritability values when deeper and more precise relatedness is available (Perrier et al., 2018). Indeed we noticed that better information of pedigree relationships increases heritability values. This was the case when we restricted heritability estimation in the second generation for offspring for which both parents were identified instead of including also offspring with one parent unknown (Appendix 6). Combining phenotypic information over two successive generations resulted in a decrease of heritability estimates compared to measures on one generation only (M5 on Figure 4a). It is worthwhile recalling that phenotypic records were taken at different ages in the two generations, given the obvious age structure in our study populations. G1 trees were about 100 years old, whereas G2 trees were between 14 to 26 years old. Estimating the genetic variance across the two generations assumes that genetic values at the different ages of given genotypes are the same or at least strongly correlated. There is however experimental evidence that age-age genetic correlation decrease when the time lag increases, due to different gene or gene effects acting at different ages (Kremer, 1992; Rweyongeza, 2016). Thus developmental related issues should also be addressed when estimating genetic variances of traits especially for long living species (Le Rouzic et al., 2013; Pigliucci, 2008). Combining phenotypic information over successive generations - at different ages - should therefore be used with cautious, or limited to traits for which age-age genetic correlations are known.

### 4.3 Large confidence intervals

Our results overall show large confidence intervals, which may in part also contribute to the variation of the estimates themselves obtained with the different methods. Confidence intervals obtained in our case by bootstrapping can be compared with theoretical expectations derived under more simplified population sampling schemes. The sampling variance of heritability estimates in a population with only distantly related individuals (G1 in our case) was shown to depend mainly on sample size and the variance of relatedness, and less to the population value of heritability itself (Vinkhuyzen et al., 2013; Visscher and Goddard, 2015). Using their method and based on the observed variance of relatedness in our oak population (Appendix 4 and Lesur et al. (2018)), we estimated the confidence intervals to amount to between 0.4 to 0.6 at each side of the estimated value of heritability. This covers almost the entire range of heritability as we observed using bootstrap estimations (Figure 5 and Tables 2 and 3). In sib group designs, sampling variances of heritability are inversely proportional to the number of families and to a lesser extent to the number of offspring per family, and depend also on the population value of heritability as well (Visscher and Goddard, 2015). Using their analytical methods in case of half sib families applied to our sampling sizes in G2, predicted standard deviation of heritability amounted to 0.24 in the *in situ* setting, with only slight variation according to the true population heritability value. Predicted standard deviation for the *ex situ* case varied between 0.05 (when heritability = 0.1) to 0.21 (when heritability = 0.28). These figures using simplified assumptions of half sib families in G2 support both the large and variable sampling variances that we have obtained with the boostrapping. They also confirm that the *ex situ* confidence intervals are expected to be lower than the *in situ* estimates given our sampling sizes imposed by the parentage analysis in the *in situ* case (Figure 1). However using more complete relatedness (***H*** versus ***G*** and versus ***A*** matrix) did not improve the precision of heritability (Figure 4) nor was it the case when more SNPs were used to assess genomic relatedness, as was also found in other cases (Perrier et al., 2018; Bérénos et al., 2014; Stanton-geddes et al., 2013). The ultimate way of improving the precision of estimation is finally to increase sample sizes, especially if one generation estimations are foreseen. While phenotypic assessments of long lived species as trees are labour demanding, current plans for estimating heritability *in situ* are to use long term genetic plots as the ISPs (Intensive Study Plots) benefiting from earlier efforts conducted for recording adaptive related traits as reproduction, growth or phenology (Gerber et al., 2014). Sample sizes of adult trees in ISPs amount usually to a few hundreds, which are too limited for accurate estimates of genetic variances. Hence these plots should be extended either spatially (increasing the number of trees) or temporally (by adding the next generation). Indeed adding one generation, not only improves precision, but also reduces bias.

### 4.4 Variation of heritability and evolvability among trait categories and species

Despite the low statistical power due to limited sample sizes, and despite considering the spatial effect as random, we found important genetic variation for multiple traits investigated regardless of the biological function to which they contribute (Table 2 and 3 and Appendix 5). While genetic variances exhibit variation among traits within a given functional group, there are some noticeable trends across groups. Most phenological traits in the two species show moderate to high heritability values. Growth, reproduction and physiological related traits exhibit intermediate levels of genetic variation despite the high heritability values for some of them, while most leaf morphological traits and secondary metabolites show low heritability values. These patterns confirm earlier results derived from metanalysis conducted in progeny tests of trees (Cornelius, 1994), but contradict theoretical prediction that fitness related traits (here growth, reproduction and phenology) would show lower genetic variation than morphological traits (Weigensberg and Roff, 1996; Price and Schluter, 1991). While difference in heritability values may also be interpreted as differences in environmental variance, evolvability is a standardized additive genetic variance and is therefore free from the environmental variance contribution and allows therefore also comparison with other species and traits (Hansen et al., 2011). With a very few exceptions evolvability values observed in *Q. petraea* and *Q. robur* (Table 2 and 3) are above the mean values of reported studies across plants and animals (Figure 5 in Hansen et al. (2011)), thus suggesting large genetic variation residing within oak species. Following the lack of correlation between evolvability and heritability, our results indicate that there is still substantial genetic variation in fitness related traits as growth and reproduction-therefore offering support for future evolution. Finally, heritability and evolvability values show similar levels in the two species with the noticeable exception of fitness related traits (growth and reproduction mainly) that show opposite trends in both species. Indeed heritability (and evolvability) are much higher for height, diameter and reproductive success in *Q. petraea* than in *Q. robur*. Similarly one can also notice that genetic variation is on average higher in *Q. petraea* for species diagnostic traits either morphological (BS,HR,NV and NL Kremer et al. (2002)) or biochemical (CWSK Prida et al. (2006)). However for physiological traits (*δ*^13^*C* which is also species diagnostic Brendel et al. (2008)) heritability is higher for *Q. robur* on the first generation and for *Q. petraea* on the second. Interestingly these species show different demographic dynamics that have been interpreted as different responses to ongoing natural selection driven by climate change (Truffaut et al., 2017). These selection pressures may have eroded more drastically genetic variation in *Q. robur* that is currently shrinking demographically in the same forest (Truffaut et al., 2017).

### 4.5 Genetic correlations

We found significant genetic correlation between traits within functional categories (Appendix 7) which may be due to either statistical, developmental, structural, functional or evolutionary correlations (Armbruster et al., 2014). With a very few exceptions, these correlations are maintained in the two species. For example, circumference, ring width and height of trees are highly correlated due to their allometric structural relationships. Similarly allometric relationships may contribute as well to the genetic correlation among leaf size related morphological traits. Evolutionary correlations may be noticeable for leaf traits contributing to species differentiation. Indeed petiole length (PL), pubescence (HR), number of intercalary veins (NV) and lamina basal shape (BS), which have been recognized as species diagnostic traits (Kremer et al., 2002), are correlated in both species. However one pair of diagnostic trait exhibits genetic correlation with opposite signs in both species (HR/PL) thus suggesting different evolutionary trait changes within each species. Correlation between leaf unfolding and timing of flowering is most likely driven by developmental integration, while leaf senescence is not correlated with spring phenology confirming earlier observations in *Q. petraea* (Firmat et al., 2017). Functional relationships underlie the correlations among physiological traits investigated in our study, which are mostly involved in water metabolism. Secondary metabolites segregate in two groups of compounds (VNL, CNFL, SYRG, PNTL, 2PHL) and (CSTG, VSCL, VSCG) exhibiting high genetic correlations within groups and low but significant correlations among the two groups. These results suggest that the two groups are likely involved in different biochemical pathways, triggering resistance or susceptibility to herbivory (Carmona et al., 2011).

Overall traits belonging to different functional categories are poorly correlated across categories (Figure7). For example, growth traits were only slightly correlated to bud phenology, and to wood density, and only seldomly to physiological traits. These results may indicate that different trait categories may evolve independently with slight genetic constraints imposed by correlation among different functional categories. However, few traits from different categories were genetically correlated, as *δ*^13^*C* and LUs, for which QTLs have been detected on the same linkage group (Brendel et al., 2008, C. Bodénès pers. comm.). However, in *Q. robur* the correlation is significant in the second generation only, and in *Q. petraea* it is significant in the first generation only.

We observed changes of the sign of correlation between generations which could be either due to ontogenic differences in the development stage at which measures were done on G1 and G2 or to genetic changes over generations. Trees of G2 were far younger than G1 trees when they were analysed, thus the difference in correlation between leaf unfolding and height between generations in *Q. robur* could be due to ontogenic differences in traits expression. Similarly, the difference in correlation sign between *δ*^13^*C* and *δ*^15^*N* can be due to a difference in isotopic composition between old and young trees.

### 4.6 Concluding remarks for future *in situ* estimation of genetic variances in trees

In this study we explored different ways for estimating evolutionary important genetic parameters *in situ* in an intensively studied oak forest. We identified limits for implementing the promising method consisting in assessing genomic relatedness in an extant stand, given the overall low variance of genetic relatedness and the rather low sampling sizes of currently used long term genetic plots in forestry. These limits can be overcome if larger sample sizes are considered, or if the approach is extended over the next generation. Overlapping age structured cohorts are often present in forest stands, and therefore allow to deepen the genetic relatedness among trees. Depending on the resources available, increasing the completeness of relatedness can be achieved by either doing a parentage analysis over two cohorts (as we did in our study) or by retrieving genomic relatedness among all sampled trees in the two cohorts. As shown in our example, parentage reconstruction resulted in larger variances of genetic relatedness as half sib and full sib progenies were identified, while half sib and full sib relationships were extremely rare in the adult cohort. This is an almost trivial outcome of the different demographic structures in an adult and juvenile cohort. Juvenile cohorts live in higher densities and thus maintain sib relationship while adult cohorts are less dense and have lost sib-sib relationships. The two generation strategy may further benefit from the additional phenotypic data recorded in the two cohorts. However this approach should be limited to traits that do not show genetically controlled developmental changes. To conclude, we anticipate that the extension of sample sizes and multiple generations assessments will enhance approaches for estimating genetic variances in forest ecosystems.

## 5 Acknowledgments

This research was supported by the European Research Council through the Advanced Grant Project TREEPEACE (no. FP7–339728). We thank the Office National des Forets and its staff in charge of the Petite Charnie Forest for their constant support and technical assistance during the long term research activities conducted at La Petite Charnie. We are grateful to the staff of the State Nursery of Guéméné Penfao for the raising of the seedlings of the progeny tests and grafting of the trees for the conservation collection. We acknowledge the contribution of Roberto Bacilieri, Maria Evangelista Bertocchi, Gaëlle Capdeville, Benjamin Dencausse, Francois Ehrenmann, Frédéric Expert, Edith Guilley, Marilyn Harrou, Thibault Leroy, Yannick Mellerin, Nastasia Merceron, André Perrin, Andrei Prida, Jean-Louis Puech, Cyrille Rathgeber, Patrick Reynet, Daniel Rittié, Guy Roussel, Armel Thöni, as well as the staff of the Experimental Units of INRA Pierroton (UE 0570, INRA, 33612 Cestas, France) for the collection of material, the plantation of the progeny tests, and for the monitoring of the numerous traits at various periods during the last thirty years. Sequencing and Genotyping for the parentage analysis and relatedness estimation was performed by Christophe Boury, Emilie Chancerel, Erwan Guichoux, Lélia Lagache at the Genome-Transcriptome facility at the Functional Genomic Center of Bordeaux. Bioinformatic analysis for SNP calling was done by Isabelle Lesur (Helix Venture, 33700 Merignac, France). UMR BIOGECO is supported by a grant overseen by the French National Research Agency (ANR) as part of the ”Investissements d’Avenir” programme Labex COTE (ANR-10-LABEX45).

## 6 Authors contribution

AK and HA conceived the study. HA conducted the data analysis with the contribution of CF and AK. AD, JML, LT and BM planned the experimental operations, organized collection of material and data, participated to the data management. GN, JMTR, LT, SD, FL contributed to the collection of physiological, growth, resilience and wood density data and their pre analysis. HA and AK wrote the manuscript and all other authors revised the manuscript.

## Supporting information

Appencides

## References

Akesson, M., Bensch, S., Hasselquist, D., Tarka, M., and Hansson, B. (2008). Estimating Heritabilities and Genetic Correlations: Comparing the” Animal Model” with Parent-Offspring Regression Using Data from a Natural Population. PloS one, 3:e1739.

Armbruster, W. S., Pelabon, C., Bolstad, G. H., and Hansen, T. F. (2014). Integrated phenotypes: understanding trait covariation in plants and animals. Philosophical Transactions of the Royal Society B, 369: 20130245.

Bacilieri, R., Ducousso, A., and Kremer, A. (1996). Comparison of morphological characters and molecular markers for the analysis of hybridization in sessile and pedunculate oak. Annales des Sciences Forestieres, 53: 79–91.

Bacilieri, R., Labbe, T., and Kremer, A. (1994). Intraspecific genetic structure in a mixed population of *Quercus petraea* (Matt.) Leibl and *Q. robur* L. Heredity, 73: 130–141.

Bérénos, C., Ellis, P. A., Pilkington, J. G., and Pemberton, J. M. (2014). Estimating quantitative genetic parameters in wild populations: A comparison of pedigree and genomic approaches. Molecular Ecology, 23: 3434–3451.

Bontemps, A., Lefèvre, F., Davi, H., and Oddou-Muratorio, S. (2016). *In situ* marker-based assessment of leaf trait evolutionary potential in a marginal European beech population. Journal of Evolutionary Biology, 29: 514–527.

Brendel, O. (2014). Is the coefficient of variation a valid measure for variability of stable isotope abundances in biological materials? Rapid Communications in Mass Spectrometry, 28: 370–376.

Brendel, O., Le Thiec, D., Scotti-Saintagne, C., Bodérès, C., Kremer, A., and Guehl, J.-M. (2008). Quantitative trait loci controlling water use efficiency and related traits in *Quercus robur* L. Tree Genetics & Genomes, 4: 263–278.

Carmona, D., Lajeunesse, M. J., and Johnson, M. T. J. (2011). Plant traits that predict resistance to herbivores. Functional Ecology, 25: 358–367.

Castellanos, M. C., Gonzalez-Martinez, S. C., and Pausas, J. G. (2015). Field heritability of a plant adaptation to fire in heterogeneous landscapes. Molecular Ecology, 24: 5633–5642.

Charmantier, A., Garant, D., and Kruuk, L. E. (2014). Quantitative genetics in the wild. Oxford university press, New York.

Coltman, D. W. (2005). Testing marker-based estimates of heritability in the wild. Molecular Ecology, 14: 2593–2599.

Contreras, M. A., Affleck, D., and Chung, W. (2011). Forest Ecology and Management Evaluating tree competition indices as predictors of basal area increment in western Montana forests. Forest Ecology and Management, 262: 1939–1949.

Cornelius, J. (1994). Heritabilities and additive genetic coefficients of variation in forest trees. Canadian journal of Forest research, 24: 372–379.

Firmat, C., Delzon, S., Louvet, J., Parmentier, J., and Kremer, A. (2017). Evolutionary dynamics of the leaf phenological cycle in an oak metapopulation along an elevation gradient. Journal of evolutionary biology, 30: 2116–2131.

Folke, C., Carpenter, S., Walker, B., Scheffer, M., Elmqvist, T., Gunderson, L., and Holling, C. S. (2004). Regime shifts, resilience, and biodiversity in ecosystem management. Annual Review of Ecology, Evolution and Systematics, 35: 557–581.

Gauzere, J., Oddou-Muratorio, S., Pichot, C., Lefevre, F., and Klein, E. (2013). Biases in quantitative genetic analyses using open-pollinated progeny tests from natural tree populations. Acta Botanica Gallica, 160: 227–238.

Gerber, S., Chadoeuf, J., Gugerli, F., Lascoux, M., Buiteveld, J., Cottrell, J., Dounavi, A., Fineschi, S., Forrest, L. L., Fogelqvist, J., Goicoechea, P. G., Jensen, J. S., Salvini, D., Vendramin, G. G., and Kremer, A. (2014). High rates of gene flow by pollen and seed in oak populations across Europe. PloS one, 9.

Guilley, E. (2000). La densité du bois de chêne sessile (Quercus petraea Liebl.). PhD thesis, Ecole Nationale Du Génie Rural Des Eaux Et Forêts.

Hadfield, J. D., Wilson, A. J., Garant, D., Sheldon, B. C., and Kruuk, L. E. B. (2010). The Misuse of BLUP in Ecology and Evolution. The American naturalist, 175: 116–125.

Hansen, T. F., Pe, C., and Houle, D. (2011). Heritability is not Evolvability. Evolutionary Biology, 38: 258.

Hegyi, F. (1974). A simulation model for managing jack-pine stands. In Fries, J., editor, *Growth Models for Tree and Stand Simulation*., pages 74–90. Royal College of Forestery, Stockholm.

Kremer, A. (1992). Predictions of age-age correlations of total height based on serial correlations between height increments in maritime pine *(Pinus pinaster* Ait.). Theoretical and Applied Genetics, 85: 152–158.

Kremer, A., Dupouey, J.-L., Deans, J. D., Cottrell, J., Csaikl, U., Finkeldey, R., Espinel, S., Jensen, J., Kleinschmit, J., Van Dam, B., Ducousso, A., Forrest, I., Lopez de Heredia, U., Lowe, A. J., Tutkova, M., Munro, R. C., Steinhoff, S., Badeau, V., Luc, J., Deans, J. D., Cottrell, J., Csaikl, U., Finkeldey, R., Espinel, S., Jensen, J., Kleinschmit, J., Dam, B. V., Ducousso, A., Forrest, I., Heredia, U. L. D., Lowe, A. J., Tutkova, M., Munro, R. C., Steinhoff, S., and Badeau, V. (2002). Leaf morphological differentiation between *Quercus robur* and *Quercus petraea* is stable across western European mixed oak stands. Annals of Forest Science, 59: 777–787.

Kruuk, L. E. B. (2004). Estimating genetic parameters in natural populations using the animal model ’. Philosophical Transactions of the Royal Society B, 359: 873–890.

Kruuk, L. E. B., and Hadfield, J. D. (2007). How to separate genetic and environmental causes of similarity between relatives. Journal of Evolutionary Biology, 20: 1890–1903.

Lagache, L., Klein, E. K., Guichoux, E., and Petit, R. J. (2013). Fine-scale environmental control of hybridization in oaks. Molecular Ecology, 22: 423–436.

Le Rouzic, A., Alvarez-Castro, J., and Hansen, T. F. (2013). The Evolution of Canalization and Evolvability in Stable and Fluctuating Environments. Evolutionary Biology, 40: 317–340.

Lesur, I., Alexandre, H., Boury, C., Chancerel, E., Plomion, C., and Kremer, A. (2018). Development of target sequence capture and estimation of genomic relatedness in a mixed oak stand. Frontiers in plant science, 9.

Lloret, F., Keeling, E. G., and Sala, A. (2011). Components of tree resilience: effects of successive low-growth episodes in old ponderosa pine forests. Oikos, pages 1909–1920.

Lynch, M. and Walsh, B. (1998). Genetics and Analysis of Quantitative Traits. Sinauer Associates, Inc, Sunderland.

Marshall, T., Slate, J., Kruuk, L. E. B., and Pemberton, J. M. (1998). Statistical confidence for likelihood-based paternity inference in natural populations. Molecular Ecology, 7: 639–655.

Morrissey, M. B., Wilson, A. J., Pemberton, J. M., and Ferguson, M. M. (2007). A framework for power and sensitivity analyses for quantitative genetic studies of natural populations, and case studies in Soay sheep *(Ovis aries*). Journal of Evolutionary Biology, 20: 2309–2321.

Mousseau, T. A., Ritland, K., and Heath, D. D. (1998). A novel method for estimating heritability using molecular markers. Heredity, 80: 218–224.

Muñoz, F. and Sanchez, L. (2018). breedR: Statistical Methods for Forest Genetic Resources Analysts.

Pemberton, J. (2008). Wild pedigrees: the way forward. Proceedings of the Royal Society B: Biological Sciences, 275: 613–621.

Perrier, C., Delahaie, B., and Charmantier, A. (2018). Heritability estimates from genomewide relatedness matrices in wild populations: Application to a passerine, using a small sample size. Molecular Ecology Resources, 18: 838–853.

Pigliucci, M. (2008). Is evolvability evolvable? Nature Reviews Genetics, 9: 75–82.

Polge, H. and Nicholls, J. (1972). Quantitative radiography and the densitometric analysis of wood. Wood Science, 5: 51–59.

Ponzoni, R. and James, J. (1978). Possible Biases in Heritability Estimates from Intraclass Correlation. Theoretical and Applied Genetics, 53: 25–27.

Price, T. D., and Schluter, D. (1991). On the low heritability of life-history traits. Evolution, 45: 853–861.

Prida, A., Boulet, J. C., Ducousso, A., Nepveu, G., and Puech, J. L. (2006). Effect of Species and Ecological Conditions on Ellagitannin Content in Oak Wood From an Even-Aged and Mixed Stand of *Quercus robur* L. And *Quercus petraea* Liebl. Annals of Forest Science, 63: 415–424.

Prida, A., Ducousso, A., Petit, R. J., Nepveu, G., and Puech, J.-L. (2007). Variation in wood volatile compounds in a mixed oak stand: strong species and spatial differentiation in whisky-lactone content. Annals of Forest Science, 64: 313–320.

Ratcliffe, B., Gamal El-Dien, O., Cappa, E. P., Porth, I., Klapste, J., Chen, C., and El-Kassaby, Y. (2017). Single-Step BLUP with Varying Genotyping Effort in Open-Pollinated *Picea glauca*. Genes, Genomes, Genetics, 7: 935–942.

Ritland, K. (2000). Marker-inferred relatedness as a tool for detecting heritability in nature. Molecular Ecology, 9: 1195–1204.

Robinson, M. R., Santure, A. W., Decauwer, I., Sheldon, B. C., and Slate, J. (2013). Partitioning of genetic variation across the genome using multimarker methods in a wild bird population. Molecular Ecology, 22: 3963–3980.

Rweyongeza, D. M. (2016). A new approach to prediction of the age-age correlation for use in tree breeding. Annals of Forest Science, 73: 1099–1111.

Schweiger, R., Kaufman, S., Laaksonen, R., Kleber, M. E., Marz, W., Eskin, E., Rosset, S., and Halperin, E. (2016). Fast and Accurate Construction of Confidence Intervals for Heritability. The American Journal of Human Genetics, 98: 1181–1192.

Stanton-geddes, J., Yoder, J. B., Briskine, R., Young, N. D., and Tiffin, P. (2013). Estimating heritability using genomic data. Methods in Ecology and Evolution, 4: 1151–1158.

Streiff, R., Labbe, T., Bacilieri, R., Steinkellner, H., Gloessl, J., and Krem (1998). Within-population genetic structure in *Quercus robur* L. and *Quercus petraea* (Matt.) Liebl. assessed with isozymes and microsatellites. Molecular Ecology, 7: 317–328.

Torres-Ruiz, J., Kremer, A., Carins, M., Brodribb, T., Larmarque, L., Ducousso, A., and Delzon, S. (submitted). Genetic differentiations in functional traits of European sessile oak populations growing in a common garden.

Truffaut, L., Chancerel, E., Ducousso, A., Dupouey, J. L., Badeau, V., Ehrenmann, F., and Kremer, A. (2017). Fine-scale species distribution changes in a mixed oak stand over two successive generations. New Phytologist, 215: 126–139.

Tumlinson, J. H. (2014). The Importance of Volatile Organic Compounds in Ecosystem Functioning. Journal of chemical ecology, 40: 212–213.

VanRaden, P. M. (2008). Efficient methods to compute genomic predictions. Journal of dairy science, 91: 4414–4423.

Villemereuil, P. D. (2018). Quantitative genetic methods depending on the nature of the phenotypic trait. Annals of the New York Academy of Sciences, pages 1–19.

Vinkhuyzen, A. A., Wray, N. R., Yang, J., Goddard, M. E., and Visscher, P. M. (2013). Estimation and Partition of Heritability in Human Populations Using Whole-Genome Analysis Methods. Annual Review of Genetics, 47: 75–95.

Visscher, P. M. (2008). Sizing up human height variation. Nature Genetics, 40: 489–490.

Visscher, P. M., and Goddard, M. E. (2015). A general unified framework to assess the sampling variance of heritability estimates using pedigree or marker-based relationships. Genetics, 199: 223–232.

Visscher, P. M., Hill, W. G., and Wray, N. R. (2008). Heritability in the genomics era - concepts and misconceptions. Nature Reviews Genetics, 9: 255–266.

Vitasse, Y., Delzon, S., and Dufre, E. (2009). Leaf phenology sensitivity to temperature in European trees: Do within-species populations exhibit similar responses ? Agricultural and forest meteorology, 149: 735–744.

Weigensberg, I. and Roff, D. A. (1996). Natural Heritabilities: Can They be Reliably Estimated in the Laboratory ? Evolution, 50: 2149–2157.

Wilson, A. J. (2008). Why h2 does not always equal V A / V P? Journal of evolutionary biology, 21: 647–650.

