## Supplementary material for "*In situ* estimation of genetic variation of functional and ecological traits in *Quercus petraea* and *Q.robur*": Appencides

### Appendix 1

### Study populations and pedigree relationship between G2 and G1 in the in situ setting

The study population consists in a mixed oak stand (*Q. petraea* and *Q. robur*) of natural origin located is the Petite Charnie State Forest (Latitude: 48.086 N; Longitude: 0.168 W) located in the North West of France. This stands is part of a long term experiment aiming at monitoring ecological and evolutionary processes in oak forests. Our study entails now two successive generations. To sum up the history of the investigations, the long term study started when a seed cut was practiced in 1989 leaving on ground 422 trees (196 *Q. petraea* and 226 *Q. robur*), about 90 to 100 years old.

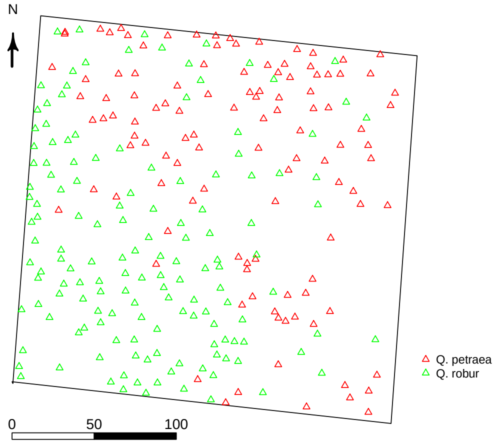


Figure 1.1: Geographic position over the forest stand of G1 trees. *Q. petraea* individuals are in red and *Q. robur* in green, scale represents distance in meters.

The seed cut was followed by two additional thinnings and a final cut was practiced in 1998-2001. The adult trees composing the stand from 1989 to 2001 correspond to our generation G1 (identified as cohort 1 in Truffaut et al. (2017)). The final clear cut of the 298 remaining trees was done over three years (1999, 2000 and 2001) to ease the harvest and manipulation of log samples for later wood and anatomical assessments. In between 1989 and 2001, as the canopy was opened by successive thinnings, seeds originating from open pollination among the remaining adult trees germinated and developed into saplings. Recruitment was quite successful resulting into a total number of about 11 000 saplings in *Q. robur* and 30 000 in *Q. petraea* (Truffaut et al. (2017), table 3) as assessed in 2014 when a demographic inventory was achieved. This sapling population constitutes generation G2 (cohort 2 in Truffaut et al. (2017)), which was 14 to 26 years old when our investigation started in summer 2014. At that time, a sampling of 2510 saplings was made in G2 corresponding to the systematic collection of one sapling every 3 to 6 m.

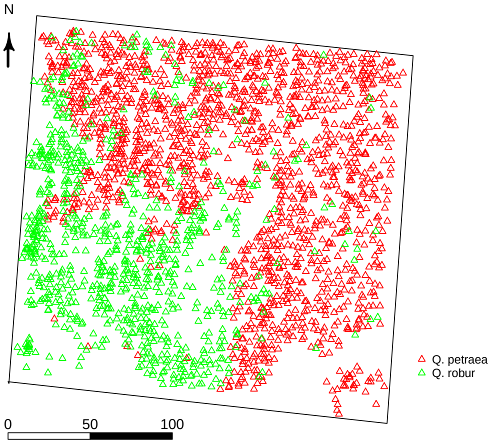


Figure 1.2: Geographic position over the forest stand of each genotyped G2 tree. *Q. petraea* individuals are in red and *Q. robur* in green, scale represents distance in meters.

A parentage analysis was conducted to assign parent offspring relationships between G1 and G2 (Truffaut et al., 2017) based on 80 SNP loci, and using CERVUS v.3.0.7 (Marshall et al., 1998) with stringent parameters, assuming no errors in genotyping (a strict exclusion analysis: 0.0 error rate) and a high confidence level (95%). The parentage analysis showed that a very few number of adult trees did not contribute to the next generation (10 *Q. robur* and 3 *Q. petraea*). For 122 of *Q. petraea* and 193 saplings of *Q. robur* the two parents could be assigned and one parent only for 636 *Q. petraea* saplings and 443 *Q. robur* saplings (Table 1.1). However oaks being monoecious it was not possible to separately assign the male or female parentage.

Based on the parentage analysis we identified a set of saplings for phenotypic assessments in G2 as a sample for estimating heritability *in situ* in G2 (M3 design, Figure 2). This set comprised offspring belonging to all full sib (FS) families detected in G2, e.g. saplings for which the two parents were found within G1, and all half sib families for which the identified parent was also parent of at least one full sib family (Table 1.2).

Table 1.1: Results of the parentage analysis conducted in G2, according to (Truffaut et al., 2017)

| | ***Q. petraea*** | ***Q. robur*** |
|---|---|---|
| **Generation G1** | | |
| Total number of G1 parental trees | 110 | 135 |
| Number of G1 trees which generated at least one offspring | 107 (97.3%) | 125 (92.6%) |
| **Generation G2** | | |
| Total number of G2 saplings sampled | 1570 | 820 |
| Number of offspring assigned to two parents | 122 (7.8%) | 193 (23.5%) |
| Number of offspring assigned to only one parent | 636 (40.5%) | 443 (54.0%) |
| Number of offspring assigned to no parent | 811 (51.7%) | 183 (22.3%) |
| Mean number of offspring per parental tree * | 8.1 | 6.2 |

Table 1.2: Breakdown of sample sizes of full and half sib families used for heritability estimation in G2 (M3 design)

| | ***Q. petraea*** | ***Q. robur*** |
|---|---|---|
| Number of full sib families | 89 | 148 |
| Mean number of offspring/ full sib family | 1.24 | 1.18 |
| Number of half sib families | 80 | 85 |
| Mean number of offspring/ full sib family | 3.24 | 2.54 |
| Number of parents | df | df |
| Total sample size | 370 | 390 |

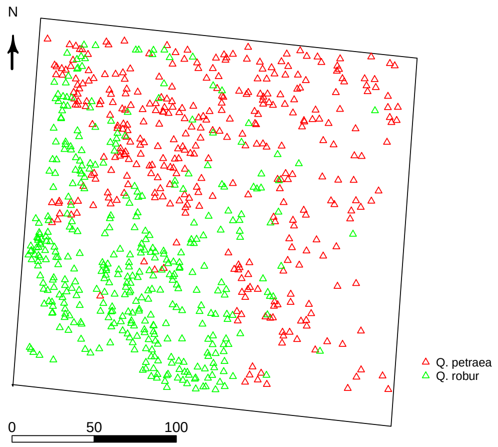


Figure 1.3: Geographic position over the forest stand of each G2 tree phenotyped at least once. *Q. petraea* individuals are in red and *Q. robur* in green, scale represents distance in meters.

### Appendix 2

### Open pollinated progeny test and pedigree relationships of *Quercus petraea* and *Quercus robur* used for *ex situ* estimation of heritability (M6 and M7 method)

In the fall 1995, acorns were collected on 23 *Q. petraea* and 28 *Q. robur* randomly sampled trees of the parental generation (G1) in parcel 26 of the Petite Charnie State Forest. Acorns were sown in the spring 1996 at the State Nursery of Guémééné Penfao (Latitude: 47.6288 N; Longitude: 1.8492 W) located in the North West of France about 150 km West of the Petite Charnie Forest. Two years later in march 1998, seedlings of *Q. petraea* and *Q. robur* were transplanted back in Petite Charnie forest in parcel 37. Parcel 37 is located about 600 meters South of parcel 26 (Figure 2.1).

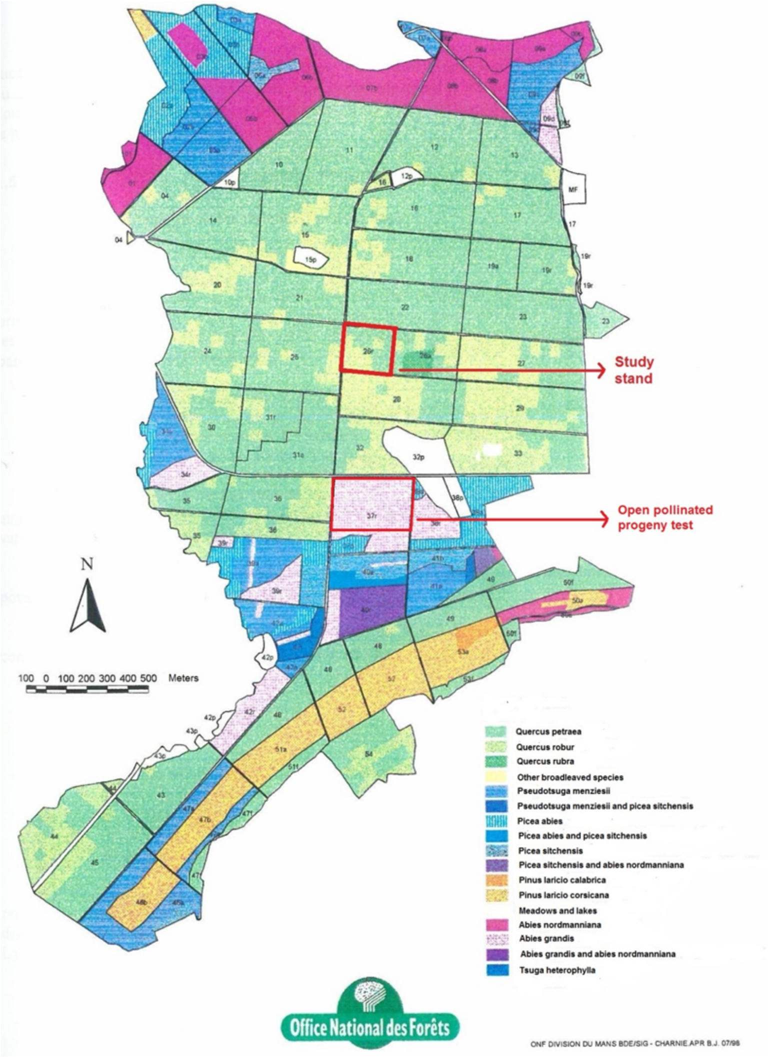


Figure 2.1: Geographic location of the study stand and the open pollinated progeny test

Two separate contiguous progeny trials were installed for *Q. petraea* and *Q. robur* according to a randomized incomplete block design. Each incomplete block comprised 7 open pollinated (OP) progeny plots, and plots comprised 6 OP trees (Table 2.1). When the plantation was made, offsprings of a given progeny were assumed to be half sibs (known female parents, unknown male parents).

In 2009, buds and leaves were harvested on all living trees of the plantation (3512 still living trees, 93% survival) for DNA extraction and genotyping with 12 microsatellites (Lagache et al., 2013, 2014). A parentage analysis, assuming no errors in genotyping (a strict exclusion analysis: 0.0 error rate) and a high confidence level (95%), was implemented to assign parents to the offspring, using CERVUS v.3.0.7 (Marshall et al. (1998), Table 2.2). Male parents were identified for 42% and 43% of the offspring of *Q. petraea* and *Q. robur*, while all female parents were confirmed by parentage analysis (Lagache et al., 2013, 2014).

Additive variances of the different phenotypic traits were estimated on the subset of 641 and 982 *Q. petraea* and *Q. robur* offspring for which male parents of G1 were identified by parentage analysis.

Table 2.1: Description of the experimental design of the two progeny tests

| | ***Q. petraea*** | ***Q. robur*** |
|---|---|---|
| Number of OP families | 23 | 28 |
| Number of blocks | 36 | 54 |
| Number of OP families/block | 7 | 7 |
| Number of trees/plot | 6 | 6 |
| Total number of trees/OP family | 18 to 108 | 18 to 44 |
| Total number of trees | 1512 | 2262 |

Table 2.2: Statistics of the parentage analysis condcuted in the open pollianted progeny tests

| | ***Q. petraea*** | ***Q. robur*** |
|---|---|---|
| Number of offspring with recovered male parent (%) | 641 (42%) | 982 (43%) |
| Number of male parents contributing to the offspring | 78 | 129 |
| Average number of offspring/male parent | 8.23 | 7.61 |
| Number of female parents contributing to the offspring | 23 | 28 |
| Average number of offspring/ female parent | 66 | 81 |

### Appendix 3

### Assessment of resilience components based on ring width analysis Data

Resilience components are based on the response of tree ring width during and after so called ”negative pointer years”, where growth was substantially lower to previous years (Lloret et al., 2011). Ring widths of the 298 trees were measured after the felling of the trees at the final cut. Assessments were made over the whole age of the trees (about 100 years old). 112 trees were felled and measured in 1999, 126 in 2000 and 60 in 2001. Ring width measurements were performed on a stem section cut on the main stem at 1.30 meter from the ground level. Ring width assessments were made later in the lab on 4 radius along the cardinal points. In what follows, we used mean values over the 4 radius.

#### Identification of negative pointer years

To identify negative pointer years we used the method based on relative growth change, initially proposed by Schweingruber et al. (1998). This method compares tree growth in a particular year to the average growth during a given number of preceding years. A calendar year was considered as a negative pointer year when at least 75% of the trees exhibited growth width 20% lower (for negative pointer year or higher for positive pointer year) than the mean of the three previous years. Computations were done within each species, given that the two species *Q.petraea* and *Q.robur* are known to respond differently to the climate (Arend et al., 2013). Admixed trees were not taken into account due to the very low number of individuals. Finally, the computations were done on 121 *Q.petraea* and 164 *Q.robur* for a total number of 285 individuals. Species assignment was based on SNP data and morphological traits (Truffaut et al., 2017). We found 7 negative pointer years in *Q. petraea* and 10 in *Q. robur* (Table 3.1).

#### Estimation of resilience components

Four resilience components were estimated (Lloret et al. (2011); see figure below)

- The component ’resistance’ (inverse of growth reduction during the episode), is conceptually identical to ’abrupt growth changes’ as described in Schweingruber et al. (1998).
- ’Recovery’ (increased growth relative to the minimum growth during the episode) is the ability of tree growth to recover after disturbance.

Table 3.1: Pointer years

| height | ***Q. petraea*** | ***Q. robur*** |
|---|---|---|
| Negative pointer years | 1921 | 1921 |
| | 1932 | 1922 |
| | 1941 | 1932 |
| | 1952 | 1952 |
| | 1954 | 1953 |
| | 1976 | 1954 |
| | 1996 | 1974 |
| | | 1976 |
| | | 1990 |
| | | 1996 |

- 'Resilience' reflects the ability of trees to reach pre-disturbance growth levels (Scheffer et al., 2001; Folke et al., 2004).
- 'Relative resilience' is resilience weighted by the damage incurred during the episode.

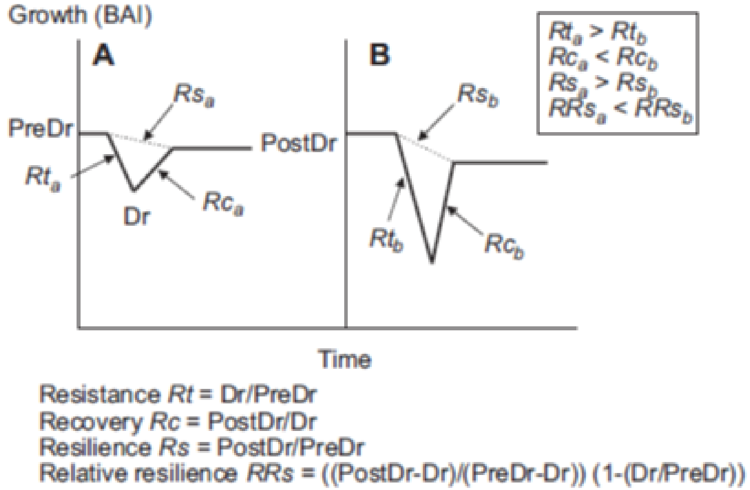


Figure 3.1: Illustration of resilience components according to Lloret et al. (2011)

To estimate components of resilience we used the package pointRes (Van der Maaten-Theunissen et al., 2015) in the R statistical suite (R Core Team, 2016). For the calculation of resilience components, we consider pre-disturbance (PreDr) and post-disturbance (PostDr) period as the average of the ring width 3 years before and after the disturbance. In order to define the lag of the pre or post disturbance period, we calculated the different resilience components by varying the lag from 2 to 5 years. We then considered the correlation matrix between the values obtained for the components in successive years and then decided on the lag period when the matrix exhibited mostly positive values. The final lag of 3 years was also used in other applications of the method (Merlin et al., 2015).

To avoid overlap between the pre- or post-disturbance period and adjacent drought events we decided to use the pre- and post-disturbance period before and after the adjacent events. For instance, the post-disturbance period for 1952 and the pre-disturbance period for 1954 overlap, so we use the 1954 post-disturbance period to calculate the 1952 indices and the 1952 pre-disturbance period for the 1954 indices.

We calculate resilience components on 3 periods corresponding to 3 different stages of tree development:

- Juvenile stage (20-30 years)
- Middle age stage (30-60 years)
- Mature stage (older than 60 years)

For *Q.petraea* (121 trees):

- Resilience components for juvenile stage corresponding to the average of years 1921 and 1932 (except for 3 trees without data on 1921)
- Resilience components for middle age stage corresponding to the average of years 1952 and 1954
- Resilience components for mature stage corresponding to the average of years 1976 and 1996 (except for 33 trees without data in 1996)

For *Q.robur* (164 trees):

- Resilience components for juvenile stage corresponding to the average of years 1921 and 1922 (except for 4 trees without data in 1921)
- Resilience components for middle age stage corresponding to the average of years 1952, 1953 and 1954.
- Resilience components for mature stage corresponding to the average of years 1976 and 1996 (except for 46 trees without data in 1996). We did not take the average between 1990 and 1996 because resistance and resilience indices were not correlated between these two years, whereas for 1976 and 1996 all the indices were positively correlated.

### Appendix 4

### Distribution and variance of genomic relatedness among individuals in both species

In a companion paper, relatedness among all pairs of individuals in the G1 population were calculated for a subset of markers corresponding to different MAF thresholds (Lesur et al., 2018). We report here the variances of relatedness corresponding to the different thresholds together with the resulting number of markers (Table 4.1). We also illustrate the distribution of pairwise relatedness with a close up on the tails with the larger values (Figure 4.1)

Table 4.1: Variance of genomic relatedness within each species depending on the MAF threshold used to select markers. var(G) is the variance of relatedness and n is the number of markers corresponding to each MAF threshold.

| MAF threshold | ***Q. petraea*** | | ***Q. robur*** | |
|---|---|---|---|---|
| | var(G) | n | var(G) | n |
| 1% | 0.002247 | 32047 | 0.000978 | 33131 |
| 5% | 0.002455 | 15274 | 0.001067 | 16408 |
| 10% | 0.002532 | 9502 | 0.001143 | 10225 |
| 15% | 0.002613 | 6753 | 0.001211 | 7143 |
| 30% | 0.002870 | 2849 | 0.001447 | 3058 |
| 40% | 0.003286 | 1454 | 0.001825 | 1561 |

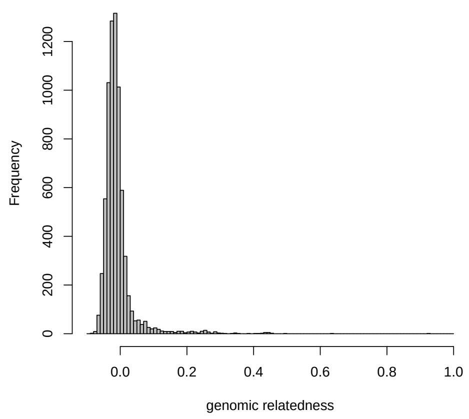


(a) *Q. petraea*

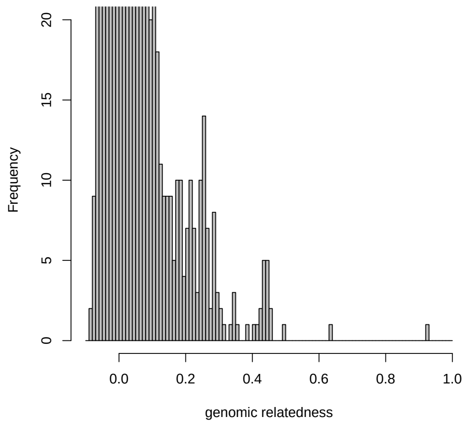


(b) *Q. petraea* zoom in

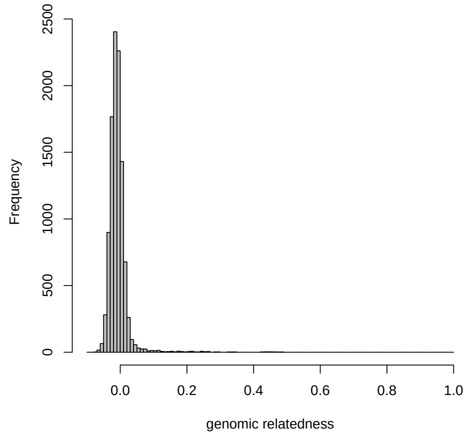


(c) *Q. robur*

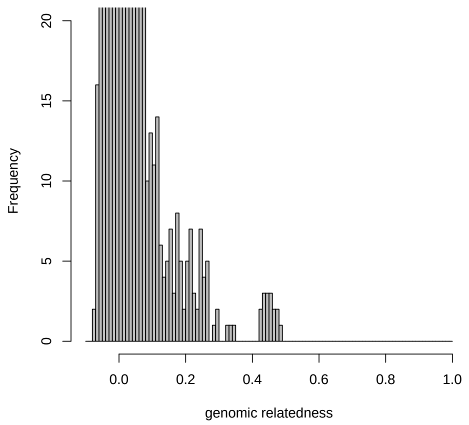


(d) *Q. robur* zoom in

Figure 4.1: Distribution of genomic relatedness computed with SNPs selected at MAF>5% for *Q. petraea* and *Q. robur*. Figures (a) and (c) are the distribution of relatedness of all individuals whereas Figures (b) and (d) are the right tails of the distribution (zoomed for classes with less that 20 occurences).

#### Appendix 5

### Summary of Variance decomposition in both species

Table 5.1: Variance estimates for *Q. petraea*. mth : the method as described in main text Figure 2 with which the trait was analysed; $\mu$: mean of the model fitted values; $V_p$ : observed inter-individual phenotypic variance; $V_a$: additive genetic variance; $V_{sp}$: variance of the spatial effects, when included in the model; $h^2_{obs}$: Heritability computed with the observed phenotypic variance; $h^2_{calc}$: Heritability computed with the phenotypic variance estimated from the model; $I_a$: Evolvability; spt : $\Delta AIC$ between full model and model without spatial effect (a positive value represent a better fit of the model with a spatial effect); add: $\Delta AIC$ between full model and model without additive effect (a positive value represent a better fit of the model with an additive effect). (exp) stands for traits with an exponential distribution. Numbers in brackets represent the 95% CI obtained from 1000 simulation bootstraps.

| trait | mth | $\mu$ | $V_p$ | $V_a$ | $V_{sp}$ | $h^2_{obs}$ | $h^2_{calc}$ | $I_a$ | spt | add |
|---|---|---|---|---|---|---|---|---|---|---|
| Growth | | | | | | | | | | |
| **CIRC** | **M1** | **190.8261** | **948.387** | **734.5 [5.1396 - 1011.15]** | **60.77 [0.6089 - 275.905]** | **0.7745 [0.006 - 0.9371]** | **0.8455 [0.0059 - 0.9831]** | **0.0202 [1e-04 - 0.0279]** | **-1.5874** | **2.3397** |
| CIRC | M3 | 33.5739 | 164.7723 | 21.09 [0.2647 - 71.5678] | 10.19 [0.0906 - 29.1438] | 0.128 [0.0017 - 0.4289] | 0.1306 [0.0017 - 0.4229] | 0.0187 [2e-04 - 0.0625] | 2.2369 | -1.3922 |
| CIRC | M4 | 33.5739 | 164.7723 | 19.45 [0.3496 - 73.7567] | 10.6 [0.1078 - 30.7925] | 0.118 [0.0024 - 0.4548] | 0.1202 [0.0024 - 0.4481] | 0.0173 [3e-04 - 0.0649] | 2.6211 | -1.4067 |
| CIRC | M5 | 70.8102 | 4817.4703 | 4.489 [0.7363 - 63.754] | 20.11 [0.1493 - 57.955] | 9e-04 [0.0022 - 0.1788] | 0.0121 [0.0021 - 0.1736] | 9e-04 [1e-04 - 0.0132] | 2.9095 | -1.9661 |
| **CIRC** | **M6** | **147.8302** | **5340.1252** | **686 [157.4975 - 1472.225]** | **1353 [696.155 - 2130.8]** | **0.1285 [0.0305 - 0.2624]** | **0.1264 [0.0303 - 0.256]** | **0.0314 [0.0072 - 0.0698]** | **-** | **10.994** |
| **CIRC** | **M7** | **147.8302** | **5340.1252** | **713.5 [153.8975 - 1423.075]** | **1342 [716.8925 - 2176.125]** | **0.1336 [0.0285 - 0.2574]** | **0.1314 [0.0279 - 0.2529]** | **0.0326 [0.0069 - 0.0656]** | **-** | **9.9973** |
| **HGHT** | **M1** | **2655.2174** | **24960.9412** | **20950 [149.0975 - 27211]** | **1097 [16.5505 - 5900.75]** | **0.8393 [0.0075 - 0.9426]** | **0.9089 [0.0075 - 0.9845]** | **0.003 [0 - 0.0039]** | **-1.7727** | **0.7856** |
| **HGHT** | **M3** | **1120.307** | **59533.7031** | **14110 [1769.125 - 27491]** | **25090 [9484.975 - 48154.25]** | **0.237 [0.0288 - 0.4772]** | **0.2201 [0.0284 - 0.4411]** | **0.0112 [0.0013 - 0.0241]** | **48.0909** | **3.2197** |
| **HGHT** | **M4** | **1120.307** | **59533.7031** | **17280 [4138.775 - 31803.25]** | **24720 [9035.8 - 45640.25]** | **0.2903 [0.0658 - 0.5666]** | **0.2697 [0.0618 - 0.5162]** | **0.0138 [0.0031 - 0.0271]** | **44.5944** | **5.1395** |
| HGHT | M5 | 1453.5584 | 452521.6554 | 317.3 [141.6525 - 7095.975] | 9744 [2627.625 - 19271.75] | 7e-04 [0.0032 - 0.1515] | 0.0064 [0.0031 - 0.1466] | 2e-04 [1e-04 - 0.0035] | 35.1047 | -2.094 |
| **HGHT** | **M6** | **620.7132** | **36857.5347** | **7321 [2886.7 - 12680]** | **14270 [7944.75 - 23640.25]** | **0.1986 [0.0796 - 0.3303]** | **0.1942 [0.0783 - 0.3255]** | **0.019 [0.0075 - 0.0335]** | **-** | **48.4126** |
| **HGHT** | **M7** | **620.7132** | **36857.5347** | **7748 [2994.225 - 13670.25]** | **14220 [8364.625 - 21672]** | **0.2102 [0.081 - 0.3611]** | **0.205 [0.0809 - 0.3509]** | **0.0201 [0.0074 - 0.0367]** | **-** | **47.3424** |
| **RSURF** | **M1** | **2631.6727** | **748389.0774** | **477600 [12766.75 - 576905]** | **9251 [431.015 - 96566.5]** | **0.6382 [0.0171 - 0.7709]** | **0.3804 [0.0117 - 0.5217]** | **0.069 [0.0019 - 0.0879]** | **-2.0347** | **1.7635** |
| RSURF | M3 | 448.7549 | 92647.5562 | 4882 [1349.225 - 5938.125] | 7260 [2447.5 - 13130.5] | 0.0527 [0.0146 - 0.0641] | 0.0228 [0.01 - 0.0646] | 0.0242 [0.0063 - 0.0457] | 1.6351 | -1.9839 |
| RSURF | M4 | 448.7549 | 92647.5562 | 2724 [530.56 - 3450.125] | 7468 [2676.65 - 12962.25] | 0.0294 [0.0057 - 0.0372] | 0.0127 [0.004 - 0.0352] | 0.0135 [0.0025 - 0.0247] | 2.0346 | -2.087 |
| RSURF | M5 | 1846.6697 | 1600417.138 | 9663 [3700.875 - 10791.75] | 6548 [1619 - 13712.25] | 0.006 [0.0023 - 0.0067] | 0.0021 [8e-04 - 0.0027] | 0.0028 [0.001 - 0.0039] | -0.2673 | -2.1721 |
| **RWDTH** | **M1** | **2.8833** | **0.1973** | **0.1551 [0.0058 - 0.1874]** | **0.0014 [2e-04 - 0.0316]** | **0.786 [0.0295 - 0.9497]** | **0.1914 [0.008 - 0.2462]** | **0.0187 [7e-04 - 0.0234]** | **-2.0624** | **2.6609** |
| RWDTH | M3 | 2.7274 | 0.833 | 0.0196 [0.0021 - 0.0286] | 0.0542 [0.0175 - 0.1046] | 0.0236 [0.0025 - 0.0344] | 0.0167 [0.0046 - 0.0722] | 0.0026 [3e-04 - 0.0041] | 2.8253 | -2.0643 |
| RWDTH | M4 | 2.7274 | 0.833 | 0.0178 [0.0019 - 0.0285] | 0.0544 [0.0171 - 0.1013] | 0.0214 [0.0023 - 0.0342] | 0.0151 [0.0044 - 0.0702] | 0.0024 [3e-04 - 0.0039] | 2.8154 | -2.0566 |
| RWDTH | M5 | 2.8256 | 0.4297 | 0.018 [0.0073 - 0.0227] | 0.0289 [0.0105 - 0.0525] | 0.0418 [0.017 - 0.0528] | 0.013 [0.0092 - 0.0313] | 0.0022 [9e-04 - 0.0029] | 1.4616 | -2.0645 |
| R success | | | | | | | | | | |
| **NOFF** | **M1** | **8.7191** | **81.1361** | **66.83 [0.9449 - 90.22]** | **14.77 [0.1413 - 42.2218]** | **0.8237 [0.011 - 0.9238]** | **0.7829 [0.0107 - 0.977]** | **0.8791 [0.0151 - 2.2972]** | **1.0722** | **0.6427** |
| Phenology | | | | | | | | | | |
| **LUs** | **M2** | **2.4219** | **1.3245** | **1.12 [0.0585 - 1.3872]** | **0.1934 [6e-04 - 0.6535]** | **0.8456 [0.0442 - 1.0474]** | **0.8465 [0.0481 - 0.9825]** | **0.1909 [0.0093 - 0.2966]** | **-** | **16.0096** |
| **LUs** | **M3** | **4.0449** | **1.9337** | **1.038 [0.3551 - 1.7991]** | **0.2248 [0.0194 - 0.5622]** | **0.5368 [0.1836 - 0.9304]** | **0.5392 [0.2011 - 0.8699]** | **0.0634 [0.0216 - 0.116]** | **2.158** | **8.0346** |
| **LUs** | **M4** | **4.0449** | **1.9337** | **0.9667 [0.2603 - 1.72]** | **0.2078 [0.0073 - 0.527]** | **0.4999 [0.1346 - 0.8895]** | **0.5073 [0.1453 - 0.8495]** | **0.0591 [0.0157 - 0.11]** | **2.2363** | **8.1052** |
| **LUs** | **M6** | **2.2461** | **1.142** | **0.3584 [0.2747 - 0.4504]** | **0.2008 [0.103 - 0.3108]** | **0.3138 [0.2405 - 0.3944]** | **0.302 [0.2385 - 0.3695]** | **0.071 [0.0505 - 0.0988]** | **-** | **188.7006** |
| **LUs** | **M7** | **2.2461** | **1.142** | **0.3547 [0.2685 - 0.4477]** | **0.2014 [0.1082 - 0.3244]** | **0.3106 [0.2351 - 0.392]** | **0.2997 [0.2336 - 0.3658]** | **0.0703 [0.0496 - 0.0992]** | **-** | **190.5778** |
| **Lud** | **M2** | **105.5218** | **22.7381** | **19.55 [0.5471 - 24.4108]** | **1.6567 [0.004 - 7.7655]** | **0.8598 [0.026 - 0.9508]** | **0.9138 [0.0241 - 0.9881]** | **-** | **-** | **20.5593** |
| **LS** | **M2** | **305.7786** | **15.3089** | **12.66 [0.3094 - 16.05]** | **1.7025 [0.0063 - 5.8742]** | **0.827 [0.019 - 0.8977]** | **0.8714 [0.0197 - 0.9793]** | **-** | **-** | **3.4352** |
| **GSL** | **M2** | **200.2139** | **34.7263** | **28.38 [0.4559 - 35.8205]** | **4.545 [0.01 - 17.9061]** | **0.8172 [0.0139 - 0.9388]** | **0.8537 [0.0145 - 0.9856]** | | **-** | **10.9387** |
| FFLW | M1 | 123.214 | 47.5872 | 14.03 [0.1188 - 27.056] | 0.2001 [0.0115 - 8.4535] | 0.2948 [0.0025 - 0.5686] | 0.9547 [0.0189 - 0.9786] | - | -2.0127 | -0.7387 |
| **MFLW** | **M1** | **120.1784** | **35.043** | **16.25 [0.2459 - 23.177]** | **0.1015 [0.0123 - 5.5061]** | **0.4637 [0.007 - 0.6614]** | **0.9826 [0.0173 - 0.9841]** | **-** | **-2.0344** | **5.2571** |
| MAR | M1 | 0.7917 | 2.293 | 0.2048 [1e-04 - 0.9147] | 0.0011 [0 - 0.1593] | 0.0893 [1e-04 - 0.9216] | 0.2191 [1e-04 - 0.9634] | 0.3268 [2e-04 - 1.6089] | -2.0465 | -1.8562 |
| Physiology | | | | | | | | | | |
| C | M2 | 455.3283 | 785.3835 | 183.8 [0.0992 - 752.3275] | - | 0.234 [1e-04 - 0.9187] | 0.2357 [1e-04 - 0.9582] | 9e-04 [0 - 0.0036] | - | -1.9429 |
| C | M3 | 459.0698 | 593.9026 | 4.096 [0.2437 - 180.8725] | 35.42 [0.0127 - 101.5075] | 0.0069 [5e-04 - 0.2948] | 0.0067 [5e-04 - 0.2969] | 0 [0 - 9e-04] | 0.5036 | -2.0228 |
| C | M4 | 459.0698 | 593.9026 | 5.062 [0.2049 - 171.8225] | 35.09 [0.0117 - 111.1825] | 0.0085 [4e-04 - 0.2838] | 0.0083 [4e-04 - 0.2785] | 0 [0 - 8e-04] | 0.2749 | -2.0167 |
| C.N | M2 | 24.3475 | 12.5891 | 1.311 [0.0014 - 11.3413] | - | 0.1041 [1e-04 - 0.9062] | 0.1046 [1e-04 - 0.9447] | 0.0022 [0 - 0.0192] | - | -1.9591 |
| C.N | M3 | 19.3065 | 4.5348 | 0.4714 [0.0171 - 1.9514] | 0.7118 [0.0798 - 1.6911] | 0.104 [0.004 - 0.4212] | 0.0992 [0.0039 - 0.4137] | 0.0013 [0 - 0.0053] | 14.7864 | -1.7479 |
| C.N | M4 | 19.3065 | 4.5348 | 0.5976 [0.0193 - 1.9458] | 0.7137 [0.0866 - 1.717] | 0.1318 [0.0045 - 0.4177] | 0.1256 [0.0044 - 0.4087] | 0.0016 [1e-04 - 0.0052] | 14.29 | -1.6342 |
| d13C | M2 | -29.6116 | 1.304 | 0.0763 [1e-04 - 1.0911] | - | 0.0585 [1e-04 - 0.8374] | 0.0587 [1e-04 - 0.8725] | 1e-04 [0 - 0.0012] | - | -1.9814 |
| **d13C** | **M3** | **-29.5366** | **1.069** | **0.3796 [0.0523 - 0.796]** | **0.1343 [0.0061 - 0.3119]** | **0.3551 [0.0509 - 0.7148]** | **0.3523 [0.0485 - 0.6779]** | **4e-04 [1e-04 - 9e-04]** | **2.7839** | **3.0466** |
| **d13C** | **M4** | **-29.5366** | **1.069** | **0.4632 [0.1123 - 0.8959]** | **0.116 [0.0053 - 0.2967]** | **0.4333 [0.1111 - 0.8432]** | **0.433 [0.1074 - 0.8202]** | **5e-04 [1e-04 - 0.001]** | **1.3399** | **5.2009** |
| **d15N** | **M2** | **-3.3505** | **1.8222** | **1.574 [0.0056 - 2.109]** | **-** | **0.8638 [0.0035 - 0.9487]** | **0.8808 [0.0035 - 0.9886]** | **0.1402 [5e-04 - 0.1922]** | **-** | **0.8272** |
| **d15N** | **M3** | **-6.4981** | **0.8863** | **0.1647 [0.0124 - 0.3822]** | **0.6586 [0.2337 - 1.1541]** | **0.1858 [0.0124 - 0.4227]** | **0.1402 [0.0112 - 0.3761]** | **0.0039 [3e-04 - 0.0096]** | **60.2578** | **0.4112** |
| d15N | M4 | -6.4981 | 0.8863 | 0.1056 [0.0077 - 0.3097] | 0.6522 [0.2844 - 1.1511] | 0.1191 [0.0079 - 0.3077] | 0.0906 [0.0072 - 0.2692] | 0.0025 [2e-04 - 0.0076] | 60.6673 | -0.9627 |
| N | M2 | 19.0384 | 6.3556 | 0.8565 [7e-04 - 5.8692] | - | 0.1348 [1e-04 - 0.9018] | 0.1355 [1e-04 - 0.9449] | 0.0024 [0 - 0.0163] | - | -1.9491 |
| N | M3 | 24.0214 | 6.9429 | 0.2496 [0.0021 - 2.2862] | 0.379 [1e-04 - 1.1774] | 0.036 [3e-04 - 0.3107] | 0.0353 [3e-04 - 0.3079] | 4e-04 [0 - 0.004] | 2.4782 | -1.9535 |

| | | | | | | | | | | |
|---|---|---|---|---|---|---|---|---|---|---|
| N | M4 | 24.0214 | 6.9429 | 0.6083 [0.009 - 2.7276] | 0.3926 [0.0015 - 1.206] | 0.0876 [0.0014 - 0.3909] | 0.0857 [0.0014 - 0.3872] | 0.0011 [0 - 0.0047] | 2.5239 | -1.7577 |
| SLA | M2 | 11.719 | 5.2619 | 9e-04 [5e-04 - 4.2129] | - | 2e-04 [1e-04 - 0.8532] | 2e-04 [1e-04 - 0.8677] | 0 [0 - 0.0309] | - | -2.001 |
| **SLA** | **M3** | **13.3166** | **12.0123** | **4.154 [0.2895 - 8.9597]** | **1.107 [0.0411 - 2.9817]** | **0.3458 [0.0239 - 0.7163]** | **0.3365 [0.0229 - 0.6885]** | **0.0234 [0.0016 - 0.0499]** | **4.8645** | **1.4142** |
| **SLA** | **M4** | **13.3166** | **12.0123** | **3.673 [0.2566 - 8.625]** | **1.094 [0.0219 - 2.8908]** | **0.3058 [0.0224 - 0.671]** | **0.2983 [0.0208 - 0.6616]** | **0.0207 [0.0015 - 0.0482]** | **4.1966** | **1.0039** |
| MLA | M2 | 44.801 | 111.6555 | 53.39 [0.0182 - 117.325] | - | 0.4782 [2e-04 - 0.9333] | 0.4869 [2e-04 - 0.982] | 0.0266 [0 - 0.0599] | - | -0.778 |
| **MLA** | **M3** | **32.0433** | **77.5867** | **20.99 [0.1232 - 51.515]** | **0.0907 [0.0026 - 4.5686]** | **0.2705 [0.0018 - 0.627]** | **0.2696 [0.0018 - 0.6251]** | **0.0204 [1e-04 - 0.049]** | **-2.0696** | **2.081** |
| MLA | M4 | 32.0433 | 77.5867 | 13.56 [0.0248 - 39.502] | 0.1071 [6e-04 - 4.3364] | 0.1748 [3e-04 - 0.5126] | 0.1739 [3e-04 - 0.5042] | 0.0132 [0 - 0.0393] | -2.0616 | -0.348 |
| Resilience | | | | | | | | | | |
| REC | M1 | 1.3393 | 0.0223 | 0 [0 - 0.0065] | 4e-04 [0 - 0.0048] | 0.0015 [1e-04 - 0.2922] | 0.0788 [0.0167 - 0.9561] | 0 [0 - 0.0037] | -1.9307 | -2.0056 |
| REL | M1 | 0.8509 | 0.0103 | 5e-04 [0 - 0.0031] | 0.0014 [0 - 0.0042] | 0.0464 [2e-04 - 0.2983] | 0.2547 [0.0107 - 0.9488] | 7e-04 [0 - 0.0042] | -0.3347 | -1.8761 |
| RET | M1 | 0.6647 | 0.0057 | 5e-04 [0 - 0.002] | 0 [0 - 9e-04] | 0.0854 [1e-04 - 0.354] | 0.9533 [0.0187 - 0.968] | 0.0011 [0 - 0.0047] | -2.0092 | -1.6311 |
| Structure | | | | | | | | | | |
| WD | M1 | 576.7698 | 886.685 | 417.4 [2.9645 - 875.3225] | 173.6 [1.667 - 477.715] | 0.4707 [0.0038 - 0.8906] | 0.448 [0.0038 - 0.9222] | 0.0013 [0 - 0.0027] | -0.959 | -0.6863 |
| **WD** | **M3** | **507.0777** | **1121.0502** | **443.2 [112.745 - 807.985]** | **268.5 [80.4813 - 559.4675]** | **0.3953 [0.1102 - 0.7261]** | **0.3883 [0.1077 - 0.6926]** | **0.0017 [4e-04 - 0.0032]** | **32.9723** | **5.0942** |
| **WD** | **M4** | **507.0777** | **1121.0502** | **492.1 [143.2675 - 838.0275]** | **255.5 [82.5705 - 527.9175]** | **0.439 [0.1317 - 0.7554]** | **0.43 [0.1357 - 0.7185]** | **0.0019 [5e-04 - 0.0033]** | **30.4821** | **3.9837** |
| **WD** | **M5** | **523.6702** | **1935.5673** | **481.8 [228.4975 - 760.4025]** | **-** | **0.2489 [0.2394 - 0.6761]** | **0.4577 [0.2398 - 0.672]** | **0.0018 [8e-04 - 0.0028]** | **-** | **29.1774** |
| Leaf morphology | | | | | | | | | | |
| **BS** | **M1** | **3.9635** | **0.6307** | **0.4317 [0.0072 - 0.572]** | **0.0418 [4e-04 - 0.1781]** | **0.6844 [0.0114 - 0.9068]** | **0.9031 [0.0161 - 0.9853]** | **0.0275 [4e-04 - 0.0369]** | **-1.5294** | **6.0746** |
| HR | M1 | 3.3708 | 0.7707 | 0.0685 [8e-04 - 0.1128] | 0.0018 [1e-04 - 0.0368] | 0.0889 [0.0011 - 0.1464] | 0.117 [0.0181 - 0.9782] | 0.006 [1e-04 - 0.01] | -2.0416 | -1.8682 |
| LDR | M1 | 26.2344 | 31.5929 | 2.39 [0.0539 - 5.2668] | 5.585 [0.7704 - 12.4017] | 0.0757 [0.0017 - 0.1667] | 0.095 [0.0083 - 0.7282] | 0.0035 [1e-04 - 0.0076] | -0.4375 | -1.978 |
| LL | M1 | 92.5365 | 310.0745 | 50.02 [0.868 - 72.99] | 25.86 [0.3115 - 64.003] | 0.1613 [0.0028 - 0.2354] | 0.1995 [0.0128 - 0.9459] | 0.0058 [1e-04 - 0.0087] | -1.107 | -1.7996 |
| LW | M1 | 28.4354 | 40.2459 | 2.989 [0.0603 - 5.0841] | 3.027 [0.0595 - 7.4134] | 0.0743 [0.0015 - 0.1263] | 0.0841 [0.0107 - 0.8963] | 0.0037 [1e-04 - 0.0063] | -1.2125 | -1.9517 |
| LWR | M1 | 30.5788 | 11.3129 | 5.039 [0.0837 - 6.6942] | 0.0272 [0.0034 - 1.2253] | 0.4454 [0.0074 - 0.5917] | 0.5172 [0.0165 - 0.9874] | 0.0054 [1e-04 - 0.0071] | -2.0678 | -0.7868 |
| **NL** | **M1** | **11.2051** | **2.3663** | **1.846 [0.0355 - 2.3403]** | **0.0162 [0.0011 - 0.4235]** | **0.7801 [0.015 - 0.989]** | **0.978 [0.0209 - 0.9879]** | **0.0147 [3e-04 - 0.019]** | **-2.0121** | **2.7978** |
| **NV** | **M1** | **0.323** | **0.2549** | **0.1939 [0.0029 - 0.2566]** | **9e-04 [1e-04 - 0.045]** | **0.7607 [0.0113 - 1.0068]** | **0.9647 [0.0166 - 0.987]** | **1.8582 [0.0284 - 3.2702]** | **-2.0185** | **0.2139** |
| OB | M1 | 54.2075 | 21.6011 | 2.206 [0.0542 - 6.5899] | 7.361 [0.4659 - 17.3665] | 0.1021 [0.0025 - 0.3051] | 0.225 [0.0074 - 0.697] | 8e-04 [0 - 0.0023] | 2.5572 | -1.7428 |
| PL | M1 | 16.9551 | 25.8648 | 0.2235 [0.0107 - 0.9484] | 2.43 [0.5555 - 4.919] | 0.0086 [4e-04 - 0.0367] | 0.0098 [0.0046 - 0.3983] | 8e-04 [0 - 0.0033] | -0.316 | -2.0532 |
| PR | M1 | 15.2147 | 9.1948 | 0.1273 [1e-04 - 0.407] | 0.074 [0 - 0.3599] | 0.0138 [0 - 0.0443] | 0.0149 [0.0138 - 0.9627] | 5e-04 [0 - 0.0017] | -2.008 | -2.0375 |
| **PV** | **M1** | **3.0765** | **31.5445** | **27.92 [0.5413 - 35.924]** | **0.2392 [0.0172 - 5.6771]** | **0.8851 [0.0172 - 1.1388]** | **0.9762 [0.0198 - 0.9877]** | **2.95 [0.053 - 5.4005]** | **-2.0164** | **1.1456** |
| SW | M1 | 20.6742 | 20.674 | 0.3935 [0.0081 - 1.1194] | 1.445 [0.0225 - 3.271] | 0.019 [4e-04 - 0.0541] | 0.0209 [0.0068 - 0.7697] | 9e-04 [0 - 0.0026] | -1.2963 | -2.0273 |
| WP | M1 | 50.3258 | 120.6494 | 55.56 [1.0239 - 76.1213] | 9.1 [0.0703 - 31.2218] | 0.4605 [0.0085 - 0.6309] | 0.6686 [0.0157 - 0.9824] | 0.0219 [4e-04 - 0.0302] | -1.2919 | -0.7932 |
| Defence | | | | | | | | | | |
| CNFL (exp) | M1 | 0.7782 | 3.8295 | 0.0048 [2e-04 - 0.2036] | 0.0584 [0 - 0.1622] | 0.0121 [7e-04 - 0.6977] | 0.0157 [7e-04 - 0.7051] | 0.0048 [2e-04 - 0.2036] | 2.564 | -2.01 |
| CSTG (exp) | M1 | 3.5445 | 23.8251 | 1e-04 [0 - 0.0147] | 0.0016 [0 - 0.0064] | 0.0034 [2e-04 - 0.8427] | 0.0036 [2e-04 - 0.8572] | 1e-04 [0 - 0.0147] | -1.3323 | -2.0219 |
| CSTL (exp) | M1 | 0.275 | 1.1465 | 1e-04 [0 - 0.2349] | 0 [0 - 0.0535] | 3e-04 [1e-04 - 0.7069] | 3e-04 [1e-04 - 0.7276] | 1e-04 [0 - 0.2349] | -2.0006 | -2.0009 |
| **CWSK (exp)** | **M1** | **1.0311** | **54.4825** | **1.529 [0.001 - 2.7502]** | **0.0084 [1e-04 - 0.479]** | **0.1219 [1e-04 - 0.2756]** | **0.1493 [1e-04 - 0.3011]** | **1.529 [0.001 - 2.7502]** | **-2.0604** | **0.8847** |
| EGNL (exp) | M1 | -1.2847 | 0.4896 | 0.0305 [1e-04 - 1.0222] | 5e-04 [0 - 0.2006] | 0.0084 [1e-04 - 0.4794] | 0.0146 [1e-04 - 0.4979] | 0.0305 [1e-04 - 1.0222] | -2.0068 | -2.0111 |
| ELAC (exp) | M1 | 0.6114 | 1.4765 | 0.1354 [1e-04 - 0.2986] | 7e-04 [0 - 0.0433] | 0.4049 [2e-04 - 0.8256] | 0.4476 [2e-04 - 0.8673] | 0.1354 [1e-04 - 0.2986] | -2.0622 | -0.0089 |
| ELTOT (exp) | M1 | 3.4654 | 261.8303 | 1e-04 [0 - 0.1629] | 0 [0 - 0.034] | 3e-04 [1e-04 - 0.7706] | 3e-04 [1e-04 - 0.7799] | 1e-04 [0 - 0.1629] | -2.0006 | -2.0009 |
| GRDN | M1 | 8.0904 | 9.1947 | 2.046 [0.0015 - 9.0711] | 0.0203 [2e-04 - 1.6156] | 0.2225 [2e-04 - 0.9266] | 0.2222 [2e-04 - 0.9655] | 0.0313 [0 - 0.1418] | -2.0442 | -1.7102 |
| MVL (exp) | M1 | -0.1098 | 0.5417 | 1e-04 [0 - 0.3001] | 1e-04 [0 - 0.0606] | 1e-04 [1e-04 - 0.7283] | 2e-04 [1e-04 - 0.7364] | 1e-04 [0 - 0.3001] | -2.0006 | -2.0009 |
| PNTL (exp) | M1 | -1.6259 | 0.018 | 0.0165 [0 - 0.238] | 0.0141 [0 - 0.0788] | 0.0404 [1e-04 - 0.7852] | 0.056 [1e-04 - 0.804] | 0.0165 [0 - 0.238] | -1.6178 | -1.9896 |
| ROBA (exp) | M1 | 1.5048 | 2.3083 | 0.0102 [4e-04 - 0.0678] | 0.0412 [0.0047 - 0.0958] | 0.0941 [0.0044 - 0.6617] | 0.0914 [0.0043 - 0.6517] | 0.0102 [4e-04 - 0.0678] | 7.0827 | -1.92 |
| ROBB | M1 | 6.8903 | 5.7242 | 0.0498 [0.0023 - 4.3551] | 0.9873 [4e-04 - 2.9472] | 0.0087 [4e-04 - 0.7703] | 0.0082 [4e-04 - 0.765] | 0.001 [0 - 0.0951] | 0.5364 | -2.0164 |
| ROBC (exp) | M1 | 1.7064 | 7.7695 | 0.006 [0 - 0.1215] | 0.0199 [0 - 0.067] | 0.032 [2e-04 - 0.7902] | 0.0369 [2e-04 - 0.8021] | 0.006 [0 - 0.1215] | -0.7547 | -1.9922 |
| ROBD | M1 | 8.6726 | 11.6933 | 1.44 [0.0016 - 11.3305] | 0.0121 [2e-04 - 2.1233] | 0.1231 [1e-04 - 0.9139] | 0.1177 [1e-04 - 0.9456] | 0.0191 [0 - 0.1537] | -2.0222 | -1.9537 |
| ROBE (exp) | M1 | 2.0943 | 5.447 | 9e-04 [1e-04 - 0.0527] | 0.0193 [0 - 0.0526] | 0.0105 [0.0018 - 0.7153] | 0.0109 [0.0018 - 0.7139] | 9e-04 [1e-04 - 0.0527] | 3.8424 | -2.0186 |
| SYRG | M1 | 5.3094 | 6.4986 | 0.4608 [5e-04 - 3.9126] | 0.0144 [1e-04 - 0.7443] | 0.0709 [1e-04 - 0.9207] | 0.1107 [1e-04 - 0.9626] | 0.0163 [0 - 0.1421] | -2.0177 | -1.9701 |
| TWSK (exp) | M1 | 0.0692 | 38.728 | 1.405 [0.0085 - 2.853] | 0.3487 [0.0015 - 1.205] | 0.1016 [6e-04 - 0.2121] | 0.0902 [6e-04 - 0.2317] | 1.405 [0.0085 - 2.853] | -1.4833 | -1.1858 |
| VNL | M1 | 3.0451 | 2.5027 | 0.4337 [5e-04 - 1.8052] | 0.2087 [1e-04 - 0.8111] | 0.1733 [3e-04 - 0.8947] | 0.2257 [3e-04 - 0.9307] | 0.0468 [1e-04 - 0.2088] | -1.4977 | -1.9173 |
| VSCG | M1 | 18.1981 | 23.872 | 0.1925 [0.0024 - 20.232] | 0.0012 [4e-04 - 4.877] | 0.0081 [1e-04 - 0.8931] | 0.0082 [1e-04 - 0.9064] | 6e-04 [0 - 0.0638] | -1.9888 | -2.0011 |
| VSCL (exp) | M1 | 0.6366 | 4.2427 | 0.001 [0 - 0.2957] | 0.0083 [0 - 0.0812] | 0.0016 [1e-04 - 0.7322] | 0.0025 [1e-04 - 0.7693] | 0.001 [0 - 0.2957] | -1.8817 | -2.0073 |
| 2PHL (exp) | M1 | -2.1278 | 0.0072 | 0.0015 [0 - 0.2844] | 0.015 [0 - 0.0877] | 0.0029 [1e-04 - 0.7588] | 0.0039 [1e-04 - 0.7791] | 0.0015 [0 - 0.2844] | -1.858 | -2.0087 |

Table 5.2: Variance estimates for *Q. robur*. mth : the method as described in main text Figure 2 with which the trait was analysed; $\mu$: mean of the model fitted values; $V_p$ : observed inter-individual phenotypic variance; $V_a$: additive genetic variance; $V_{sp}$: variance of the spatial effects, when included in the model; $h^2_{obs}$: Heritability computed with the observed phenotypic variance; $h^2_{calc}$: Heritability computed with the phenotypic variance estimated from the model; $I_a$: Evolvability; spt : $\Delta AIC$ between full model and model without spatial effect (a positive value represent a better fit of the model with a spatial effect); add: $\Delta AIC$ between full model and model without additive effect (a positive value represent a better fit of the model with an additive effect). (exp) stands for traits with an exponential distribution. Numbers in brackets represent the 95% CI obtained from 1000 simulation bootstraps.

| trait | mth | $\mu$ | $V_p$ | $V_a$ | $V_{sp}$ | $h^2_{obs}$ | $h^2_{calc}$ | $I_a$ | spt | add |
|---|---|---|---|---|---|---|---|---|---|---|
| Growth | | | | | | | | | | |
| CIRC | M1 | 167.4394 | 554.8436 | 68.84 [0.881 - 425.6625] | 77.27 [0.0893 - 223.4575] | 0.1241 [0.0019 - 0.7982] | 0.1308 [0.0019 - 0.8438] | 0.0025 [0 - 0.0155] | 0.1687 | -1.9547 |
| CIRC | M3 | 27.386 | 78.7278 | 9.051 [0.2835 - 30.491] | 10.33 [1.2242 - 23.4105] | 0.115 [0.0042 - 0.4097] | 0.1233 [0.0042 - 0.4018] | 0.0121 [4e-04 - 0.0416] | 6.5267 | -0.8949 |
| CIRC | M4 | 27.386 | 78.7278 | 7.941 [0.2766 - 25.9303] | 10.16 [1.5138 - 23.7812] | 0.1009 [0.004 - 0.3657] | 0.1084 [0.0039 - 0.3509] | 0.0106 [3e-04 - 0.0352] | 5.9601 | -1.1618 |
| CIRC | M5 | 69.727 | 4351.2221 | 19.43 [0.6732 - 63.071] | 11.49 [0.1628 - 28.7048] | 0.0045 [0.0032 - 0.2697] | 0.0849 [0.0032 - 0.2693] | 0.004 [1e-04 - 0.0131] | 0.4691 | -0.2909 |
| **CIRC** | **M6** | **159.4237** | **5111.5907** | **827.4 [343.9225 - 1476.1]** | **470.6 [254.28 - 718.0225]** | **0.1619 [0.067 - 0.2757]** | **0.1606 [0.0668 - 0.2736]** | **0.0326 [0.013 - 0.0593]** | **-** | **34.895** |
| **CIRC** | **M7** | **159.4237** | **5111.5907** | **800.8 [280.975 - 1462.2]** | **474.5 [249.805 - 746.0275]** | **0.1567 [0.0568 - 0.2805]** | **0.1556 [0.0565 - 0.274]** | **0.0315 [0.0105 - 0.0567]** | **-** | **35.3801** |
| HGHT | M1 | 2518.4848 | 26338.5265 | 7525 [223.655 - 19331.75] | 18000 [5160.775 - 35534.25] | 0.2857 [0.0076 - 0.6516] | 0.2081 [0.0071 - 0.6294] | 0.0012 [0 - 0.0031] | 13.6569 | -1.2132 |
| **HGHT** | **M3** | **943.4935** | **37848.1152** | **7360 [381.425 - 15770]** | **3603 [330.4475 - 8654.375]** | **0.1945 [0.0124 - 0.4939]** | **0.2347 [0.0116 - 0.4829]** | **0.0083 [4e-04 - 0.0176]** | **5.9275** | **2.1782** |
| HGHT | M4 | 943.4935 | 37848.1152 | 5088 [191.8975 - 12862.75] | 4134 [856.8975 - 9055.1] | 0.1344 [0.0063 - 0.4298] | 0.1602 [0.006 - 0.4212] | 0.0057 [2e-04 - 0.0144] | 7.1963 | -0.0542 |
| HGHT | M5 | 1356.4464 | 514791.618 | 1643 [73.307 - 6167] | 2635 [300.285 - 6117.075] | 0.0032 [0.0024 - 0.1927] | 0.0523 [0.0024 - 0.1894] | 9e-04 [0 - 0.0033] | 4.0747 | -1.3056 |
| **HGHT** | **M6** | **632.0172** | **30825.812** | **6896 [3179.3 - 11720.5]** | **7158 [4358.95 - 10561.25]** | **0.2237 [0.1043 - 0.372]** | **0.2205 [0.1021 - 0.3663]** | **0.0173 [0.0076 - 0.0299]** | **-** | **74.3619** |
| **HGHT** | **M7** | **632.0172** | **30825.812** | **6660 [2968.825 - 10711.75]** | **7179 [4458.575 - 10412.5]** | **0.2161 [0.1038 - 0.3415]** | **0.2135 [0.1034 - 0.3388]** | **0.0167 [0.0073 - 0.0275]** | **-** | **75.413** |
| RSURF | M1 | 1976.6407 | 351707.3123 | 6022 [756.8225 - 7854.625] | 59300 [26017.5 - 100760] | 0.0171 [0.0022 - 0.0223] | 0.0087 [0.0018 - 0.0214] | 0.0015 [2e-04 - 0.0021] | 0.3481 | -2.0624 |
| RSURF | M3 | 357.1333 | 58445.0405 | 9712 [4384.6 - 11093] | 20620 [9209.775 - 35045.25] | 0.1662 [0.075 - 0.1898] | 0.0463 [0.0231 - 0.0866] | 0.0761 [0.0235 - 0.2209] | 14.2753 | -0.2808 |
| RSURF | M4 | 357.1333 | 58445.0405 | 8800 [3872.05 - 10140.5] | 20360 [10157.75 - 34850.75] | 0.1506 [0.0663 - 0.1735] | 0.042 [0.0192 - 0.077] | 0.069 [0.0198 - 0.2466] | 13.5281 | -0.7364 |
| RSURF | M5 | 1558.1192 | 782897.1412 | 6733 [2716.275 - 7865.325] | 15120 [6236 - 26552] | 0.0086 [0.0035 - 0.01] | 0.0036 [0.0015 - 0.0049] | 0.0028 [0.001 - 0.0036] | 3.3323 | -2.1245 |
| RWDTH | M1 | 2.4974 | 0.1383 | 0.0039 [6e-04 - 0.005] | 0.018 [0.0063 - 0.0345] | 0.0283 [0.0044 - 0.0359] | 0.0059 [0.0011 - 0.0095] | 6e-04 [1e-04 - 8e-04] | -0.5393 | -2.1188 |
| **RWDTH** | **M3** | **2.6695** | **0.6782** | **0.1497 [0.0697 - 0.1749]** | **0.0059 [1e-04 - 0.0212]** | **0.2207 [0.1028 - 0.2579]** | **0.1356 [0.1026 - 0.3313]** | **0.021 [0.01 - 0.0266]** | **-1.8866** | **1.7777** |
| **RWDTH** | **M4** | **2.6695** | **0.6782** | **0.1539 [0.0665 - 0.1799]** | **0.0046 [1e-04 - 0.0196]** | **0.2269 [0.0981 - 0.2653]** | **0.1394 [0.098 - 0.3382]** | **0.0216 [0.0086 - 0.0274]** | **-1.9536** | **1.8914** |
| **RWDTH** | **M5** | **2.5443** | **0.2844** | **0.0741 [0.0518 - 0.0852]** | **0.0016 [1e-04 - 0.0057]** | **0.2606 [0.1822 - 0.2995]** | **0.0677 [0.0694 - 0.138]** | **0.0114 [0.0078 - 0.0137]** | **-2.194** | **2.532** |
| R success | | | | | | | | | | |
| NOFF | M1 | 6.093 | 26.9444 | 2.947 [0.0481 - 21.9028] | 4.198 [0.0148 - 12.0928] | 0.1094 [0.0019 - 0.8008] | 0.1061 [0.0019 - 0.8353] | 0.0794 [0.001 - 0.7408] | 0.7685 | -1.8921 |
| Phenology | | | | | | | | | | |
| **LUs** | **M2** | **3.3609** | **1.5921** | **1.288 [0.0435 - 1.5541]** | **0.1008 [3e-04 - 0.3423]** | **0.809 [0.0273 - 0.9762]** | **0.9154 [0.0319 - 0.9826]** | **0.114 [0.0039 - 0.1399]** | **-** | **5.9259** |
| **LUs** | **M3** | **3.7171** | **1.5781** | **0.9981 [0.4315 - 1.5749]** | **0.0353 [9e-04 - 0.14]** | **0.6325 [0.2734 - 0.998]** | **0.6385 [0.2999 - 0.9267]** | **0.0722 [0.0316 - 0.1184]** | **-1.6954** | **22.6171** |
| **LUs** | **M4** | **3.7171** | **1.5781** | **0.8643 [0.3354 - 1.4832]** | **0.0389 [0.0011 - 0.1531]** | **0.5477 [0.2126 - 0.9399]** | **0.5475 [0.237 - 0.8672]** | **0.0626 [0.0243 - 0.1113]** | **-1.6381** | **15.3716** |
| **LUs** | **M6** | **1.9972** | **1.2233** | **0.4913 [0.4267 - 0.5642]** | **0.1485 [0.0952 - 0.2207]** | **0.4016 [0.3488 - 0.4612]** | **0.4136 [0.367 - 0.4588]** | **0.1232 [0.0945 - 0.1598]** | **-** | **621.9963** |
| **LUs** | **M7** | **1.9972** | **1.2233** | **0.4912 [0.4184 - 0.5621]** | **0.1491 [0.0923 - 0.2207]** | **0.4015 [0.342 - 0.4595]** | **0.4133 [0.3619 - 0.4608]** | **0.1231 [0.0966 - 0.1619]** | **-** | **620.7063** |
| **Lud** | **M2** | **101.9689** | **39.1092** | **32.06 [1.033 - 38.3775]** | **1.6463 [0.0064 - 7.3301]** | **0.8198 [0.0294 - 0.9099]** | **0.9005 [0.0315 - 0.9853]** | **-** | **-** | **2.1757** |
| LS | M2 | 306.3261 | 14.4259 | 1.051 [0.0404 - 1.6353] | 4.2619 [1.6723 - 8.0648] | 0.0729 [0.0069 - 0.2854] | 0.0663 [0.0088 - 0.3961] | - | - | -1.9854 |
| **GSL** | **M2** | **204.0522** | **43.1537** | **32.79 [0.7395 - 40.053]** | **5.3522 [0.0219 - 13.9751]** | **0.7598 [0.0179 - 0.8593]** | **0.7823 [0.0193 - 0.9767]** | **8e-04 [0 - 0.001]** | **-** | **0.6816** |
| **FFLW** | **M1** | **120.7989** | **27.054** | **13.48 [0.2953 - 17.7203]** | **0.0319 [0.0067 - 2.9453]** | **0.4983 [0.0109 - 0.655]** | **0.9713 [0.0223 - 0.9823]** | **-** | **-2.0204** | **2.5067** |
| **MFLW** | **M1** | **117.9556** | **32.2575** | **16.89 [0.3639 - 22.2225]** | **0.2 [0.0085 - 4.3279]** | **0.5236 [0.0113 - 0.6889]** | **0.9717 [0.0199 - 0.9827]** | **-** | **-1.9784** | **4.3772** |
| MAR | M1 | 1.0294 | 2.9028 | 0.6461 [0.0013 - 1.018] | 0.0017 [0 - 0.0967] | 0.2226 [0.0014 - 0.8969] | 0.7126 [0.0014 - 0.9811] | 0.6097 [0.0012 - 1.0633] | -2.0766 | -0.4242 |
| Physiology | | | | | | | | | | |
| C | M2 | 461.5233 | 66.8823 | 49.59 [0.0621 - 73.6105] | - | 0.7415 [9e-04 - 0.9049] | 0.7757 [9e-04 - 0.9865] | 2e-04 [0 - 3e-04] | - | -0.4399 |
| C | M3 | 465.0559 | 255.5694 | 27.4 [0.0605 - 102.8125] | 4.069 [0.002 - 20.8815] | 0.1072 [2e-04 - 0.4023] | 0.1063 [2e-04 - 0.3963] | 1e-04 [0 - 5e-04] | -1.3914 | -1.6093 |
| C | M4 | 465.0559 | 255.5694 | 30.02 [0.1115 - 101.955] | 4.043 [0.004 - 23.6722] | 0.1175 [4e-04 - 0.3935] | 0.1164 [4e-04 - 0.39] | 1e-04 [0 - 5e-04] | -1.3964 | -1.5194 |
| C.N | M2 | 23.6579 | 12.7377 | 0.0068 [0.0016 - 11.3212] | - | 5e-04 [1e-04 - 0.9293] | 5e-04 [1e-04 - 0.9782] | 0 [0 - 0.0205] | - | -2.0013 |
| **C.N** | **M3** | **20.0517** | **5.7817** | **1.447 [0.0206 - 3.3416]** | **0.1601 [0.0031 - 0.5774]** | **0.2503 [0.0038 - 0.5779]** | **0.2606 [0.0038 - 0.5746]** | **0.0036 [1e-04 - 0.0083]** | **-1.1782** | **2.3859** |
| **C.N** | **M4** | **20.0517** | **5.7817** | **1.448 [0.0283 - 3.1997]** | **0.1597 [0.002 - 0.6604]** | **0.2504 [0.0056 - 0.5558]** | **0.2607 [0.0056 - 0.5464]** | **0.0036 [1e-04 - 0.008]** | **-1.1816** | **2.4033** |
| **d13C** | **M2** | **-30.0694** | **1.1119** | **0.971 [0.0092 - 1.162]** | **-** | **0.8733 [0.0088 - 0.9029]** | **0.9784 [0.0088 - 0.9883]** | **0.0011 [0 - 0.0013]** | **-** | **8.3192** |
| **d13C** | **M3** | **-29.7046** | **0.8841** | **0.2998 [0.058 - 0.5783]** | **0.3881 [0.1438 - 0.7227]** | **0.3391 [0.0559 - 0.5607]** | **0.274 [0.0551 - 0.5318]** | **3e-04 [1e-04 - 6e-04]** | **26.1661** | **6.3232** |
| **d13C** | **M4** | **-29.7046** | **0.8841** | **0.2989 [0.0724 - 0.5788]** | **0.4088 [0.1439 - 0.7801]** | **0.3381 [0.0745 - 0.5583]** | **0.2679 [0.0716 - 0.5309]** | **3e-04 [1e-04 - 7e-04]** | **25.7712** | **5.6912** |
| **d15N** | **M2** | **-2.276** | **1.5707** | **1.404 [0.0039 - 1.672]** | **-** | **0.8938 [0.0024 - 0.9052]** | **0.9743 [0.0024 - 0.9877]** | **0.271 [7e-04 - 0.331]** | **-** | **4.7475** |
| **d15N** | **M3** | **-4.1561** | **2.6515** | **0.2026 [0.0165 - 0.5285]** | **1.123 [0.4823 - 1.9801]** | **0.0764 [0.0089 - 0.2954]** | **0.0946 [0.0077 - 0.268]** | **0.0117 [9e-04 - 0.0344]** | **58.8223** | **0.35** |
| d15N | M4 | -4.1561 | 2.6515 | 0.1691 [0.0136 - 0.4852] | 1.14 [0.4696 - 2.0155] | 0.0638 [0.0074 - 0.2671] | 0.0784 [0.0066 - 0.234] | 0.0098 [8e-04 - 0.0315] | 59.5558 | -0.199 |
| N | M2 | 19.8252 | 6.0234 | 0.0343 [9e-04 - 4.9336] | - | 0.0057 [1e-04 - 0.8576] | 0.0057 [1e-04 - 0.902] | 1e-04 [0 - 0.0126] | - | -2.001 |

| | | | | | | | | | | |
|---|---|---|---|---|---|---|---|---|---|---|
| **N** | **M3** | **23.505** | **7.7211** | **2.924 [0.5577 - 5.5019]** | **0.1466 [0.005 - 0.6786]** | **0.3787 [0.082 - 0.707]** | **0.3944 [0.0817 - 0.695]** | **0.0053 [0.001 - 0.01]** | **-1.4443** | **6.4934** |
| **N** | **M4** | **23.505** | **7.7211** | **2.911 [0.6071 - 5.6224]** | **0.1237 [0.0033 - 0.6132]** | **0.377 [0.0866 - 0.7294]** | **0.3931 [0.0861 - 0.7051]** | **0.0053 [0.0011 - 0.0103]** | **-1.6372** | **6.5752** |
| **SLA** | **M2** | **13.7571** | **9.381** | **8.225 [0.2446 - 10.2308]** | **-** | **0.8768 [0.0293 - 0.9048]** | **0.9415 [0.0293 - 0.9875]** | **0.0435 [0.0013 - 0.0546]** | **-** | **2.5715** |
| SLA | M3 | 13.7343 | 11.9319 | 2.168 [0.1142 - 5.305] | 4.953 [1.6634 - 9.5934] | 0.1817 [0.0095 - 0.4322] | 0.1569 [0.0087 - 0.3973] | 0.0115 [6e-04 - 0.0294] | 35.4158 | -0.788 |
| SLA | M4 | 13.7343 | 11.9319 | 2.464 [0.1919 - 5.7007] | 5.015 [1.8376 - 9.4703] | 0.2065 [0.0149 - 0.436] | 0.1772 [0.0137 - 0.4194] | 0.0131 [0.0011 - 0.0314] | 32.2456 | -0.6621 |
| **MLA** | **M2** | **35.7683** | **104.5311** | **68.57 [0.0394 - 112.3025]** | **-** | **0.656 [4e-04 - 0.9018]** | **0.6905 [4e-04 - 0.9836]** | **0.0536 [0 - 0.0877]** | **-** | **0.385** |
| MLA | M3 | 23.2063 | 46.1773 | 0.2818 [0.1432 - 12.8742] | 8.248 [1.7688 - 18.1232] | 0.0061 [0.0031 - 0.2819] | 0.0058 [0.0031 - 0.2668] | 5e-04 [3e-04 - 0.0239] | 9.3583 | -2.0832 |
| MLA | M4 | 23.2063 | 46.1773 | 0.2888 [0.136 - 11.4905] | 8.236 [1.3268 - 17.8665] | 0.0063 [0.0031 - 0.2392] | 0.0059 [0.0031 - 0.2321] | 5e-04 [3e-04 - 0.0208] | 8.6468 | -2.0827 |
| Resilience | | | | | | | | | | |
| REC | M1 | 1.6164 | 0.1241 | 0.0163 [0 - 0.0416] | 0.0046 [0 - 0.0248] | 0.1315 [2e-04 - 0.3353] | 0.7537 [0.0191 - 0.9609] | 0.0062 [0 - 0.0161] | -1.872 | -1.308 |
| REL | M1 | 0.8583 | 0.0515 | 2e-04 [1e-04 - 0.0107] | 0.0137 [4e-04 - 0.031] | 0.0035 [0.0016 - 0.208] | 0.0128 [0.008 - 0.607] | 2e-04 [1e-04 - 0.0157] | 12.9654 | -2.0389 |
| RET | M1 | 0.5792 | 0.0141 | 0 [0 - 0.0027] | 0.002 [0 - 0.0058] | 0.0029 [4e-04 - 0.1906] | 0.0194 [0.0107 - 0.9043] | 1e-04 [0 - 0.0082] | 6.2005 | -2.0336 |
| Structure | | | | | | | | | | |
| **WD** | **M1** | **549.5415** | **756.3841** | **600 [7.7446 - 745.0125]** | **15.18 [0.2725 - 105.0075]** | **0.7932 [0.0114 - 0.8962]** | **0.9393 [0.0113 - 0.9828]** | **0.002 [0 - 0.0025]** | **-1.995** | **3.3434** |
| **WD** | **M3** | **498.8194** | **5043.3787** | **1062 [10.777 - 2639.15]** | **98.55 [0.5989 - 480.21]** | **0.2106 [0.0023 - 0.5159]** | **0.2083 [0.0023 - 0.5154]** | **0.0043 [0 - 0.0107]** | **-1.6473** | **0.0846** |
| WD | M4 | 498.8194 | 5043.3787 | 902.8 [9.3607 - 2512.825] | 137.7 [0.6702 - 529.9375] | 0.179 [0.0019 - 0.4709] | 0.1762 [0.0019 - 0.4672] | 0.0036 [0 - 0.0101] | -1.4534 | -0.5424 |
| **WD** | **M5** | **514.0477** | **4295.2229** | **753.7 [31.3128 - 1537.1]** | **-** | **0.1755 [0.0085 - 0.3961]** | **0.1996 [0.0085 - 0.3944]** | **0.0029 [1e-04 - 0.0058]** | **-** | **3.0998** |
| Leaf morphology | | | | | | | | | | |
| **BS** | **M1** | **6.5928** | **1.5555** | **0.7368 [0.0197 - 0.9625]** | **0.8134 [0.1613 - 1.605]** | **0.4737 [0.0126 - 0.6188]** | **0.47 [0.0133 - 0.756]** | **0.017 [5e-04 - 0.0233]** | **12.2287** | **2.8342** |
| HR | M1 | 1.3284 | 0.4631 | 0.0322 [7e-04 - 0.0436] | 2e-04 [0 - 0.0073] | 0.0695 [0.0015 - 0.0942] | 0.0817 [0.023 - 0.981] | 0.0182 [4e-04 - 0.0251] | -2.0191 | -1.8387 |
| **LDR** | **M1** | **42.949** | **79.3401** | **50.91 [1.0907 - 62.3365]** | **0.0364 [0.0214 - 8.1577]** | **0.6417 [0.0137 - 0.7857]** | **0.9607 [0.0213 - 0.9855]** | **0.0276 [6e-04 - 0.0339]** | **-2.0179** | **2.0453** |
| LL | M1 | 78.4243 | 199.7889 | 76.48 [1.705 - 95.4317] | 0.5553 [0.0315 - 13.663] | 0.3828 [0.0085 - 0.4777] | 0.4482 [0.0232 - 0.9859] | 0.0124 [3e-04 - 0.0155] | -2.0589 | -0.0828 |
| **LW** | **M1** | **24.7889** | **26.5269** | **13.31 [0.3694 - 16.661]** | **0.078 [0.0056 - 2.2466]** | **0.5018 [0.0139 - 0.6281]** | **0.6213 [0.0289 - 0.9859]** | **0.0217 [6e-04 - 0.027]** | **-2.0501** | **1.3548** |
| **LWR** | **M1** | **31.3872** | **8.436** | **5.263 [0.1264 - 6.6193]** | **0.1679 [0.0025 - 1.2042]** | **0.6239 [0.015 - 0.7846]** | **0.9285 [0.0234 - 0.9832]** | **0.0053 [1e-04 - 0.0067]** | **-1.927** | **0.4713** |
| NL | M1 | 8.7356 | 1.7074 | 0.4353 [0.0097 - 0.6261] | 0.1062 [8e-04 - 0.3138] | 0.2549 [0.0057 - 0.3667] | 0.3231 [0.0192 - 0.9655] | 0.0057 [1e-04 - 0.0083] | -1.3837 | -1.5014 |
| NV | M1 | 2.9019 | 2.1461 | 1.032 [0.0256 - 1.32] | 0.2621 [0.0023 - 0.7418] | 0.4809 [0.0119 - 0.6151] | 0.659 [0.0211 - 0.9623] | 0.1225 [0.003 - 0.1835] | 2.8135 | -0.3589 |
| **OB** | **M1** | **57.0648** | **26.9943** | **10.45 [0.2126 - 15.4515]** | **0.023 [0.0055 - 3.1632]** | **0.3871 [0.0079 - 0.5724]** | **0.9737 [0.0231 - 0.9776]** | **0.0032 [1e-04 - 0.0047]** | **-2.0258** | **0.9377** |
| PL | M1 | 4.6418 | 4.5531 | 0.4451 [0.0095 - 0.6128] | 0.0047 [3e-04 - 0.1191] | 0.0978 [0.0021 - 0.1346] | 0.1075 [0.022 - 0.9793] | 0.0207 [4e-04 - 0.0288] | -2.02 | -1.9069 |
| PR | M1 | 5.3415 | 5.0951 | 2.057 [0.0599 - 2.5661] | 0.0196 [9e-04 - 0.3659] | 0.4037 [0.0118 - 0.5036] | 0.481 [0.0268 - 0.9852] | 0.0721 [0.002 - 0.0902] | -2.0326 | -1.2302 |
| PV | M1 | 33.8468 | 274.5848 | 144.9 [3.6387 - 187.6225] | 47.05 [0.4336 - 111.4025] | 0.5277 [0.0133 - 0.6833] | 0.6929 [0.017 - 0.9604] | 0.1265 [0.0033 - 0.1969] | 6.132 | -0.4726 |
| SW | M1 | 13.7697 | 8.478 | 2.14 [0.0437 - 2.843] | 0.0165 [0.0011 - 0.5144] | 0.2524 [0.0052 - 0.3353] | 0.3142 [0.0204 - 0.9833] | 0.0113 [2e-04 - 0.0151] | -2.0632 | -1.1659 |
| **WP** | **M1** | **44.6226** | **81.5738** | **54 [1.2526 - 69.0618]** | **0.0586 [0.021 - 7.5011]** | **0.662 [0.0154 - 0.8466]** | **0.9353 [0.0242 - 0.9855]** | **0.0271 [6e-04 - 0.0351]** | **-2.0166** | **3.2263** |
| Defence | | | | | | | | | | |
| CNFL (exp) | M1 | 0.9685 | 6.9258 | 0.0011 [1e-04 - 0.2613] | 0.0052 [0 - 0.0577] | 0.0016 [1e-04 - 0.6908] | 0.0029 [1e-04 - 0.7173] | 0.0011 [1e-04 - 0.2613] | -1.9374 | -2.0061 |
| CSTG (exp) | M1 | 3.496 | 19.9558 | 2e-04 [0 - 0.0122] | 0 [0 - 0.0018] | 0.012 [2e-04 - 0.8275] | 0.0145 [2e-04 - 0.8701] | 2e-04 [0 - 0.0122] | -1.9972 | -2.0012 |
| **CSTL (exp)** | **M1** | **0.2009** | **1.308** | **0.2337 [0.0032 - 0.2793]** | **0.0105 [1e-04 - 0.0555]** | **0.6977 [0.0135 - 0.7892]** | **0.8293 [0.0134 - 0.8738]** | **0.2337 [0.0032 - 0.2793]** | **-1.3352** | **5.5538** |
| CWSK (exp) | M1 | -1.9029 | 0.0807 | 0.0577 [2e-04 - 0.936] | 0.0013 [0 - 0.1585] | 0.0283 [1e-04 - 0.4838] | 0.03 [1e-04 - 0.5346] | 0.0577 [2e-04 - 0.936] | -2.0114 | -1.9961 |
| EGNL (exp) | M1 | -1.6995 | 0.1494 | 0.102 [3e-04 - 0.7043] | 0.0822 [0 - 0.2724] | 0.0326 [2e-04 - 0.5273] | 0.0787 [2e-04 - 0.5714] | 0.102 [3e-04 - 0.7043] | 0.0327 | -1.9502 |
| ELAC (exp) | M1 | 0.5203 | 0.7528 | 0.0209 [1e-04 - 0.1533] | 0.0119 [0 - 0.0534] | 0.0956 [4e-04 - 0.7563] | 0.1048 [4e-04 - 0.7964] | 0.0209 [1e-04 - 0.1533] | -0.858 | -1.9036 |
| ELTOT (exp) | M1 | 3.8073 | 393.525 | 0.001 [0 - 0.1225] | 0.0102 [0 - 0.0402] | 0.005 [2e-04 - 0.7187] | 0.0058 [2e-04 - 0.7285] | 0.001 [0 - 0.1225] | -0.3366 | -2.0132 |
| GRDN | M1 | 6.5559 | 8.3491 | 4.519 [0.0037 - 8.8821] | 0.0244 [2e-04 - 1.0253] | 0.5413 [5e-04 - 0.894] | 0.5555 [5e-04 - 0.9798] | 0.1051 [1e-04 - 0.208] | -2.0671 | -0.8921 |
| MVL (exp) | M1 | -0.3239 | 0.2823 | 0.0624 [0.0011 - 0.401] | 0.0929 [5e-04 - 0.2548] | 0.0985 [0.0021 - 0.6133] | 0.0937 [0.0021 - 0.6516] | 0.0624 [0.0011 - 0.401] | 2.2135 | -1.9168 |
| PNTL (exp) | M1 | -1.5888 | 0.0262 | 0.0233 [3e-04 - 0.2924] | 0.0535 [0 - 0.1595] | 0.0361 [6e-04 - 0.6607] | 0.0518 [6e-04 - 0.6917] | 0.0233 [3e-04 - 0.2924] | 3.2031 | -1.9722 |
| ROBA (exp) | M1 | 1.8007 | 6.348 | 4e-04 [0 - 0.0851] | 0.0079 [0 - 0.0289] | 0.0027 [2e-04 - 0.7385] | 0.0034 [2e-04 - 0.7804] | 4e-04 [0 - 0.0851] | 0.1862 | -2.0272 |
| ROBB | M1 | 6.3067 | 4.3557 | 1.248 [0.0012 - 4.3232] | 0.0528 [1e-04 - 0.5983] | 0.2865 [3e-04 - 0.881] | 0.292 [3e-04 - 0.9548] | 0.0314 [0 - 0.1084] | -2.0077 | -1.6943 |
| ROBC (exp) | M1 | 1.5316 | 6.2893 | 0.0444 [0 - 0.1954] | 7e-04 [0 - 0.0265] | 0.1827 [2e-04 - 0.7875] | 0.1978 [2e-04 - 0.8566] | 0.0444 [0 - 0.1954] | -2.0222 | -1.7505 |
| ROBD | M1 | 8.4567 | 12.5036 | 0.0498 [0.002 - 9.1923] | 2e-04 [1e-04 - 1.277] | 0.004 [2e-04 - 0.7879] | 0.0042 [2e-04 - 0.8345] | 7e-04 [0 - 0.1294] | -1.9665 | -2.0014 |
| ROBE (exp) | M1 | 1.894 | 3.3626 | 0.0603 [2e-04 - 0.0897] | 1e-04 [0 - 0.0094] | 0.5705 [0.0024 - 0.8595] | 0.7185 [0.0024 - 0.945] | 0.0603 [2e-04 - 0.0897] | -2.0445 | -0.5345 |
| SYRG | M1 | 6.6793 | 12.5141 | 0.0633 [0.0018 - 7.2206] | 0.3356 [1e-04 - 1.961] | 0.0051 [2e-04 - 0.8473] | 0.0071 [2e-04 - 0.8994] | 0.0014 [0 - 0.169] | -1.553 | -2.0067 |
| **TWSK (exp)** | **M1** | **-2.2078** | **0.3077** | **0.9215 [0.0027 - 1.2623]** | **0.0202 [1e-04 - 0.1954]** | **0.4144 [0.0014 - 0.5226]** | **0.4587 [0.0014 - 0.6016]** | **0.9215 [0.0027 - 1.2623]** | **-1.9625** | **0.9468** |
| VNL | M1 | 3.7411 | 3.8353 | 8e-04 [5e-04 - 2.2532] | 0 [0 - 0.2876] | 2e-04 [2e-04 - 0.8456] | 3e-04 [2e-04 - 0.8922] | 1e-04 [0 - 0.1648] | -2.0006 | -2.0013 |
| VSCG | M1 | 22.9715 | 25.0507 | 13.16 [0.0254 - 23.8702] | 0.3441 [8e-04 - 3.8481] | 0.5253 [0.001 - 0.89] | 0.5782 [0.001 - 0.977] | 0.0249 [0 - 0.0455] | -1.954 | -0.4071 |
| **VSCL (exp)** | **M1** | **0.6863** | **3.3081** | **0.2731 [0.0038 - 0.3283]** | **0.0113 [1e-04 - 0.0573]** | **0.6084 [0.0138 - 0.7703]** | **0.7989 [0.0131 - 0.8591]** | **0.2731 [0.0038 - 0.3283]** | **-1.3436** | **2.3264** |
| 2PHL (exp) | M1 | -2.0999 | 0.0179 | 1e-04 [1e-04 - 0.3599] | 0 [0 - 0.0546] | 1e-04 [1e-04 - 0.6675] | 2e-04 [1e-04 - 0.7121] | 1e-04 [1e-04 - 0.3599] | -2.0006 | -2.0013 |

### Appendix 6

### Comparison of heritability estimation when one parent vs. two parents are used to built the pedigree relationships

The parentage analysis performed on the second generation individuals enabled to retrieve both parents for 122 individuals in *Q. petraea* and 193 in *Q. robur*, and at least one parent for 758 individuals in *Q. petraea* and 636 in *Q. robur* (Truffaut et al., 2017). To make the most complete usage of phenotypic data available on the second generation, we presented the analyses performed on all individuals with at least one parent known. However, when using only individuals with both parents known, the number of individuals decreases but the pedigree becomes more complete. We present here the comparison of heritability estimation ($h^2_{calc}$) with two different sets of individuals: (1) all phenotyped individuals with at least one parent known, and (2) all phenotyped individuals with two parents known.

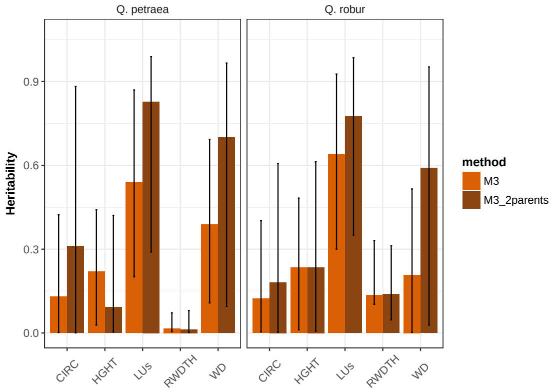


(a) method M3

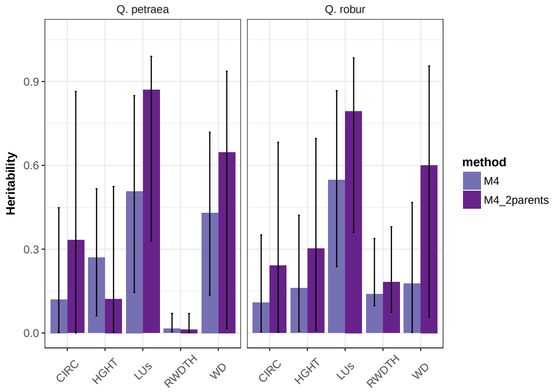


(b) method M4

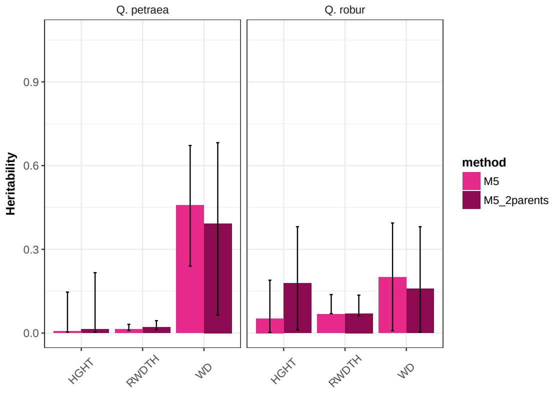


(c) method M5

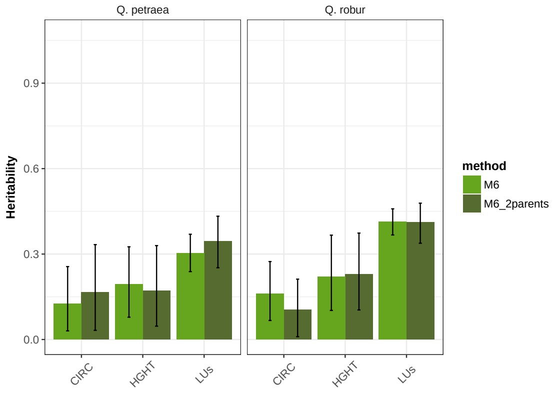


(d) method M6

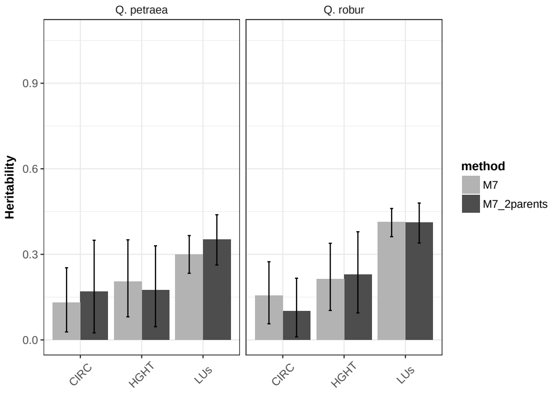


(e) method M7

Figure 6.1: Comparison of heritability estimates ($h^2_{calc}$) when one or two parents are used to built the pedigree. Methods M3 to M7 correspond to the methods described in the main text (Figure 2).

#### Appendix 7

### Genetic correlation among traits

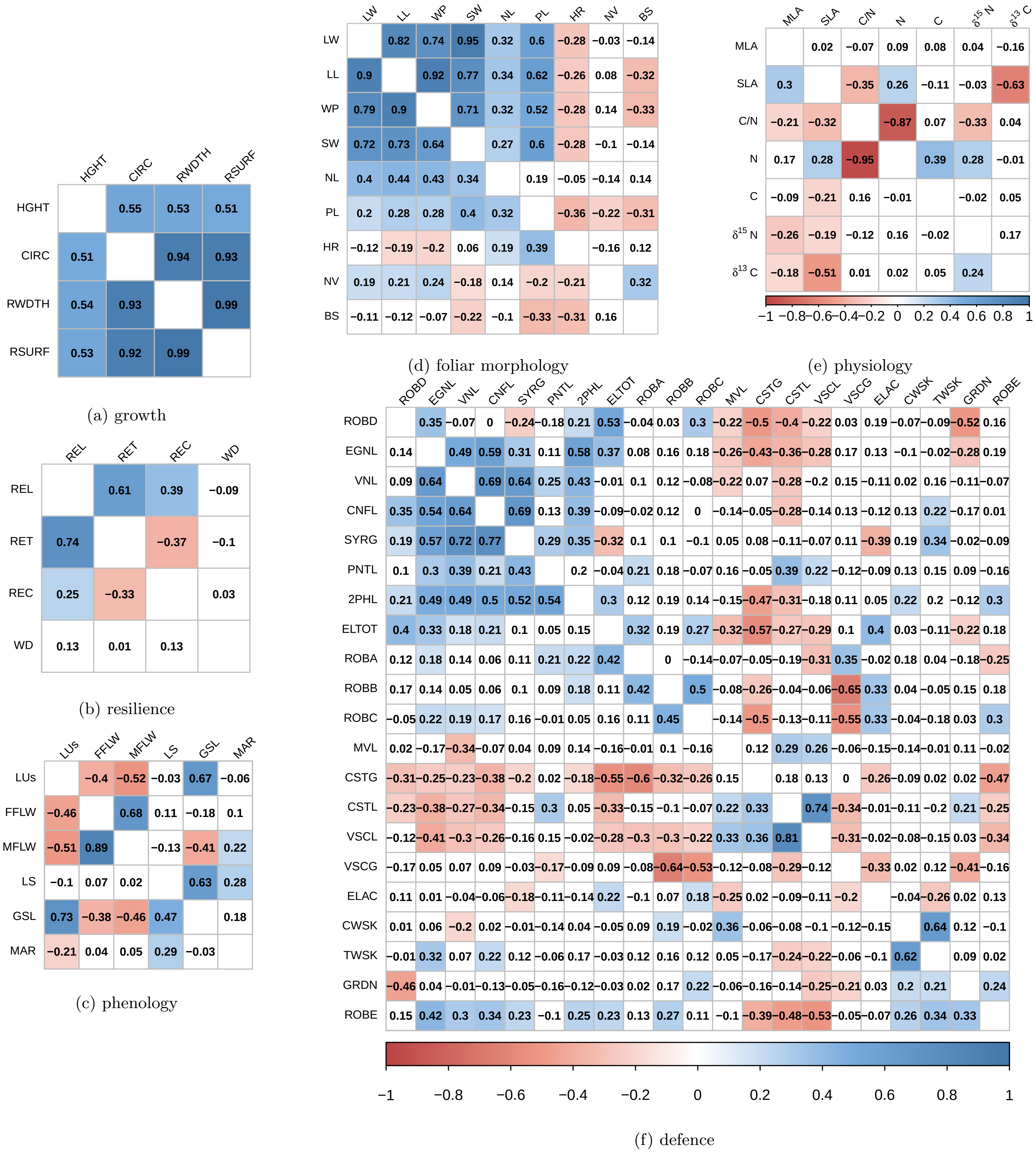


Figure 7.1: Genetic correlations computed from Pearson correlation, for each trait category (a: growth, b: resilience, c: phenology, d: physiology, e: foliar morphology, f: defence). Correlation coefficients for *Q. petraea* are above the matrix diagonals and *Q. robur* are below the diagonal. Colors correspond to the correlation sign (gradient from blue for positive to red for negative correlations). Only correlations significant at a 5% threshold are colored.

### Bibliography


Arend, M., Brem, A., Kuster, T. M., and Gunthardt-Goerg, M. S. (2013). Seasonal photosynthetic responses of European oaks to drought and elevated daytime temperature. *Plant biology*, 15:169–176.

Folke, C., Carpenter, S., Walker, B., Scheffer, M., Elmqvist, T., Gunderson, L., and Holling, C. S. (2004). Regime shifts, resilience, and biodiversity in ecosystem management. *Annual Review of Ecology, Evolution and Systematics*, 35:557–581.

Lagache, L., Klein, E. K., Ducousso, A., and Petit, R. J. (2014). Distinct male reproductive strategies in two closely related oak species. *Molecular Ecology*, 23(17):4331–4343.

Lagache, L., Klein, E. K., Guichoux, E., and Petit, R. J. (2013). Fine-scale environmental control of hybridization in oaks. *Molecular Ecology*, 22(2):423–436.

Lesur, I., Alexandre, H., Boury, C., Chancerel, E., Plomion, C., and Kremer, A. (2018). Development of target sequence capture and estimation of genomic relatedness in a mixed oak stand. *Frontiers in plant science*, page doi: 10.3389.

Lloret, F., Keeling, E. G., and Sala, A. (2011). Components of tree resilience : effects of successive low-growth episodes in old ponderosa pine forests. *Oikos*, (May):1909–1920.

Marshall, T., Slate, J., Kruuk, L. E. B., and Pemberton, J. M. (1998). Statistical confidence for likelihood-based paternity inference in natural populations. *Molecular ecology*, 7:639–655.

Merlin, M., Perot, T., Perret, S., Korboulewsky, N., and Vallet, P. (2015). Effects of stand composition and tree size on resistance and resilience to drought in sessile oak and Scots pine. *Forest Ecology and Management*, 339:22–33.

R Core Team (2016). *R: a language and environment for statistical computing.* R Foundation For Statistical Computing, Vienna, Austria.

Scheffer, M., Carpenter, S., Foley, J. A., Folke, C., and Walker, B. (2001). Catastrophic shifts in ecosystems. *Nature*, 413(October):591–596.

Schweingruber, F., Eckstein, D., Serre-Bachet, F., and Bräker, O. (1998). Identification, presentation and interpretation of event years and pointer years in dendrochronology. *Dendochronologia*, 8:9–38.

Truffaut, L., Chancerel, E., Ducousso, A., Dupouey, J. L., Badeau, V., Ehrenmann, F., and Kremer, A. (2017). Fine-scale species distribution changes in a mixed oak stand over two successive generations. *New Phytologist*, 215:126–139.

Van der Maaten-Theunissen, M., van der Maaten, E., and Bouriaud, O. (2015). pointRes: An R package to analyze pointer years and components of resilience. *Dendochronologia*, 35:34–38.
