## Supplementary material for "*In situ* estimation of genetic variation of functional and ecological traits in *Quercus petraea* and *Q.robur*": script.r

```
#############################################################################################################
# this script presents the analyses used to estimate evolutionary parameters in 
# Alexandre H, Truffaut L, Ducousso A, Louvet JM, Nepveu G, Torres-Ruiz J, Lagane F, Firmat C, Delzon S and Kremer A.
# In situ estimation of genetic variation of functional and ecological traits in Quercus petraea and Q.robur
# 
# It is devided into three parts : 
# 1. the modified version of funtcion breedR.sample.phenotype (original function from package BreedR, Munoz and Sanchez 2018)
# 2. utility functions used to perform heritability estimation
# 3. the final script run for each trait and each species
#
# parts 1 and 2 have to be sourced, and part 3 is modified according to the trait/method/species one wants to analyse
# before starting place all the data files in the working directory and source R files from breedR package; library(breedR); files <- list.files("~/work/R/",full.names=T) ; lapply(as.list(files),source)
#
#############################################################################################################

#############################################################################################################
#############################################################################################################
# 1. breedR.sample.phenotype new version
#############################################################################################################
#############################################################################################################
#'
#' breedR.sample.phenotype(fixed   = c(mu = 10, x = 2),
#'                         random = list(u = list(nlevels = 3,
#'                                                sigma2  = 1,
#'													obs.level = "ind"			)), # ("ind" or "obs")
#'                         genetic = list(model    = 'add_animal',
#'                                        Nparents = c(10, 10),
#'                                        sigma2_a = 2,
#'                                        check.factorial = FALSE),
#'						   generic = list(covar = gen.cov.ok,
#'										  sigma = 1,
#'										  incid = Z.incid),
#'                         spatial = list(model     = 'AR',
#'                                        grid.size = c(5, 5),
#'                                        rho       = c(.2, .8),
#'                                        sigma2_s  = 1),
#'                         residual.variance = 1,
#'                         N = 150,
#'                         obs.matrix = obs.mat)
breedR.sample.phenotype.new <- function(fixed = NULL,
                                    random.ind = NULL,
									random.obs = NULL,
                                    genetic = NULL,
                                    param.spatial = NULL,
									generic = NULL,
                                    residual.variance = 1,
                                    N = NULL,
                                    obs.matrix = NULL) {

 # Number of observations (measurements)
if( !is.null(obs.matrix) ) {
  Nobs <- nrow(obs.matrix)
  Nind <- ncol(obs.matrix)
} else {
  if( is.null(N) ) stop('Please specify sample size (argument N)\n')
  Nobs <- N
  Nind <- N
  obs.matrix <- diag(N)
}

# Number of individuals in the pedigree
if( !is.null(genetic) ) {
  Nobs.ped <- Nobs + sum(genetic$Nparents)
  Nped <- Nind + sum(genetic$Nparents)
} else {
  Nped <- Nind
  Nobs.ped <- Nind
}

components <- list()
phenotype  <- rep(0, Nobs)

# Fixed
if( !is.null(fixed) ) {
  X <- cbind(1,
             matrix(stats::runif(Nobs*(length(fixed) - 1)),
                    nrow = Nobs))
  phenotype <- phenotype + X %*% fixed
  
  components$X <- as.data.frame(X)
  if( is.null(names(fixed)) ) {
    if( length(fixed) == 1 ) names(fixed) <- 'X'
    else names(fixed) <- paste('X', seq_along(fixed)-1, sep = '')
  }
  names(components$X) <- names(fixed)
}

# Random.ind
if( !is.null(random.ind) ) {
  make.random.single <- function(x, N) {
    lev <- sample(x$nlevels, N, replace = TRUE)
    val <- stats::rnorm(x$nlevels, sd = sqrt(x$sigma2))
    return(factor(val[lev], levels = val))
  }
  
   rnd.ind <- as.data.frame(lapply(random.ind,
                                  make.random.single,
                                  N = Nind))
								  
  components$rnd.ind<- obs.matrix%*%apply(rnd.ind,2,as.numeric)
  
  phenotype <- phenotype +apply(obs.matrix%*%apply(rnd.ind,2,as.numeric),1,sum)
}
  
# Random.obs
if( !is.null(random.obs) ) {
  make.random.single <- function(x) {
    lev <- 1:x$nlevels
    val <- stats::rnorm(x$nlevels, sd = sqrt(x$sigma2))
    return(factor(val[lev], levels = val))
  }
  
  rnd.obs <- lapply(random.obs,make.random.single)

  RANDOM <- lapply(rnd.obs,function(y,x) unlist(apply(as.matrix(colSums(obs.matrix)),1,function(x) {
    if (length(y)<x[1]){sample(y,x[1],replace=T)} else {sample(y,x[1],replace=F)}
  })))

  components$rnd.obs <- sapply(RANDOM,function(x) as.numeric(levels(x))[x])
  
  phenotype <- phenotype +apply(components$rnd.obs,1,sum)
}


# Spatial
if( !is.null(param.spatial) ) {
  
  coord <- expand.grid(lapply(param.spatial$grid.size, seq))
  
  # Randomly distribute the observed trees over the grid
  if (Nind > dim(coord)[1]){
    ord <- sample(dim(coord)[1],Nind,replace=T)
  } else ord <- sample(Nind)
  # if( !is.null(genetic) ){
  #   # Make room for founders
  #   # The first sum(genetic$Nparents) will go out of range and result
  #   # in coordinates NA
  #   arrange <- c(dim(coord)[1] + 1:sum(genetic$Nparents), ord)
  # } else
  arrange <- ord
  
  # components <- cbind(components, coord[arrange, ])
  
  if( param.spatial$model == 'AR') {
    components$spatial  <- obs.matrix%*%breedR.sample.AR(param.spatial$grid.size,
                                                         param.spatial$rho,
                                                         param.spatial$sigma2_s)[arrange, 'V1']
    components$coord <- as.data.frame(obs.matrix%*%as.matrix(coord[ord,]))
    names(components$coord)	<- c("lat","long")  
  } else if( param.spatial$model == 'splines') {
    if( !exists('n.knots', param.spatial) ) param.spatial$n.knots <- NULL
    # this one comes in the right order, because it is already
    # multiplied by the incidence matrix
    
    # to be pultiplied by the obs.matrix
    components$spatial  <- obs.matrix%*%breedR.sample.splines(coord,
                                                              param.spatial$n.knots,
                                                              param.spatial$sigma2_s)[arrange, 'V1']
    components$coord <- as.data.frame(obs.matrix%*%as.matrix(coord[ord,]))
    names(components$coord)	<- c("lat","long") 															
  } else stop('Please specify spatial model.')
  
  phenotype <- rowSums(cbind(phenotype, components$spatial), na.rm = TRUE)
}

# Genetic
if( !is.null(genetic) ) {
  
  # Simulate components
  if( is.null(genetic$check.factorial) ) {cf <- TRUE}else {cf <- genetic$check.factorial}
  
  ped <- suppressWarnings(breedR.sample.pedigree(Nind,
                                                 genetic$Nparents,
                                                 check.factorial = cf))
  
  
  components$BV  <- obs.matrix%*%as.matrix(na.omit(cbind(as.data.frame(ped),breedR.sample.BV(ped, genetic$sigma2_a))))
  
  phenotype <- phenotype + as.matrix(components$BV[,"BV"])
  
} 

# Generic
if( !is.null(generic) ) {
  # Simulate components
  gen <- suppressWarnings(breedR.sample.generic(generic$covar,generic$sigma))
  
  components$generic <- as.data.frame(as.matrix(generic$incid%*%gen))
  # Include in the phenotype the corresponding components
  phenotype <- phenotype +
    apply(as.matrix(components$generic),
          1, sum)
} #else generic$sigma = 0

# Residual
components$resid <- stats::rnorm(Nobs, sd = sqrt(residual.variance))
phenotype <- phenotype + components$resid

# Phenotype
components$phenotype <- phenotype

# add parents if genetic components exisits
if (!is.null(genetic)){
  mat <- lapply(components, function(x) as.matrix(x))
  # compo <- lapply(mat, function(x) obs.matrix%*%x)
  cc <- as.data.frame(mat)
  parents <- as.data.frame(matrix(NA,ncol=ncol(cc),nrow=sum(genetic$Nparents)))
  names(parents) <- names(cc)
  components <- rbind(parents,cc)
  components$BV.self[1:sum(genetic$Nparents)] <- 1:sum(genetic$Nparents)
} else {
  mat <- lapply(components, function(x) as.matrix(x))
  # compo <- lapply(mat, function(x) obs.matrix%*%x)
  components <- as.data.frame(mat)
}

# Preferred order of the columns for the first elements
pref.ord <- c('BV.self', 'BV.sire', 'BV.dam', 'row', 'col')
reord <- stats::na.omit(match(pref.ord, names(components)))
if( length(reord) > 0 ) {
  components <- c(components[, reord], components[, -reord])
}

dat <- do.call('data.frame', components)

  return(dat)
}

#' @rdname simulation
#' @param size numeric. A vector of length two with the number of rows and
#'   columns in the field trial
#' @param rho numeric. A vector of length two with the autocorrelation
#'   parameters for the row and column autoregressive processes
#' @param sigma2 numeric. The marginal variance
#' @details \code{breedR.sample.AR} simulates a two-dimensional spatial process
#'   as the kronecker product of first-order autoregressive processes in each
#'   dimension.
breedR.sample.AR <- function(size, rho, sigma2, N = 1){

  # Precision matrices of 1D-AR processes (unscaled)
  Q1d <- mapply(build.AR1d, size, rho)

  # Precision matrix for the AR1(rho_x) x AR1(rho_y) process
  # when locations are stacked following the standards of R:
  # by columns: first vary x and then y
  # (1, 1), (2, 1), ..., (n_x, 1), (1, 2), ..., (n_x, 2), ...
  # Only the lower triangle
  Uinv <- kronecker(Q1d[[2]], Q1d[[1]])

  scaling <- prod(1/(1-rho**2))/sigma2
  Uinv <- Uinv * scaling

  # Simulated samples
  # Note: MASS::mvrnorm only draws samples from the covariance Matrix
  if( !requireNamespace("spam", quietly = TRUE) ) {
    stop("Package spam needed for this simulating AR model. Please install.",
         call. = FALSE)
  }
  ans <- t(spam::rmvnorm.prec(N, Q = Uinv))

  return(as.data.frame(ans))
}

#' @rdname simulation
#' @param coord numeric. A two-column matrix(-like) with spatial coordinates.
#' @param nkn numeric. A vector of length two with the number of (inner) knots
#'   in each dimension
#' @details \code{breedR.sample.splines} simulates a two-dimensional spatial
#'   process as the kronecker product of B-splines processes in each dimension.
breedR.sample.splines <- function(coord, nkn, sigma2, N = 1){

  splines_struct <- breedr_splines(coord, nkn)

  Umat <- get_structure(splines_struct)

  # Simulated samples of effects
  if( !requireNamespace("MASS", quietly = TRUE) ) {
    stop("Package MASS needed for this simulating splines. Please install.",
         call. = FALSE)
  }
  eff <- t(matrix(MASS::mvrnorm(N, mu = rep(0, dim(Umat)[1]), Sigma = Umat*sigma2),
                  nrow = N))

  # Multiply by incidence matrix to get observations at individual level
  ans <- model.matrix(splines_struct) %*% eff

  # cast to matrix (rather than possibly Matrix)
  # and then to data.frame (no direct way)
  return(as.data.frame(as.matrix(ans)))
}

#' @rdname simulation
#' @param ped a pedigree object
#' @param Sigma numeric. The additive genetic variance. Either a variance for a
#'   single additive genetic effect, or a positive-definite matrix with the
#'   covariance structure for a set of correlated genetic effects
#' @details \code{breedR.sample.BV} simulates a set of breeding values (BV)
#'   given a pedigree
breedR.sample.BV <- function(ped, Sigma, N = 1) {

  # Precision matrices are more sparse
  Ainv <- pedigreemm::getAInv(ped)
  Q <- kronecker(solve(Sigma), Ainv)

  # number of genetic effects
  neff <- dim(as.matrix(Sigma))[1]

  # A dim(Ainv)[1] x dim(Sigma)[1] vector with simulated BV
  ans <- matrix(spam::rmvnorm.prec(N, Q = Q),
                nrow = dim(Ainv)[1])
  if(neff == 1) colnames(ans) <- 'BV'
  else colnames(ans) <- paste('BV', 1:neff, sep ='')

  return(ans)
}

breedR.sample.generic <- function(covar, Sigma, N = 1) {

  # Precision matrices are more sparse
  Ainv <- solve(covar)
  Q <- kronecker(solve(Sigma), Ainv)

  # number of generic effects
  neff <- dim(as.matrix(Sigma))[1]

  # A dim(Ainv)[1] x dim(Sigma)[1] vector with simulated BV
  ans <- matrix(spam::rmvnorm.prec(N, Q = Q),
                nrow = dim(Ainv)[1])
  if(neff == 1) colnames(ans) <- 'generic'
  else colnames(ans) <- paste('generic', 1:neff, sep ='')

  return(ans)
}

#' @rdname simulation
#' @param Nobs numeric. Number of individuals to sample
#' @param Nparents numeric. Vector of length two. Number of dams and sires to
#'   randomly mate.
#' @param check.factorial logical. If TRUE (default), it checks whether all the
#'   possible matings had taken place at least once. If not, it stops with an
#'   error.
#' @details \code{breedR.sample.pedigree} simulates a one-generation pedigree
#'   from random mating of independent founders. Note that if
#'   \code{check.factorial} is \code{FALSE}, you can have some founders removed
#'   from the pedigree.
#' @importFrom stats xtabs
breedR.sample.pedigree <- function(Nobs, Nparents, check.factorial = TRUE) {
  stopifnot(length(Nparents) == 2)
  if( is.null(names(Nparents)) ) names(Nparents) <- c('mum', 'dad')
  ped.obs <- data.frame(id = 1:Nobs + sum(Nparents),
                        dad = sample(Nparents['dad'],
                                     size = Nobs,
                                     replace = TRUE),
                        mum = sample(Nparents['mum'],
                                     size = Nobs,
                                     replace = TRUE) + Nparents['dad'])
  # Case half-sibs balanced
  if( Nparents['dad'] == 0 & Nobs %% Nparents['mum'] == 0) {
    ped.obs <- data.frame(id = 1:Nobs + sum(Nparents),
                          dad = 0,
                          mum = sample(rep(1:Nparents['mum'], length = Nobs)))
  }
  fullped <- build_pedigree(1:3, data = ped.obs)

  # checks
  if( check.factorial )
    stopifnot(all(xtabs( ~ dad + mum, data = ped.obs) > 0))
  stopifnot(all(check_pedigree(fullped)))

  return(fullped)
}


#' @rdname simulation
#' @param dim numeric. Dimension of the effect (e.g. n. of traits)
#' @param var numeric matrix. (Co)variance matrix
#' @param Nlevels numeric. Number of individuals values to sample
#' @param labels character vector of labels for each level.
#' @param vname string. A name for the resulting variables
#' @details \code{breedR.sample.ranef} simulates a random effect with a given
#'   variance.
breedR.sample.ranef <-
  function(dim, var, Nlevels, labels = NULL, N = Nlevels, vname = 'X') {
    if(!is.null(dim(var))) stopifnot(identical(dim(var), rep(as.integer(dim), 2)))
    if(!is.null(labels)) stopifnot(all.equal(length(labels), Nlevels))

    ## Simulate Nlevels correlated vectors of dimension dim
    U <- chol(var)
    values <- matrix(stats::rnorm(dim*Nlevels), ncol = dim) %*% U

    ## Sample N observations of the 'factor'
    if (N == Nlevels) {
      idx <- seq_len(Nlevels)
    } else {
      ## either N > Nlevels or N < Nlevels
      idx <- sample(seq_len(Nlevels), N, replace = TRUE)
    }
    ans <- data.frame(values[idx, ])

    ## variable names from variance matrix colnames
    if (!is.null(colnames(var)))
      trait_names <- colnames(var)
    else trait_names <- seq_len(dim)
    varnames <- paste(vname, trait_names, sep = "_")

    if (is.null(labels)) {
      ## variable names from variance matrix colnames
      if (!is.null(colnames(var)))
        labels <- colnames(var)
      else labels <- seq_len(dim)
      names(ans) <- varnames
    } else {
      ## factor labels as an additional variable
      ans <- cbind(factor(labels[idx]), ans)
      names(ans) <- c(vname, varnames)
    }


    return(ans)
  }

  
#############################################################################################################
#############################################################################################################
# 2. Utiliy functions
#############################################################################################################
#############################################################################################################
# function to compute heritability and evolvability 
library(Matrix)
library(synbreed)
library(breedR)
library(QGglmm)

# Source the R files from package breedR


# determine the species assignation of offspring
DeterminSpecies <- function(dat.phen,nb.parents){
  ASSIGNATION <- function(x){
    if(x["generation"]=="parent"){x["assignation.individu"]}else{
      if(x["generation"]=="offspring"){
        if(nb.parents==2){
          if(!is.na(x["assignation.mere"]) & !is.na(x["assignation.pere"]) & x["assignation.mere"]==x["assignation.pere"]){x["assignation.mere"]
          }else{NA}
        }else{
          if(nb.parents==1){
            if(!is.na(x["assignation.mere"]) & !is.na(x["assignation.pere"]) & x["assignation.mere"]==x["assignation.pere"]){x["assignation.mere"]}else{
              if(is.na(x["assignation.mere"]) & !is.na(x["assignation.pere"])){x["assignation.pere"]}else{
                if(is.na(x["assignation.pere"]) & !is.na(x["assignation.mere"])){x["assignation.mere"]}else{NA}
              }
            }
          }
        }
        
      }
    }
  }
  pheno. <- dat.phen
  pheno.$espece <- apply(pheno.,1,ASSIGNATION)
  return(pheno.)
}

#  extract exactly the phenotypic data we need
GetPheno <- function(Pheno,Trait,Generation,Espece,Condition){
  pheno. <- droplevels(Pheno[(Pheno$trait==Trait) & (Pheno$generation%in%strsplit(Generation,",")[[1]]) & (Pheno$espece==Espece) & (Pheno$condition%in%Condition),])
  pheno. <- droplevels(pheno.[!is.na(pheno.$individu_genotype),])
  return(pheno.)
}

# compute genomic relatedness matrix (G)
ComputeG <- function(maf,dat.phen){
  trait <- unique(dat.phen$trait)
  espece <- unique(dat.phen$espece)
  form.maf <- paste("maf",maf,"_newVersion.R",sep="")
  to.find <- paste("*",espece,"*",form.maf,"*",sep="")
  mat.name <- list.files(".",pattern=glob2rx(to.find))
  
  RealizedPD <- nearPD(get(load(mat.name)), keepDiag = T)
  G = matrix(RealizedPD[[1]]@x, nrow = RealizedPD[[1]]@Dim[1])
  attr(G, "dimnames") = RealizedPD[[1]]@Dimnames
  gen.cov <- as(G, "dsCMatrix")
  
  # remplacer les mauvais noms Z
  dimnames(gen.cov)[[1]][grepl("indivZ[0-9][0-9][0-9]",dimnames(gen.cov)[[1]])] <- dimnames(gen.cov)[[2]][grepl("indivZ[0-9][0-9][0-9]",dimnames(gen.cov)[[1]])] <- gsub("indivZ","indiv",dimnames(gen.cov)[[1]][grepl("indivZ[0-9][0-9][0-9]",dimnames(gen.cov)[[1]])])
  dimnames(gen.cov)[[1]][grepl("indivZ[0-9][0-9]",dimnames(gen.cov)[[1]])] <- dimnames(gen.cov)[[2]][grepl("indivZ[0-9][0-9]",dimnames(gen.cov)[[1]])] <- gsub("indivZ","indiv0",dimnames(gen.cov)[[1]][grepl("indivZ[0-9][0-9]",dimnames(gen.cov)[[1]])])
  
  to.keep <- dimnames(gen.cov)[[1]]%in%dat.phen$individu_genotype[dat.phen$trait==trait]
  gen.cov <- gen.cov[to.keep,to.keep]
  
  return(gen.cov)
}

# compute pedigree relatedness matrix (A)
ComputeA <- function(dat.phen){
  ped.desc <- unique(dat.phen[,c("individu_genotype","pere","mere")])
  ped.par <- data.frame(individu_genotype = unique(c(ped.desc$pere,ped.desc$mere)),
                        pere=NA,
                        mere=NA)
  ped.par <- ped.par[!is.na(ped.par$individu_genotype),]
  PED <- rbind(ped.par,ped.desc)
  p <- create.gpData(pedigree=create.pedigree(ID=PED[,1],Par1=PED[,2],Par2=PED[,3]))
  A <- kin(p,ret="add")
  B <- apply(A,c(1,2),as.vector)
  C <- as(B, "dsCMatrix")
  
  return(C)
}

GetRelatednessMatrix <- function(mat.type,maf,dat.phen){
  if(mat.type=="G"){
    r.mat <- ComputeG(maf,dat.phen)
  }
  if(mat.type=="A"){
    r.mat <- ComputeA(dat.phen)
  }
  if(mat.type=="H"){
    G <- ComputeG(maf,dat.phen)
    A <- ComputeA(dat.phen)
    
    G <- G[dimnames(G)[[1]]%in%dimnames(A)[[1]],dimnames(G)[[2]]%in%dimnames(A)[[2]]]
    
    A_11 <- A[which(dimnames(A)[[1]]%in%dimnames(G)[[1]]==F),which(dimnames(A)[[1]]%in%dimnames(G)[[1]]==F)] # apparentement entre individus non genotyp?s
    A_22 <- A[which(dimnames(A)[[1]]%in%dimnames(G)[[1]]),which(dimnames(A)[[1]]%in%dimnames(G)[[1]])] # apparentement entre individus g?notyp?s
    A_12 <- A[which(dimnames(A)[[1]]%in%dimnames(G)[[1]]==F),which(dimnames(A)[[1]]%in%dimnames(G)[[1]])] # apparentement entre individus non g?notyp?s et genotyp?s
    A_21 <- A[which(dimnames(A)[[1]]%in%dimnames(G)[[1]]),which(dimnames(A)[[1]]%in%dimnames(G)[[1]]==F)] # apparentement entre individus g?notyp?s et non genotyp?s
    
    # correction de G
    mdG <- mean(diag(G))
    mdA <- mean(diag(A_22))
    mG <- mean(G)
    mA <- mean(A_22)
    
    alpha <- (mdA*mG-mA*mdG)/(mG-mdG)
    beta <- (mdA-alpha)/mdG
    
    G <- beta*G+alpha
    G <- as.matrix(G)
    
    m1 <- A_11+A_12%*%solve(A_22)%*%(G-A_22)%*%solve(A_22)%*%A_21
    m2 <- A_12%*%solve(A_22)%*%G
    m3 <- G%*%solve(A_22)%*%A_21
    m4 <- G
    r.mat <- rbind(cbind(as.matrix(m1),as.matrix(m2)),cbind(as.matrix(m3),as.matrix(m4)))
  }
  return(r.mat)
}

AnimalModel <- function(dat.phen, repeated.measures, fix.eff, ran.eff, spatial.eff, perform.signif,gen.cov){
  
  gen.cov.ok <- gen.cov[dimnames(gen.cov)[[1]]%in%dat.phen$individu_genotype,dimnames(gen.cov)[[2]]%in%dat.phen$individu_genotype]
  # class(gen.cov.ok) <- "matrix"
  gen.cov.ok <- as.matrix(gen.cov.ok)
  pheno.ok <- dat.phen[dat.phen$individu_genotype%in%dimnames(gen.cov)[[1]],]
  # ordonner les lignes dans le bon ordre
  pheno.ok <- pheno.ok[order(as.character(pheno.ok$individu_genotype)),]
  pheno.ok <- droplevels(pheno.ok)
  gen.cov.ok <- gen.cov.ok[order(dimnames(gen.cov.ok)[[1]]),order(dimnames(gen.cov.ok)[[1]])]
  # v?rifier que tout est dans le bon ordre
  if(is.na(table(!as.character(unique(pheno.ok$individu_genotype))==as.character(dimnames(gen.cov.ok)[[1]]))["FALSE"])){
    stop("error in the order of individuals")
  }
  
  #4. creer la matrice d'incidence pour effet additif genetique -----
  n.x <- length(pheno.ok$individu_genotype)
  n.y <- length(unique(pheno.ok$individu_genotype))
  Z <- matrix(nrow=n.x,ncol=n.y)
  for(column in 1:n.y){
    Z[,column] <- c(rep(0,times=sum(table(pheno.ok$individu_genotype)[order(names(table(pheno.ok$individu_genotype)))][0:(column-1)])),
                    rep(1,times=table(pheno.ok$individu_genotype)[order(names(table(pheno.ok$individu_genotype)))][column]),
                    rep(0,times= n.x - (sum(table(pheno.ok$individu_genotype)[order(names(table(pheno.ok$individu_genotype)))][0:(column-1)])+table(pheno.ok$individu_genotype)[order(names(table(pheno.ok$individu_genotype)))][column]))
    )
  }
  
  Z.incid <- as(cbind(Z),"indMatrix")
  
  if (spatial.eff==T){
    CalcLongLat <- function(x,nb.class){
      CAT <- seq(min(x),max(x),(max(x)-min(x))/nb.class)
      newL <- lapply(x,function (y) {if(y==max(CAT)){max(CAT[CAT<y])}else{max(CAT[CAT<=y])}})
      return(newL)
    }
    pheno.ok$LONG <- unlist(CalcLongLat(pheno.ok$longitude,10))
    pheno.ok$LAT <- unlist(CalcLongLat(pheno.ok$latitude,10))
    
    Spat <- list(model = 'splines',
                 coord = pheno.ok[, c('LONG','LAT')],
                 n.knots = c(6,6))
  }else {Spat <- NULL}
  
  if(unique(pheno.ok$distribution)=="gaussian"){
    Fix <- as.formula(paste("valeur ~",fix.eff))
  }else{
    pheno.ok$logval <- log(pheno.ok$valeur)
    Fix <- as.formula(paste("logval ~",fix.eff))
  }
  
  if (repeated.measures==T){
    Ran <- as.formula(paste("~ ",ran.eff,"+ individu_genotype"))
  }else{Ran <- as.formula(paste("~",ran.eff))}
  
  Gen <- list(gen = list(incidence=Z.incid,
                         covariance=gen.cov.ok))
  
  mod <- remlf90 (fixed= Fix,
                  random= Ran ,
                  spatial = Spat,
                  generic= Gen,
                  data= pheno.ok,
                  method="em")
  
  if (perform.signif==T){
    mod.less.add <- remlf90 (fixed= Fix,
                             random= Ran ,
                             spatial = Spat,
                             generic= NULL,
                             data= pheno.ok,
                             method="em")
    mod.less.spt <- remlf90 (fixed= Fix,
                             random= Ran ,
                             spatial = NULL,
                             generic= Gen,
                             data= pheno.ok,
                             method="em")
    signif.add <- mod.less.add$fit$AIC - mod$fit$AIC
    signif.spt <- mod.less.spt$fit$AIC - mod$fit$AIC
  }else{
    signif.add <- NA
    signif.spt <- NA
  }
  
  res <- list(pheno=pheno.ok,
              kin = gen.cov.ok,
              model=mod,
              signif.add=signif.add,
              signif.spt=signif.spt,
              incid=Z.incid)
  return(res)
}

ComputeHerit <- function(mod,repeated.measures,mat.type,maf,nb.parents){
  m <- data.frame(espece=NA)
  
  m$Va <- mod$model$var["gen",]
  if("spatial"%in%rownames(mod$model$var)){
    m$Vsp <- mod$model$var["spatial",]
  }else{if("bloc"%in%rownames(mod$model$var) & "line"%in%rownames(mod$model$var)){
    m$Vsp <- mod$model$var["bloc",]+mod$model$var["line",]
  }else{if("bloc"%in%rownames(mod$model$var)){
    m$Vsp <- mod$model$var["bloc",]
  }else{m$Vsp <- NA}}}
  
  m$Vr <- mod$model$var["Residual",]
  m$mu <- mean(fitted(mod$model))
  m$pheno_mean <- mean(mod$pheno$valeur)
  m$Vp_obs <- var(mod$pheno$valeur)
  m$h2_calc <- mod$model$var["gen",]/sum(mod$model$var)  
  m$Ia <- mod$model$var["gen",]/(mean(mod$pheno$valeur)^2)
  
  if(unique(mod$pheno$distribution)=="gaussian"){
    m$h2_obs <- mod$model$var["gen",]/var(mod$pheno$valeur)
    m$Va.traitscale <- m$Va
    m$Vp.traitscale <- sum(mod$model$var)
    m$mean.traitscale <- mean(fitted(mod$model))
    m$h2.traitscale <- mod$model$var["gen",]/sum(mod$model$var)
    m$Ia.traitscale <- mod$model$var["gen",]/(mean(fitted(mod$model))^2)
  }else
  {
    custom <- list(inv.link   = function(x) {exp(x)},
                   #Inverse of the log-link, i.e. exponential
                   var.func   = function(x) {0},
                   #No "distribution variance", as we just transformed the data
                   d.inv.link = function(x) {exp(x)})
    #Derivative of the inverse log-link, i.e. exponential
    
    out <- QGparams(mu =mean(fitted(mod$model)), 
                    var.p = sum(mod$model$var),
                    var.a = mod$model$var["gen",],
                    custom.model = custom)
    
    out2 <- QGparams(mu =mean(fitted(mod$model)), 
                     var.p = var(log(mod$pheno$valeur)),
                     var.a = mod$model$var["gen",],
                     custom.model = custom)
    
    m$h2_obs <- unlist(out2["h2.obs"])
    m$Va.traitscale <-unlist(out["var.a.obs"])
    m$Vp.traitscale <- unlist(out["var.obs"])
    m$mean.traitscale <- unlist(out["mean.obs"])
    m$h2.traitscale <- unlist(out["h2.obs"])
    m$Ia.traitscale <- unlist(out["var.a.obs"]/(out["mean.obs"]^2))
  }
  
  if (repeated.measures==T){
    m$h2_sans.intra.ind <- mod$model$var["gen",]/sum(mod$model$var[rownames(mod$model$var)!="Residual"])
    anova1 <- lm(valeur~individu_genotype,data=mod$pheno)
    m$Vp_inter <- var(mod$pheno$valeur)*anova(anova1)["individu_genotype","Sum Sq"]/sum(anova(anova1)[,"Sum Sq"])
    m$h2_obs_inter <- m$Va/m$Vp_inter
  }else{m$h2_sans.intra.ind <- m$h2_calc
  m$Vp_inter <- m$Vp_obs
  m$h2_obs_inter <- m$h2_obs}
  
  m$trait <- unique(mod$pheno$trait)
  m$espece <- unique(mod$pheno$espece)
  m$signif.spatial <- mod$signif.spt
  m$signif.additive <- mod$signif.add
  m$trait.distrib <- unique(mod$pheno$distribution)
  m$condition <- unique(mod$pheno$condition)
  m$generation <- paste(unique(mod$pheno$generation)[1],unique(mod$pheno$generation)[2],sep=" and ")
  m$matrice <- mat.type
  m$maf <- maf
  m$nb.parents <- nb.parents
  
  M <- as.character(m)
  names(M) <- names(m)
  return(M[order(names(M))])
}

ComputeHeritBOOTSTRAP <- function(mod,repeated.measures,mat.type,maf,nb.parents){
  m <- data.frame(espece=NA)
  
  m$Va <- mod$model$var["gen",]
  if("spatial"%in%rownames(mod$model$var)){
    m$Vsp <- mod$model$var["spatial",]
  }else{if("bloc"%in%rownames(mod$model$var) & "line"%in%rownames(mod$model$var)){
    m$Vsp <- mod$model$var["bloc",]+mod$model$var["line",]
  }else{if("bloc"%in%rownames(mod$model$var)){
    m$Vsp <- mod$model$var["bloc",]
  }else{m$Vsp <- NA}}}
  
  m$Vr <- mod$model$var["Residual",]
  m$mu <- mean(fitted(mod$model))
  m$pheno_mean <- mean(mod$pheno$valeur)
  m$Vp_obs <- var(mod$pheno$valeur)
  m$h2_calc <- mod$model$var["gen",]/sum(mod$model$var)  
  m$Ia <- mod$model$var["gen",]/(mean(mod$pheno$valeur)^2)
  
  if(unique(mod$pheno$distribution)=="gaussian"){
    m$h2_obs <- mod$model$var["gen",]/var(mod$pheno$valeur)
    m$Va.traitscale <- m$Va
    m$Vp.traitscale <- sum(mod$model$var)
    m$mean.traitscale <- mean(fitted(mod$model))
    m$h2.traitscale <- mod$model$var["gen",]/sum(mod$model$var)
    m$Ia.traitscale <- mod$model$var["gen",]/(mean(fitted(mod$model))^2)
  }else
  {
    custom <- list(inv.link   = function(x) {exp(x)},
                   #Inverse of the log-link, i.e. exponential
                   var.func   = function(x) {0},
                   #No "distribution variance", as we just transformed the data
                   d.inv.link = function(x) {exp(x)})
    #Derivative of the inverse log-link, i.e. exponential
    
    out <- QGparams(mu =mean(fitted(mod$model)), 
                    var.p = sum(mod$model$var),
                    var.a = mod$model$var["gen",],
                    custom.model = custom)
    
    out2 <- QGparams(mu =mean(fitted(mod$model)), 
                     var.p = var(mod$pheno$valeur),
                     var.a = mod$model$var["gen",],
                     custom.model = custom)
    
    m$h2_obs <- unlist(out2["h2.obs"])
    m$Va.traitscale <-unlist(out["var.a.obs"])
    m$Vp.traitscale <- unlist(out["var.obs"])
    m$mean.traitscale <- unlist(out["mean.obs"])
    m$h2.traitscale <- unlist(out["h2.obs"])
    m$Ia.traitscale <- unlist(out["var.a.obs"]/(out["mean.obs"]^2))
  }
  
  if (repeated.measures==T){
    m$h2_sans.intra.ind <- mod$model$var["gen",]/sum(mod$model$var[rownames(mod$model$var)!="Residual"])
    anova1 <- lm(valeur~individu_genotype,data=mod$pheno)
    m$Vp_inter <- var(mod$pheno$valeur)*anova(anova1)["individu_genotype","Sum Sq"]/sum(anova(anova1)[,"Sum Sq"])
    m$h2_obs_inter <- m$Va/m$Vp_inter
  }else{m$h2_sans.intra.ind <- m$h2_calc
  m$Vp_inter <- m$Vp_obs
  m$h2_obs_inter <- m$h2_obs}
  
  m$trait <- unique(mod$pheno$trait)
  m$espece <- unique(mod$pheno$espece)
  m$signif.spatial <- mod$signif.spt
  m$signif.additive <- mod$signif.add
  m$trait.distrib <- unique(mod$pheno$distribution)
  m$condition <- unique(mod$pheno$condition)
  m$generation <- paste(unique(mod$pheno$generation)[1],unique(mod$pheno$generation)[2],sep=" and ")
  m$matrice <- mat.type
  m$maf <- maf
  m$nb.parents <- nb.parents
  
  M <- as.character(m)
  names(M) <- names(m)
  return(M[order(names(M))])
}


AnimalModelBootstrap <- function(mod,nb.boot,mat.type,maf,nb.parents){
  
  resample_quercus <- function() {
    
    if(unique(mod$pheno$distribution)=="gaussian"){
      Fix <- mean(mod$pheno$valeur)
    }else{Fix <- mean(mod$pheno$logval)}
    
    
    # Ran.obs
    variable.ran.obs <- rownames(mod$model$var)[rownames(mod$model$var)%in%c("annee","date")]
    if(length(variable.ran.obs)>0){
      Ran.obs <- lapply(variable.ran.obs,function(x){list(nlevels=length(unique(mod$pheno[,x])),
                                                          sigma2=mod$model$var[rownames(mod$model$var)==x])})
      names(Ran.obs) <- variable.ran.obs
    }else{Ran.obs <- NULL}
    
    # Ran.ind
    variable.ran.ind <- rownames(mod$model$var)[rownames(mod$model$var)%in%c("bloc","line")]
    if ("line"%in%names(mod$model$ranef) & "bloc"%in%names(mod$model$ranef)){
      Ran.ind <- lapply(variable.ran.ind,function(x){list(nlevels=length(unique(mod$pheno[,x])),
                                                          sigma2=mod$model$var[rownames(mod$model$var)==x])})
      names(Ran.ind) <- variable.ran.ind
    }else{
      if((!"line"%in%names(mod$model$ranef)) & "bloc"%in%names(mod$model$ranef)){
        bloc.nelevels <- length(unique(mod$pheno$bloc))
        bloc.sigma2 <- mod$model$var[rownames(mod$model$var)=="bloc"]
        Ran.ind <- list(bloc=list(nlevels=bloc.nelevels,
                                  sigma2=bloc.sigma2),
                        line=list(nlevels=1,
                                  sigma2=0))
      }else{Ran.ind <- NULL}
    }    
    
    Gen <- list(covar=mod$kin,
                sigma=mod$model$var["gen",],
                incid=mod$incid)
    
    if ("spatial"%in%rownames(mod$model$var)){
      Spat <- list(model     = 'splines',
                   grid.size = c(10, 10),
                   n.knots       = c(6,6),
                   sigma2_s  = mod$model$var["spatial",])
    } else{Spat <- NULL}
    
    Res <- mod$model$var["Residual",]
    
    Nind <- length(unique(mod$pheno$individu_genotype))
    dat <- breedR.sample.phenotype.new(fixed = Fix,
                                       random.ind=Ran.ind,
                                       random.obs=Ran.obs,
                                       genetic=NULL,
                                       generic=Gen,
                                       param.spatial=Spat,
                                       residual.variance=Res,
                                       N=Nind,
                                       obs.matrix = mod$incid)
    
    # rename variables to keep the original names
    names(dat) <- gsub("rnd\\.", "", names(dat))
    names(dat) <- gsub("ind\\.", "", names(dat))
    
    return(dat)
  }
  
  sim_target <- function(dat = resample_quercus()) {
    if ("spatial"%in%rownames(mod$model$var)){
      Spat <- list(model = 'splines',
                   coord = dat[, c('coord.long','coord.lat')],
                   n.knots = c(6,6))
    }else {Spat <- NULL}
    
    if ("individu_genotype"%in%names(mod$model$ranef)){
      colnames(mod$incid) <-1:ncol(mod$incid)
      
      g <- unlist(lapply(colnames(mod$incid),function(x){i=which(colnames(mod$incid)==x)
      a =rep(x,colSums(mod$incid)[i])
      return(a)}))
      
      dat$individu_genotype <- g
    }
    
    if (length(names(mod$model$ranef)[!names(mod$model$ranef)%in%c("spatial","gen")])>0){
      Ran <- as.formula(paste("~ ",paste(names(mod$model$ranef)[!names(mod$model$ranef)%in%c("spatial","gen")],
                                         sep="", collapse="+"),sep="" ))
    }else{Ran <- as.formula("~NULL")}
    
    
    
    Gen <- list(gen = list(incidence=mod$incid,
                           covariance=mod$kin))
    ## Fit the original model
    ## avoiding messages about initial variances spec
    res <- suppressMessages(
      remlf90 (fixed= phenotype ~X,
               random= Ran ,
               spatial = Spat,
               generic= Gen,
               data= dat,
               method="em")
    )
    
    ## Extract point estimates of all variances
    ## and point estimate of heritability
    dat$valeur <- dat$phenotype
    dat$distribution <- unique(mod$pheno$distribution)
    
    dat$trait <- unique(mod$pheno$trait)
    dat$espece <- unique(mod$pheno$espece)
    dat$condition <- unique(mod$pheno$condition)
    dat$generation <- mod$pheno$generation
    
    mod.boot <- list(pheno=dat,
                     kin=mod$kin,
                     model=res,
                     signif.add=NA,
                     signif.spt=NA,
                     incid=mod$incid)
    ans <- ComputeHeritBOOTSTRAP(mod=mod.boot,repeated.measures=("individu_genotype"%in%names(mod$model$ranef)),mat.type,maf,nb.parents)
    
    
    fix.eff <- data.frame(eff_null=rep(0,nrow(mod$pheno)))
    for(fix.variable in names(mod$model$fixed)[names(mod$model$fixed)%in%names(mod$pheno)]){
      if(is.numeric(mod$pheno[,fix.variable])){
        fix.eff[,fix.variable] <- as.numeric(mod$model$fixed[fix.variable][[1]][[1]]["value"])*mod$pheno[,fix.variable]
        assign(paste("eff_",fix.variable,sep=""),as.numeric(mod$model$fixed[fix.variable][[1]][[1]]["value"])*mod$pheno[,fix.variable])
      }else{
        fix.eff[,fix.variable] <- unlist(apply(mod$pheno,1,function(x){mod$model$fixed[fix.variable][[1]][[1]]["value"][x[fix.variable],] }))
        assign(paste("eff_",fix.variable,sep=""),
               unlist(apply(mod$pheno,1,function(x){mod$model$fixed[fix.variable][[1]][[1]]["value"][x[fix.variable],] })))
      }
    }
    eff.global <- unlist(apply(fix.eff,1,sum))
    dat$pheno.global <- dat$phenotype+eff.global
    
    var.pheno1 <- var(dat$phenotype)
    var.pheno2 <- var(dat$pheno.global)
    
    if ("individu_genotype"%in%names(dat)){
      m3 <- lm(phenotype~individu_genotype,data=dat)
      m4 <- lm(pheno.global~individu_genotype,data=dat)
      var.interind3 <- var(dat$phenotype)*anova(m3)["individu_genotype","Sum Sq"]/sum(anova(m3)[,"Sum Sq"])
      var.interind4 <- var(dat$pheno.global)*anova(m4)["individu_genotype","Sum Sq"]/sum(anova(m4)[,"Sum Sq"])
    }else{
      var.interind3 <- var(dat$phenotype)
      var.interind4 <- var(dat$pheno.global)
    }
    
    
    ans["Vp_inter.sansfix"] <- as.character(var.interind3)
    ans["VP_inter.avecfix"] <- as.character(var.interind4)
    ans["h2_inter.sansfix"] <- summary(res)$var["gen",]/var.interind3
    ans["h2_inter.avecfix"] <- summary(res)$var["gen",]/var.interind4
    
    return(ans)
  }
  
  empirical_dist <- as.data.frame(t(replicate(nb.boot, sim_target())))
  
  return(empirical_dist)
  
}   

# Assemble results
Herit <- function(Pheno,Trait,Generation,Condition,Espece, fix.eff, ran.eff, spatial.eff, repeated.measures, mat.type,maf=NA, perform.signif,peform.bootstrap=F,nb.boot,nb.parents){
  NewPheno <- GetPheno(Pheno,Trait,Generation,Espece,Condition)
  temp <- as.data.frame(NewPheno[,names(NewPheno)%in%strsplit(fix.eff,"[:+:]")[[1]]])
  rownames(temp) <- rownames(NewPheno)
  temp <- na.omit(temp)
  NewPheno <- NewPheno[rownames(temp),]
  if (mat.type=="H" & Generation=="offspring"){
    GEN.temp <- "parent,offspring"
    parents <- na.omit(unique(c(NewPheno$mere,NewPheno$pere)))
    parents.pheno <- data.frame(matrix(ncol=ncol(NewPheno),nrow=length(parents)))
    names(parents.pheno) <- names(NewPheno)
    parents.pheno$individu_genotype <- parents
    parents.pheno$espece <- unique(NewPheno$espece)
    parents.pheno$trait <- unique(NewPheno$trait)
    New.pheno.withpar <- rbind(parents.pheno,NewPheno)
    R.Matrix <- GetRelatednessMatrix(mat.type,maf,dat.phen=New.pheno.withpar)
  }else{R.Matrix <- GetRelatednessMatrix(mat.type,maf,dat.phen=NewPheno)}
  
  A.Mod <- AnimalModel(dat.phen=NewPheno, repeated.measures, fix.eff, ran.eff, spatial.eff, perform.signif,gen.cov=R.Matrix)
  
  H2 <- ComputeHerit(mod=A.Mod,repeated.measures,mat.type,maf,nb.parents)
  
  if(peform.bootstrap==T){
    H2.boot <- AnimalModelBootstrap(mod=A.Mod,nb.boot,mat.type,maf,nb.parents)
  }else{H2.boot <- NA}
  
  return(list(estimates=H2,
              bootstrap=H2.boot))
}

BreedingValues <- function(Pheno,Trait,Generation,Condition,Espece, fix.eff, ran.eff, spatial.eff, repeated.measures, mat.type,maf=NA,perform.signif){
  NewPheno <- GetPheno(Pheno,Trait,Generation,Espece,Condition)
  temp <- as.data.frame(NewPheno[,names(NewPheno)%in%strsplit(fix.eff,"[:+:]")[[1]]])
  rownames(temp) <- rownames(NewPheno)
  temp <- na.omit(temp)
  NewPheno <- NewPheno[rownames(temp),]
  if (mat.type=="H" & Generation=="offspring"){
    GEN.temp <- "parent,offspring"
    parents <- na.omit(unique(c(NewPheno$mere,NewPheno$pere)))
    parents.pheno <- data.frame(matrix(ncol=ncol(NewPheno),nrow=length(parents)))
    names(parents.pheno) <- names(NewPheno)
    parents.pheno$individu_genotype <- parents
    parents.pheno$espece <- unique(NewPheno$espece)
    parents.pheno$trait <- unique(NewPheno$trait)
    New.pheno.withpar <- rbind(parents.pheno,NewPheno)
    R.Matrix <- GetRelatednessMatrix(mat.type,maf,dat.phen=New.pheno.withpar)
  }else{R.Matrix <- GetRelatednessMatrix(mat.type,maf,dat.phen=NewPheno)}
  
  mod <- AnimalModel(dat.phen=NewPheno, repeated.measures, fix.eff, ran.eff, spatial.eff, perform.signif,gen.cov=R.Matrix)
  
  res <- cbind(dimnames(mod$kin)[[1]],mod$model$ranef$gen[[1]])
  names(res) <- c("individu",paste("BV_",Trait,sep=""),paste("s.e_",Trait,sep=""))
  
  return(res)
}


#############################################################################################################
#############################################################################################################
# 3. trait analysis
#############################################################################################################
#############################################################################################################
the.trait <- "HGHT" # trait Acronym
the.method <- "M1" # method "M1" or "M2" or "M3" or "M4" or "M5" or "M6" or "M7"
the.maf <- 5 # MAF threshold used to compute genomic relatedness
the.par <- 1 # number of parents : 1 or 2 (if error message in spatial effect, change line 581 and 582 into     pheno.ok$LONG <- unlist(CalcLongLat(pheno.ok$longitude,8)) /   pheno.ok$LAT <- unlist(CalcLongLat(pheno.ok$latitude,8))
do.boot <- TRUE # whether to perform bootstrap or not TRUE or FALSE
the.boot <- 1 # number of simulation bootstrap to do (each simulation can be very long especially for traits measured multiple times per individual e.g RWDTH, RSURF)
the.species <- "petraea" # "petraea" or "robur"

# load phenotypic data
pheno <- read.table("phenotypes.txt",h=T,sep="\t",stringsAsFactors = F)
pheno$espece <- pheno$assignation.individu
pheno1 <- DeterminSpecies(dat.phen=pheno,nb.parents=the.par)

# load animal models despcription (this file does not contain all possible models)
list.mod <- read.table("models.txt",h=T,sep="\t",stringsAsFactors = F)

the.generation <- unique(list.mod$generation[list.mod$trait_abbrev==the.trait & list.mod$method==the.method])
the.condition <- unique(list.mod$condition[list.mod$trait_abbrev==the.trait & list.mod$method==the.method])
the.fixeff <- unique(list.mod$fix_eff[list.mod$trait_abbrev==the.trait & list.mod$method==the.method])
the.raneff <- unique(list.mod$random_eff[list.mod$trait_abbrev==the.trait & list.mod$method==the.method])
the.spatialeff <- unique(list.mod$spatial_effect[list.mod$trait_abbrev==the.trait & list.mod$method==the.method])
the.repeat <- unique(list.mod$repeated_measures[list.mod$trait_abbrev==the.trait & list.mod$method==the.method])
the.mat <- unique(list.mod$matrice[list.mod$trait_abbrev==the.trait & list.mod$method==the.method])

pheno2 <- pheno1[pheno1$trait.abb==the.trait,]
if (length(unique(pheno2$trait))>1){
  the.trait.name <- unique(pheno2$trait[pheno2$generation%in%strsplit(the.generation,",")[[1]] & pheno2$condition==the.condition])
}else{the.trait.name <- unique(pheno2$trait)}


res.herit <- Herit(Pheno=pheno1,Trait=the.trait.name,Generation=the.generation,Condition=the.condition,
                           Espece=the.species, fix.eff=the.fixeff , ran.eff=the.raneff, spatial.eff=the.spatialeff,
                           repeated.measures=the.repeat, mat.type=the.mat,maf=the.maf, perform.signif=T,
                           peform.bootstrap=do.boot,nb.boot=the.boot,nb.parents=the.par)

res.bv <- BreedingValues(Pheno=pheno1,Trait=the.trait.name,Generation=the.generation,Condition=the.condition,
                                 Espece=the.species, fix.eff=the.fixeff , ran.eff=the.raneff, spatial.eff=the.spatialeff,
                                 repeated.measures=the.repeat, mat.type=the.mat,maf=the.maf, perform.signif=F)

  
```
